# *nHOTAIRM1* scaffolds ANXA11-dependent assemblies to drive axonal mRNA localization in human motor neurons

**DOI:** 10.64898/2026.09.16.752050

**Authors:** Paolo Tollis, Tiziana Santini, Adriano Setti, Walter Visciglio, Jessica Rea, Davide Mariani, Alessandro Belvedere, Monica Ballarino, Irene Bozzoni, Pietro Laneve

## Abstract

Long noncoding RNAs (lncRNAs) are well recognized as regulators of neuronal development and function, yet their integration into canonical axonal transport pathways, an essential component of spatial gene regulation in highly polarized cells, remains incompletely defined. It is still unclear to what extent individual lncRNAs confer cargo selectivity and couple discrete mRNA cohorts to established transport machineries. Here we identify a cytoplasmic role for the neuronal isoform of *HOTAIRM1* (*nHOTAIRM1*) in shaping the axonal RNA landscape of human iPSC-derived spinal motor neurons (spMNs). We show that *nHOTAIRM1* associates with Annexin A11 (ANXA11), a factor linked to lysosome-coupled movement of RNA granules and engages a defined subset of MN-relevant mRNAs. Transcriptome-wide and targeted interaction assays, together with computational mapping and steric competition, support direct RNA-RNA pairing between *nHOTAIRM1* and mRNAs involved in cytoskeletal organization and synaptic or vesicular functions, while spatial analyses place these RNA pairs in close proximity within the soma and neurites. Loss-of-function experiments further show that *nHOTAIRM1* depletion reduces the association of these mRNAs with ANXA11-positive complexes and diminishes their enrichment in neuronal projections, an effect that is recapitulated by ANXA11 depletion. Collectively, these findings support a model in which *nHOTAIRM1* contributes to the selective recruitment of cargo mRNAs into ANXA11-associated transport assemblies, thereby promoting their localization within distal neurites. More broadly, our findings provide mechanistic insight into how a lncRNA can regulate mRNA sorting and compartment-specific RNA trafficking in human spMNs.

## Introduction

Motor neurons (MNs) are highly polarized cells whose axonal projections can extend over long distances, imposing stringent demands on intracellular transport to maintain morphology, compartmentalized functions, and survival.^1^ A central component of this intracellular logistics system is the active transport of mRNAs to distal compartments, where local translation sustains synaptic function and structural integrity. Axonal mRNA transport and anchoring are orchestrated by RNA-binding proteins (RBPs) that recognize cis-acting elements within target transcripts and assemble them into transport-competent ribonucleoprotein (RNP) granules. These RNP complexes are conveyed along the cytoskeleton by molecular motors, such as kinesins and dyneins, which ensure the precise spatiotemporal delivery of transcripts to their sites of function.^2^ Within these granules, mRNAs undergo compartment-specific control of stability, translation, isoform usage and decay, thereby shaping the local transcriptome and proteome in response to developmental and activity-dependent cues.^3^

Prototypical RBPs such as HuD, FMRP, IGF2BP1 and Staufen illustrate this principle by controlling the localization and fate of growth-associated or synaptic mRNAs, and their dysfunction is tightly linked to axonal and synaptic pathology.^4,5^

While prototypical RBPs have been extensively characterized, recent studies reveal additional layers of complexity in RNP granule assembly, including the emerging role of lncRNAs.

These molecules can act as architectural scaffolds^6–8^ or regulatory adaptors within neuronal RNPs, influencing RNA cargo selection, transport, and local regulation, yet how such lncRNA-dependent mechanisms operate in MNs remains largely unexplored.^9^ This role aligns well with the structural properties of lncRNAs: they are long, highly structured molecules with multiple binding surfaces for RBP and RNA, whose modular domains and relaxed coding constraints allow them to encode localization motifs and multivalent interaction sites, making them ideal organizers of RNP granules and RNA transport routes in neurons. Evidence for lncRNA involvement in subcellular RNA localization is accumulating across neuronal subtypes. In embryonic motor axons, *MALAT1*, *RMST*, *MEG3* and *MIAT* associate with RBPs and motor machineries, consistent with roles in microtubule-based RNA transport.^10^ In hippocampal neurons, ADEPTR influences the localization of actin-related transcripts in an activity-dependent manner^11^ while the synaptically localized *SLAMR* establishes a feedback loop with the motor KIF5C.^12,13^ Finally, in dorsal root ganglion neurons, *ALAE* influences trafficking of synaptic mRNAs during circuit maturation,^14^ while *Dubr* promotes axon elongation via YTHDF1/3-linked translational control.^15^ These examples, while still fragmentary, support the idea of a “lncRNA cargo code” that helps tune synaptic transcriptomes and proteomes.^12^ However, existing studies do not directly demonstrate how a single lncRNA can mechanistically couple a canonical axonal transport factor to the selective recruitment of specific mRNAs in MNs. Consequently, a unified framework explaining how lncRNAs may instruct RNA cargo specificity within axonal transport granules warrants further investigation.

In this study, we show that the lncRNA *nHOTAIRM1* addresses this question by coupling a canonical axonal transport factor to a defined set of mRNA cargoes in human spMNs.

We previously introduced *nHOTAIRM1* as a nuclear regulator of the proneural master gene *NEUROGENIN2* at the transcriptional level, thereby influencing neuronal lineage commitment, ^16^ and more recently, we described the role of *nHOTAIRM1* in regulating MN cell fate, architecture and activity.^17^ Here, we extend the functional repertoire of *nHOTAIRM1*, showing that, in human spMNs, it physically associates with ANXA11 and specific axon- and synapse-related mRNAs. *nHOTAIRM1* forms RNA-RNA duplexes with these transcripts, colocalizes with them in distal processes, and contributes to their incorporation into ANXA11-positive RNP granules and to their enrichment within neurites. The interaction with ANXA11 is mechanistically relevant since this Ca²⁺-dependent protein was described to couple RNA granules to motile lysosomes in axons.^18^ Consistently, ANXA11 knockdown significantly reduces the neuritic localization of *TPM1*, *YKT6* and *SPTBN1* mRNAs, supporting the functional relevance of the *nHOTAIRM1*-ANXA11 axis in their transport.

These findings uncover *nHOTAIRM1* as a molecular adaptor that couples a transport factor to a specific cohort of MN mRNA cargoes. More broadly, they provide a mechanistic example of how lncRNAs can directly instruct axonal mRNA cargo selection and delivery through RNA-RNA interactions, adding a noncoding regulatory layer to the logic of neuronal RNA trafficking and MN physiology.

## Results

### *nHOTAIRM1* associates with ANXA11 in spMNs

We previously characterized *nHOTAIRM1*, a neuronal isoform of *HOTAIRM1*^16^ enriched in the spinal cord, in a human iPSC-based model of spMN differentiation. To investigate its function, we generated two independent homozygous CRISPR/Cas9-edited iPSC clones in which insertion of a premature polyadenylation signal within the 5′ region of *HOTAIRM1* exon 1 induced early transcriptional termination. Analysis of these lines established *nHOTAIRM1* as a pro-MN regulator at the MN-interneuron fate decision and showed that its loss impairs neurite branching and the expression of genes involved in synaptic connectivity and neurotransmission^17^. The loss-of-function analyses described below were performed using KO#2, one of these previously validated clones.

Notably, our previous work also showed that *nHOTAIRM1* localizes not only in the soma but also along neurites and in distal axonal segments of differentiated spMNs.^17^ Here, we dissected this spatial distribution using hybridization chain reaction (HCR) RNA-FISH imaging technology, which provides greater target RNA detection sensitivity and spatial resolution than the previously employed biotin-labeled RNA-FISH. This analysis confirmed the presence of discrete *nHOTAIRM1* signals throughout the soma and extending into neuronal projections, including distal segments (Supplementary Fig. 1A).

To investigate whether this compartmentalized localization was accompanied by specific protein interactions, we re-analyzed the *nHOTAIRM1* protein interactome previously obtained via RNA antisense purification coupled with mass spectrometry (RAP-MS) in spMNs.^16^ Among the proteins enriched with *nHOTAIRM1*, ANXA11 emerged as a candidate interactor of particular interest.

ANXA11 is a Ca²⁺l1ldependent phospholipidl1lbinding protein involved in calcium signaling and membrane dynamics, with established roles in vesicle trafficking and membranel1lassociated processes, including lysosomel1lassociated RNA granule transport.^18^ Recent structural and functional studies indicate that ANXA11 harbors regions with reported RNAl1lbinding properties, located in its C-terminal core domain, and acts as a molecular tether that couples RNA granules to motile lysosomes in neuronal axons, thereby coordinating their long-range trafficking.^18,19^

To spatially contextualize and further validate this candidate RNA-protein interaction, we first examined ANXA11 distribution in spMNs by immunofluorescence (Supplementary Fig. 1B). This analysis revealed a punctate pattern in both the soma and neurites, qualitatively resembling the subcellular distribution of *nHOTAIRM1*.

To obtain biochemical evidence of the *nHOTAIRM1*-ANXA11 association, we performed ANXA11 crosslinking immunoprecipitation (CLIP) assay in spMNs (Fig. 1A, left panel). The results confirmed that *nHOTAIRM1* was significantly enriched in ANXA11-bound RNA fractions supporting their association in a physiological neuronal context (Fig. 1A, right panel).

**Figure 1.**
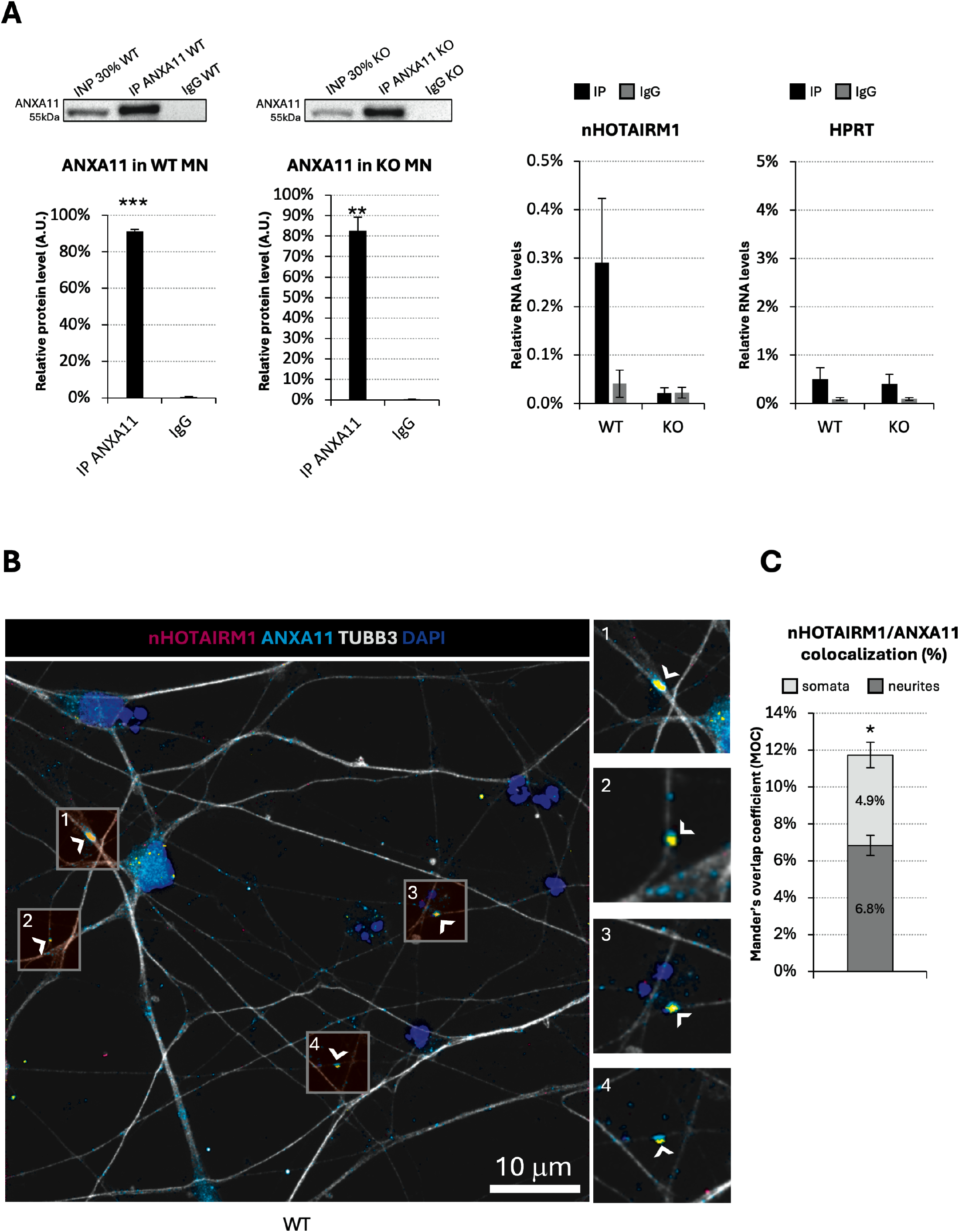
*nHOTAIRM1* and ANXA11 interaction in iPSC-derived spMNs. (A) CLIP assay for ANXA11 in whole cell extracts of iPSC-derived spMNs. Left panels: immunoblot analysis of ANXA11 immunoprecipitated (IP) protein fraction and IgG negative control over input (Inp) in WT and *nHOTAIRM1* KO spMNs. Immunoblots shown in the figure are representative of three independent experiments. Blots were cropped for clarity. Full-length uncropped blots are provided in the Supplemental Material. Right panels: qRT-PCR analysis of *nHOTAIRM1* and *HPRT* RNA enrichment over input, in IP and IgG fractions. Data are expressed as percentage of input. *HPRT* was used as a negative control because i) it is not expected to specifically associate with *nHOTAIRM1*-ANXA11 assemblies, ii) it has no established role in axonal RNA transport, and iii) it is not enriched in the *nHOTAIRM1* RNA interactome. *N*□=□3. (B) Representative RNA-FISH for *nHOTAIRM1* (magenta), IF for TUBB3 (gray), ANXA11 (cyan) and DAPI (blue) in WT spMNs at day 12 of differentiation. Scale bars: 10 µm. (C) Histogram showing the Mander’s overlap coefficient (MOC) between *nHOTAIRM1* and ANXA11 in somata and neurites of WT spMNs. MOC values quantify the fraction of *nHOTAIRM1* signal overlapping ANXA11. ≥50 cells, derived from at least three independent biological replicates, were analyzed. Mean values ± SEM are indicated. Statistical significance was determined by unpaired two-tailed Student’s t test using biological replicates as independent units. *P < 0.05, **P < 0.01, ***P < 0.001.

We then combined RNA-FISH for *nHOTAIRM1* and immunofluorescence for ANXA11 to assess their spatial colocalization. Quantitative imaging analysis showed that approximately 12% of *nHOTAIRM1 foci* colocalized with ANXA11-positive signals (Fig. 1B,C). Notably, almost 5% of these events occurred within the soma and almost 7% in neuronal projections (Fig. 1C), where ANXA11-mediated RNA granule trafficking occurs.^18^ Together, these results indicate that a defined sub-population of *nHOTAIRM1* localizes to neurites where it physically associates with ANXA11, suggesting the possibility that *nHOTAIRM1* contributes to ANXA11-mediated control of RNA trafficking and dynamics in spMN processes.

### The *nHOTAIRM1*-ANXA11 association involves discrete predicted interaction regions

To gain further mechanistic insight into the *nHOTAIRM1*-ANXA11 association, we used *cat*RAPID omics v2.1^20^ to predict the potential protein and RNA interaction determinants (Supplementary Fig. 2A). The analysis of interaction propensity revealed that the putative binding surface of ANXA11 was not distributed across the full-length protein but was largely confined to its C-terminal half (Supplementary Fig. 2A).

Domain annotation using the InterPro database^21^ revealed that essentially all high-propensity fragment pairs mapped within the first three (I,II and III) of the four annexin repeats of the ANXA11 core (Pfam PF00191; residues 204-269, 276-341, 359-425 and 435-500), whereas the extended N-terminal region (approximately residues 1-190) was uniformly predicted as non-interacting (Supplementary Fig. 2A).

The N-terminal region also displayed low AlphaFold confidence (pLDDT; Supplementary Fig. 2A) and was predicted to be intrinsically disordered by the MobiDB-lite consensus (Supplementary Fig. 2A), suggesting a potential role in ANXA11 phase separation and RNA granule formation.^22^ Overall, these features distinguish a flexible, intrinsically disordered N-terminal region from the structured annexin-repeat core, where the predicted *nHOTAIRM1*-interacting surfaces are localized.

On the RNA side, the predicted contacts were similarly non-uniform: rather than extending across the entire *nHOTAIRM1* sequence, they mapped to two discrete regions in the 5’ half of the neuronal isoform, representing the candidate interaction regions (predicted binding domain, BD) of the lncRNA: nucleotides from 76 to 127 (*nHOTAIRM1*-ANXA11 BD1) and nucleotides from 193 to 252 (*nHOTAIRM1*-ANXA11 BD2).

In parallel with the computational inspections, we sought reciprocal, RNA-centric support for the *HOTAIRM1*-ANXA11 association, orthogonal to previous ANXA11 CLIP assays. Native *nHOTAIRM1* RNA pull-down assays were performed in iPSC-derived spMNs at day 12 of differentiation, followed by ANXA11 immunodetection. *nHOTAIRM1* was captured using 18 biotinylated 20-nt antisense DNA probes spanning the entire transcript. As shown in Supplementary Fig. 2B, ANXA11 protein was clearly enriched in the *nHOTAIRM1* RNA pull-down fraction compared with the negative control.

Together, these analyses provide complementary information: computational profiling identifies candidate regions that may contribute to the *nHOTAIRM1*-ANXA11 interaction, whereas RNA pull-down strengthens the biochemical evidence for their association in spMNs.

### *nHOTAIRM1* physically interacts with MN-relevant mRNAs

Given that, in spMNs, *nHOTAIRM1* physically associates with ANXA11, a key mediator of lysosome-associated RNA-granule axonal transport, and that coding-noncoding RNA crosstalk frequently involve RNA-RNA interactions,^23–25^ we performed native *nHOTAIRM1* RNA pull-down assays followed by RNA-seq to identify RNA partners potentially involved in this transport axis (Fig. 2A,B). To increase the stringency of the analysis, we used two independent sets of biotinylated DNA probes, comprising alternating antisense oligonucleotides (even and odd) distributed along the entire sequence of *nHOTAIRM1*. Analysis of three biological replicates showed homogeneous clustering of the samples (Supplementary Fig. 3A) and robust enrichment of *nHOTAIRM1* in both even and odd RNA pull-down fractions compared with input and the lacZ control (Supplementary Fig. 3B). Among 14,136 genes detected (mean input FPKM > 1), we identified 27 high-confidence *nHOTAIRM1* RNA interactors consistently enriched in both even and odd RNA pull-down sets (log2 pull-down/input > 0.59; FDR < 0.05; Supplementary Fig. 3C, left) and specifically enriched in the lncRNA pull-down relative to the LacZ control (for further details, see Methods, Supplementary Fig. 3C, right, and Supplementary Fig. 3D). We validated the RNA-seq results by qRT-PCR on a subset of ten interactors selected across the range of significantly enriched transcripts (log2FC > 0.59; FDR < 0.05), including top-, intermediate-, and lower-ranking candidates, to avoid bias toward the most strongly enriched hits (Supplementary Fig. 4A).

**Figure 2.**
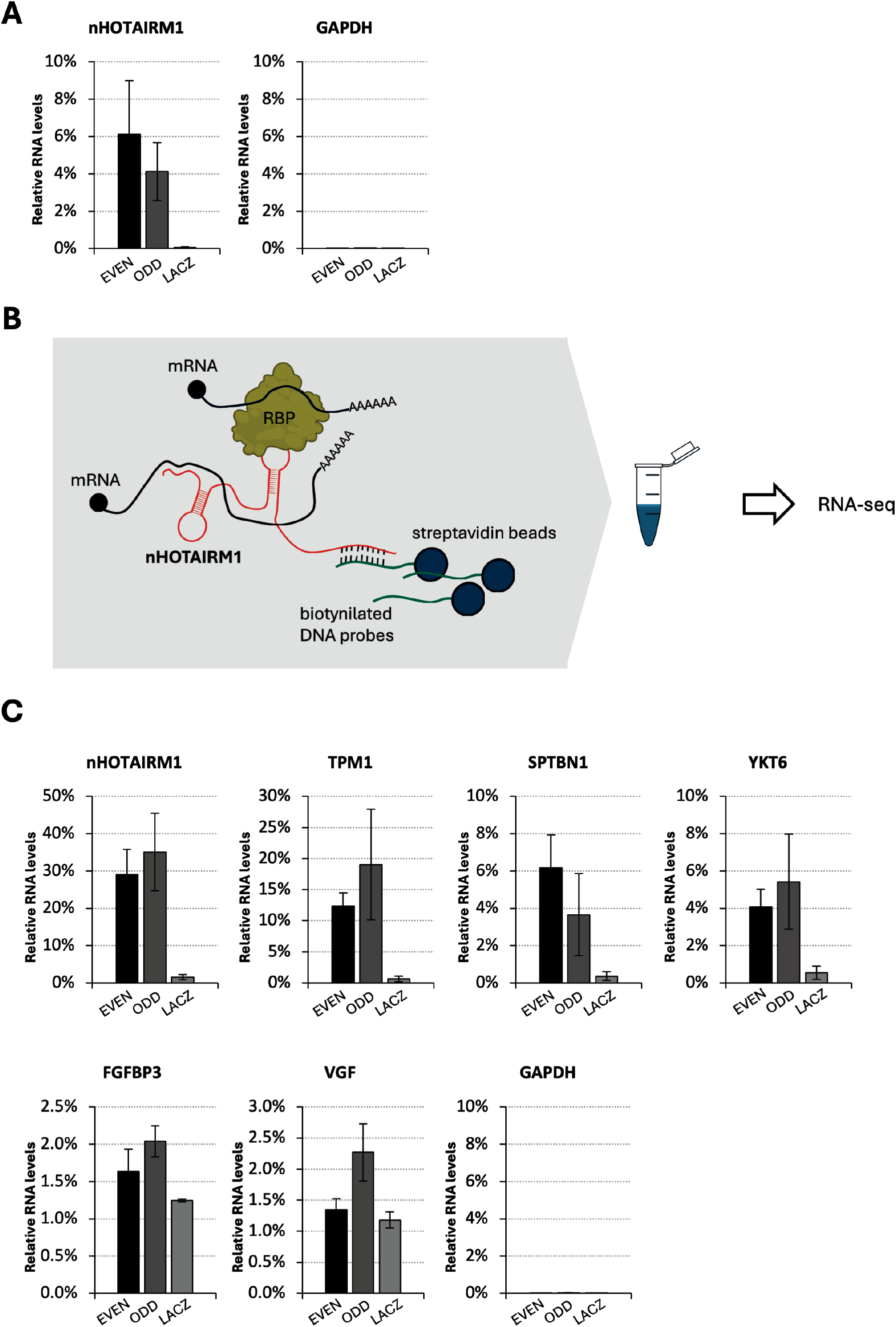
*nHOTAIRM1* native RNA pull-down sequencing in iPSC-derived spMNs. (A) qRT-PCR analysis of RNA enrichment in EVEN, ODD and LacZ fractions relative to input, from native *nHOTAIRM1* RNA pull-down experiments performed in iPSC-derived spMN extracts. Data are expressed as percentage of input. *GAPDH* was used as a negative control transcript. Data represent mean ± SEM from three independent biological replicates *(N = 3)*. (B) Schematic representation of the experimental workflow used to identify *nHOTAIRM1*-associated transcripts. Endogenous *nHOTAIRM1* (red) forms RNA-RNA interactions with candidate mRNAs (black) and associates with RNA-binding proteins (RBP). Biotinylated antisense DNA probes (green) complementary to *nHOTAIRM1* are hybridized to the lncRNA and captured using streptavidin-coated magnetic beads (blue), enabling the isolation of *nHOTAIRM1*-containing ribonucleoprotein complexes under native conditions. Co-purified RNAs are subsequently analyzed by RNA sequencing (RNA-seq) to identify enriched interacting transcripts. (C) qRT-PCR analysis of RNA enrichment in EVEN, ODD and LacZ fractions relative to input, from AMT-crosslinked *nHOTAIRM1* RNA pull-down experiments performed in living iPSC-derived spMNs. Data are expressed as percentage of input. *GAPDH* was used as a negative control transcript. Data represent mean ± SEM from three independent biological replicates *(N = 3)*.

Among the top 27 *nHOTAIRM1* RNA interactors identified, we focused on a subset of five mRNAs (*TPM1*, *YKT6*, *SPTBN1*, *VGF* and *FGFBP3*) with established relevance to MN-specific functions and/or axonal architecture and transport (Supplementary Fig. 5).^26–32^

Since native RNA pull-down assays can capture both direct (base complementarity-mediated) and indirect (protein-mediated) RNA-RNA associations, we next asked which of these transcripts could physically anneal with *nHOTAIRM1* in spMNs. We leveraged 4′-aminomethyltrioxsalen (AMT)-mediated crosslinking, which selectively stabilizes base-paired RNA duplexes occurring in living cells,^33^ followed by *nHOTAIRM1* RNA pull-down assays in spMNs.^33,34^ As shown in Fig. 2C, three out of five mRNAs, namely *TPM1*, *YKT6* and *SPTBN1*, were significantly enriched in *nHOTAIRM1*-specific fractions, supporting their identification as candidate direct RNA-RNA interactors of the lncRNA. The enrichment of these three mRNAs was also independently confirmed in native RNA pull-down fractions via qRT-PCR analysis (Supplementary Fig. 4B,C).

### *nHOTAIRM1* engages selected mRNA partners through discrete RNA-RNA pairing regions

To define candidate pairing interfaces between *nHOTAIRM1* and its mRNA partners, we performed an integrative computational analysis combining IntaRNA, RIsearch2 and RIME, and retained only consensus interaction regions supported by all three algorithms. This analysis identified discrete regions of *nHOTAIRM1* predicted to pair with *TPM1*, *YKT6* and *SPTBN1*, forming energetically favorable RNA-RNA duplexes (Supplementary Fig. 6). The confinement of these predictions to defined portions of the lncRNA suggested candidate pairing modules that could be functionally interrogated. With respect to the predicted ANXA11-contacting regions of *nHOTAIRM1* (Supplementary Fig. 2A), it is noteworthy that *nHOTAIRM1*-ANXA11 BD2 is compatible with the RNA-RNA duplexes predicted for *SPTBN1* and *TPM1* (Supplementary Figs. 6 and 7), whereas *nHOTAIRM1*-ANXA11 BD1 is compatible with the predicted *nHOTAIRM1*-*YKT6* duplex (Supplementary Figs. 6 and 7). This arrangement suggests that *nHOTAIRM1* may simultaneously accommodate target-mRNA pairing and ANXA11 binding, as illustrated in our model of the *nHOTAIRM1*-ANXA11-mRNA assembly. To test the contribution of these regions experimentally, we designed non-degradative DNA/LNA mixmer oligonucleotides, hereafter referred to as competitors, that sterically block the predicted *nHOTAIRM1* interaction sites (Supplementary Fig. 7). To enable steric competition in differentiated spMNs, we first established a SilentFect-based transfection protocol at day 12 that allowed reproducible delivery of the oligonucleotides under conditions compatible with subsequent biochemical analyses. A pool of four 20nt-long competitors was transfected at a final concentration of 100nM, and their effects were assessed after 48h by native *nHOTAIRM1* RNA pull-down with a single probe set, as described above (Fig. 3A). This analysis showed a consistent reduction in the recovery of the corresponding mRNA targets compared to cells treated with scramble (Fig. 3A).

**Figure 3.**
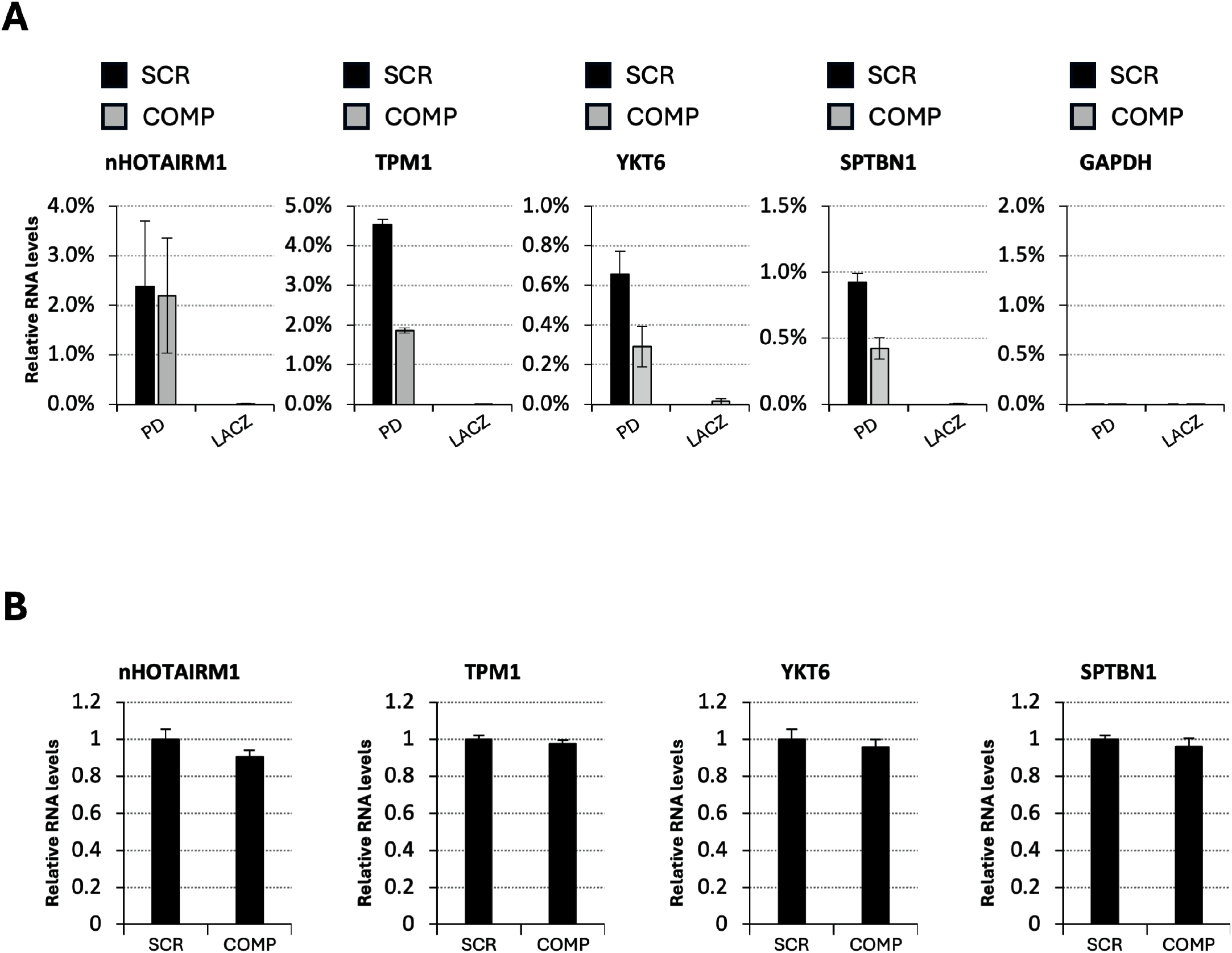
Steric blockade of predicted *nHOTAIRM1*-mRNA binding regions reduces target recovery in *nHOTAIRM1* RNA pull-down fractions. (A) qRT-PCR analysis of *nHOTAIRM1*, *TPM1*, *YKT6*, *SPTBN1* and *GAPDH* enrichment in *nHOTAIRM1* RNA pull-down (PD) and LacZ control fractions from spMNs transfected with either a scramble DNA/LNA steric-blocker (SCR) or an equimolar mixture of four DNA/LNA steric-blocking competitors (COMP) targeting the predicted *nHOTAIRM1* interaction regions involved in pairing with *TPM1*, *YKT6* and *SPTBN1*. Data are expressed as percentage of input. *GAPDH* was used as a negative control transcript. Data are presented as mean ± SEM from two independent biological replicates (*N* = 2). (B) qRT-PCR analysis of total cellular *nHOTAIRM1*, *TPM1*, *YKT6* and *SPTBN1* RNA levels in SCR- and COMP-transfected spMNs. RNA levels were normalized to the corresponding scramble-control condition. Data are presented as mean ± SEM from two independent biological replicates (*N* = 2).

Importantly, competitor treatment did not significantly alter the total cellular levels of *nHOTAIRM1*, *TPM1*, *YKT6*, or *SPTBN1* (Fig. 3B), indicating that the reduced recovery of the mRNA partners reflected disruption of their association with the lncRNA rather than changes in transcript abundance. *nHOTAIRM1* recovery in the RNA pull-down fraction was also unaffected, excluding an effect of the competitors on lncRNA capture efficiency. Together, computational mapping and site-directed steric competition functionally support the relevance of the predicted interaction regions and provide convergent evidence consistent with direct RNA-RNA pairing between *nHOTAIRM1* and *TPM1*, *YKT6*, and *SPTBN1* transcripts.

### *nHOTAIRM1* spatially colocalizes with its mRNA interactors in spMNs

We next asked whether *nHOTAIRM1* could contribute, together with the transport factor ANXA11, to the subcellular localization of its mRNA partners.

We first performed double-target RNA-FISH to examine the spatial relationship between the lncRNA and each transcript. Representative images revealed discrete colocalization events between *nHOTAIRM1* and *TPM1* (Fig. 4A), *YKT6* (Fig. 5A), or *SPTBN1* (Fig. 6A) both the soma and neurite projections.

**Figure 4.**
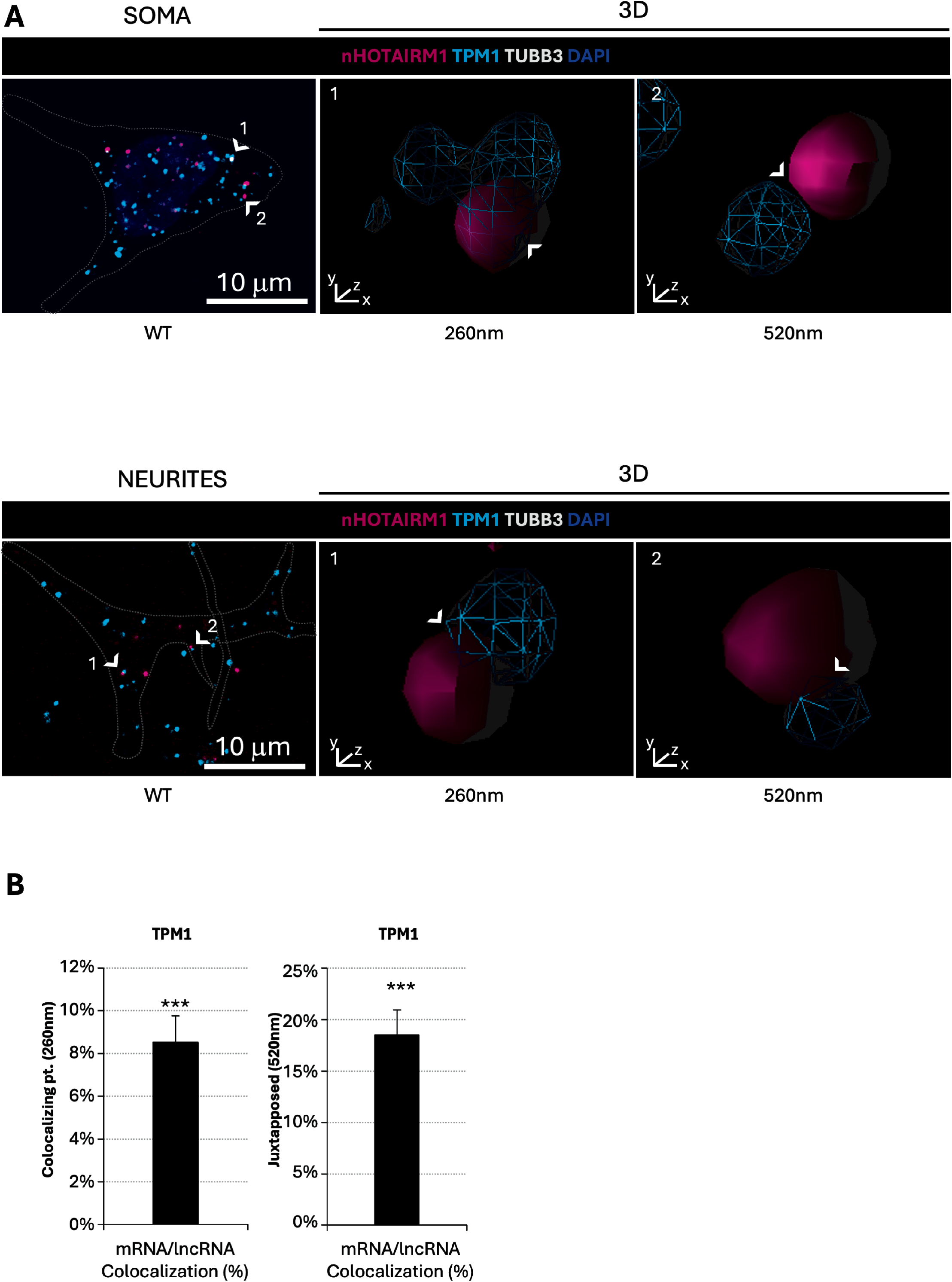
Colocalization of *nHOTAIRM1* with *TPM1* mRNA in iPSC-derived spMNs. (A) 3D surface reconstructions showing colocalization events between *nHOTAIRM1* and *TPM1* mRNA. Representative double-target RNA-FISH for *nHOTAIRM1* (magenta) and *TPM1* mRNA (cyan), combined with IF for TUBB3 (gray) and DAPI (blue) in WT spMNs. Upper panels (SOMA): Confocal maximum projection (left) and 3D renderings of the RNA-RNA interaction events indicated by white arrowheads (center and right) illustrating RNA puncta classified as colocalizing (≤260 nm) or adjacent (≤520 nm). The dashed line delineates soma boundaries. Lower panels (NEURITES): Maximum projections (left) and 3D renderings of the RNA-RNA interaction events indicated by white arrowheads (center and right) showing spatial proximity between *nHOTAIRM1* and *TPM1* mRNA within neuritic compartments. The dashed line delineates neurite boundaries. Images were acquired at 100x magnification using confocal microscopy with a Z-step size of 0.2□µm. Scale bars: 10□µm. (B) Quantification of mRNA/lncRNA colocalization events in somata and neurites. Percentages represent fully overlapping particles (≤260□nm, left) and adjacent particles (≤520□nm, right). Distance thresholds were defined based on the optical resolution limit. Statistical significance was determined by one-sample two-tailed Student’s t test using biological replicates as independent units. *P < 0.05, **P < 0.01, ***P < 0.001.

**Figure 5.**
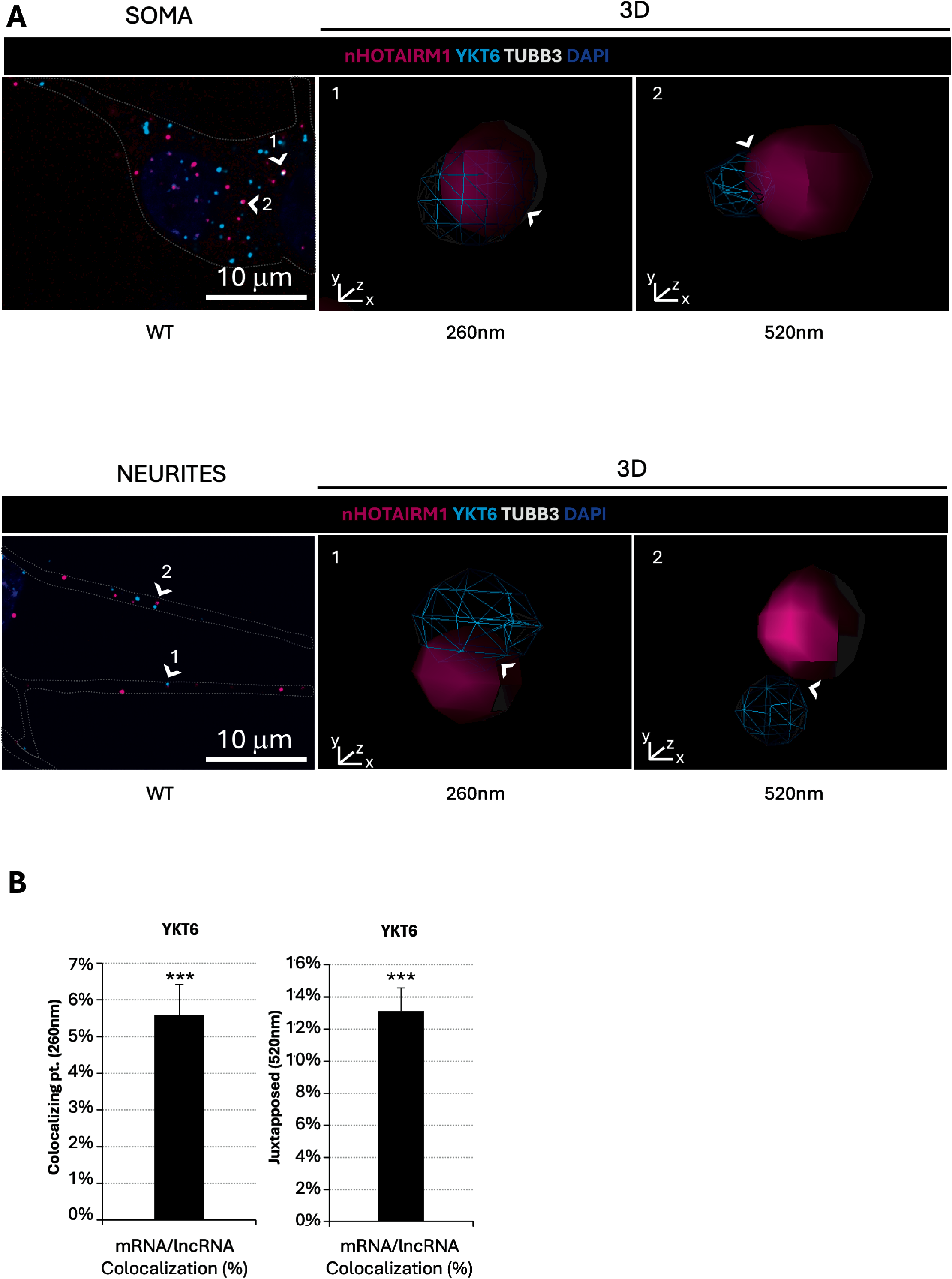
Colocalization of *nHOTAIRM1* with *YKT6* mRNA in spMNs. (A) 3D surface reconstructions showing colocalization events between *nHOTAIRM1* and *YKT6* mRNA. Representative double-target RNA-FISH for *nHOTAIRM1* (magenta) and *YKT6* mRNA (cyan), combined with IF for TUBB3 (gray) and DAPI (blue) in WT spMNs. Upper panels (SOMA): Confocal maximum projection (left) and 3D renderings of the RNA-RNA interaction events indicated by white arrowheads (center and right) illustrating RNA puncta classified as colocalizing (≤260 nm) or adjacent (≤520 nm). The dashed line delineates soma boundaries. Lower panels (NEURITES): Maximum projections (left) and 3D renderings of the RNA-RNA interaction events indicated by white arrowheads (center and right) showing spatial proximity between *nHOTAIRM1* and *YKT6* mRNA within neuritic compartments. The dashed line delineates neurite boundaries. Images were acquired at 100x magnification using confocal microscopy with a Z-step size of 0.2□µm. Scale bars: 10□µm. (B) Quantification of mRNA/lncRNA colocalization events in somata and neurites. Percentages represent fully overlapping particles (≤260□nm, left) and adjacent particles (≤520□nm, right). Distance thresholds were defined based on the optical resolution limit. Statistical significance was determined by one-sample two-tailed Student’s t test using biological replicates as independent units. *P < 0.05, **P < 0.01, ***P < 0.001.

**Figure 6.**
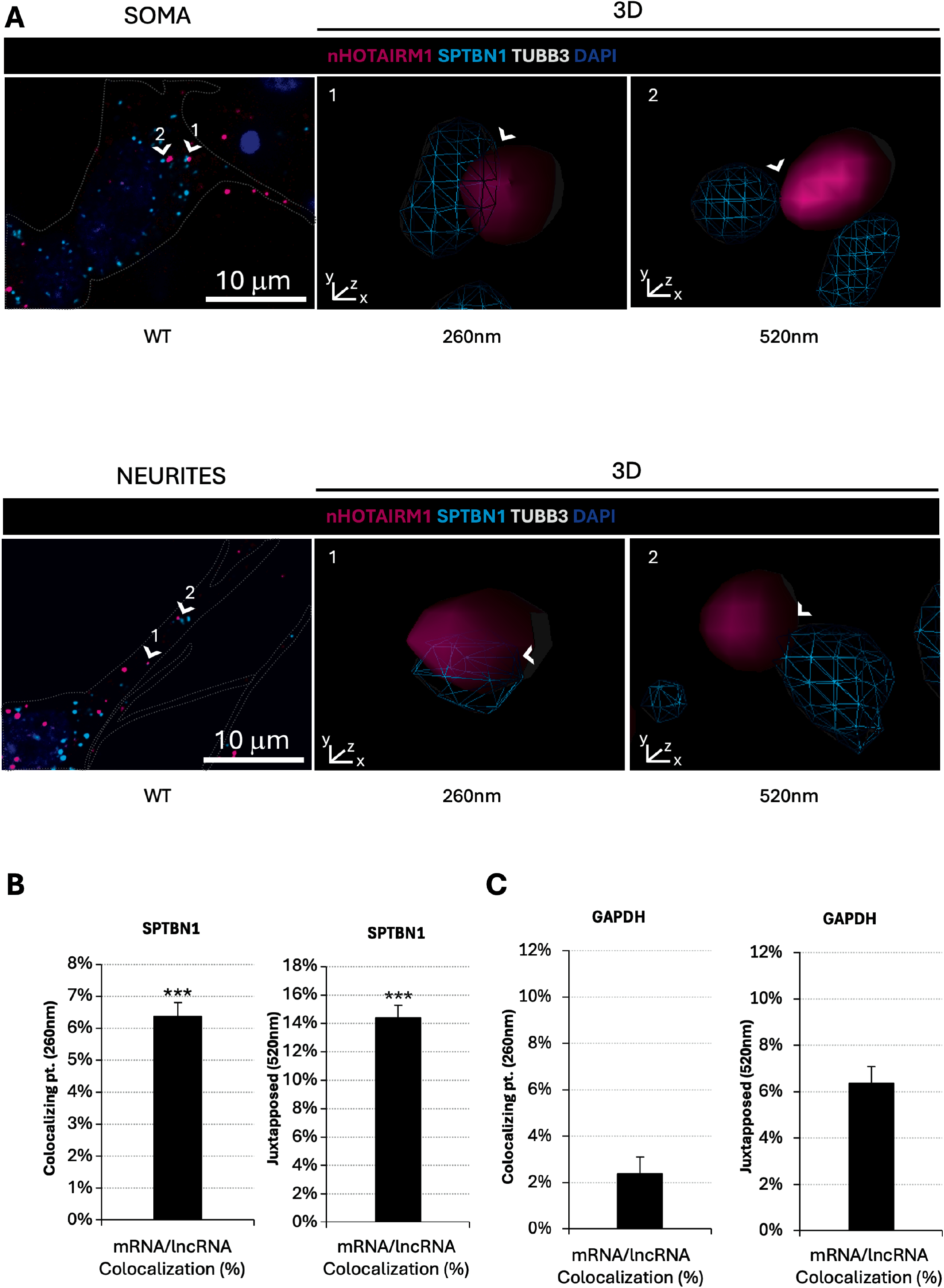
Colocalization of *nHOTAIRM1* with SPTBN1 mRNA in spMNs. (A) 3D surface reconstructions showing colocalization events between *nHOTAIRM1* and *SPTBN1* mRNA. Representative double-target RNA-FISH for *nHOTAIRM1* (magenta) and *SPTBN1* mRNA (cyan), combined with IF for TUBB3 (gray) and DAPI (blue) in WT spMNs. Upper panels (SOMA): Confocal maximum projection (left) and 3D renderings of the RNA-RNA interaction events indicated by white arrowheads (center and right) illustrating RNA puncta classified as colocalizing (≤260 nm) or adjacent (≤520 nm). The dashed line delineates soma boundaries. Lower panels (NEURITES): Maximum projections (left) and 3D renderings of the RNA-RNA interaction events indicated by white arrowheads (center and right) showing spatial proximity between *nHOTAIRM1* and *SPTBN1* mRNA within neuritic compartments. The dashed line delineates neurite boundaries. Images were acquired at 100x magnification using confocal microscopy with a Z-step size of 0.2□µm. Scale bars: 10□µm. (B) Quantification of mRNA/lncRNA colocalization events in somata and neurites. Percentages represent fully overlapping particles (≤260□nm, left) and adjacent particles (≤520□nm, right). Distance thresholds were defined based on the optical resolution limit. Statistical significance was determined by one-sample two-tailed Student’s t test using biological replicates as independent units. (C) Quantification of *nHOTAIRM1*/*GAPDH* proximity events under the same imaging and analysis conditions. *GAPDH* was used as a negative-control transcript. Histograms show the percentage of fully overlapping RNA puncta (≤260□nm, left) and adjacent RNA puncta (≤520□nm, right). Data are presented as mean□±□SEM from three independent biological replicates. *P < 0.05, **P < 0.01, ***P < 0.001.

To quantify the extent of spatial proximity between the molecules, we measured the percentage of signal pairs separated by ≤260 nm (Figs. 4B, 5B and 6B, left panel) and ≤520 nm (Figs. 4B, 5B and 6B, right panel), using previously validated nanoscale proximity criteria.^35^ This analysis enabled us to distinguish fully overlapping from closely juxtaposed signals (see Fig. 6C for negative control quantification).

A discrete fraction of *nHOTAIRM1* molecules was found in nanoscale proximity to *TPM1*, *YKT6* and *SPTBN1* transcripts in spMNs (see Supplementary Figs. 8-10 for digital enlargements), consistent with the formation of mRNA-lncRNA assemblies potentially underpinning *nHOTAIRM1*-dependent control of RNA localization and trafficking.

Although AMT-crosslinked RNA pull-down, computational mapping, and steric-blocking competition assays supported direct *nHOTAIRM1*-mRNA pairing, these approaches did not formally exclude the contribution of additional intermediary RNAs. We therefore searched the stringently defined *nHOTAIRM1* interactome identified by RNA pull-down RNA-seq (Supplementary Fig. 11A) for candidate mediator RNAs potentially able to bridge the lncRNA and each of its three mRNA interactors.

Candidate mediators were selected through an integrated strategy requiring reproducible enrichment with *nHOTAIRM1* and predicted interactions with RNA partners through distinct, non-overlapping regions (see analytical details in Materials and Methods).

Starting from *nHOTAIRM1* regions predicted to interact with *TPM1*, *YKT6*, and *SPTBN1* (Supplementary Fig. 6), we searched for potential mediator RNAs by analyzing each *nHOTAIRM1*-mRNA pair from the list of *nHOTAIRM1* mRNA interactors selected by the RNA pull-down sequencing analysis.

Guided by *nHOTAIRM1* sequence, we screened for candidate mediator RNAs capable of establishing a bridging RNA-RNA interaction between the lncRNA and each of its three validated mRNA partners. Candidates were prioritized based on their predicted ability to interact with both *nHOTAIRM1* and the corresponding mRNA through distinct, sterically compatible RNA-RNA interfaces and on a more favorable predicted affinity (interaction energy) for the mRNA than the corresponding direct *nHOTAIRM1-mRNA* interaction (see Materials and Methods). This analysis identified *RNF5* as a candidate mediator for the *nHOTAIRM1*-*SPTBN1* association, *ZC3H18* for *nHOTAIRM1*-*TPM1*, and *POLR1E* for *nHOTAIRM1*-*YKT6*. *FAM98A* emerged as a shared candidate predicted to bridge *nHOTAIRM1* with each of the three mRNA partners (Supplementary Fig. 11B,C).

To test whether these potential mediator transcripts were present within *nHOTAIRM1*-mRNA assemblies, we performed triple-target RNA-FISH assays, simultaneously detecting *nHOTAIRM1*, one mRNA partner, and the corresponding putative mediator RNA (workflow shown in Supplementary Fig. 12). *RNF5* was analyzed with *nHOTAIRM1* and *SPTBN1*, *ZC3H18* with *nHOTAIRM1* and *TPM1*, *POLR1E* with *nHOTAIRM1* and *YKT6*, and *FAM98A* separately with each of the three *nHOTAIRM1*-mRNA pairs (Supplementary Fig. 13A-F, lower panels).

In each combination, the percentage of colocalizing particles measured in the triple-target RNA-FISH assays closely matched that previously observed by double-target RNA-FISH, thereby providing an internal experimental control that confirmed the preservation of the established spatial proximity patterns (Supplementary Fig. 13A-F, upper panels, left chart). We then quantified the occurrence of each candidate mediator at these sites. Despite robust detection of all transcripts (Supplementary Figs. 14 and 15), none was detected at the corresponding *nHOTAIRM1*-mRNA colocalization site in any of the six combinations examined, resulting in no detectable triple-colocalization events (Supplementary Fig. 13A-F, upper panels, right chart). These findings support a model in which the detected associations of *nHOTAIRM1* with *TPM1*, *YKT6*, and *SPTBN1* do not require a stable RNA bridge, at least among the prioritized candidates and under the conditions tested.

### *nHOTAIRM1* promotes the neuritic localization of its mRNA partners and their recruitment into ANXA11-positive complexes

Having extensively established, through double- and triple-target RNA-FISH and AMT-crosslinked RNA pull-down assays, the direct interaction between *nHOTAIRM1* and its cargo mRNAs, and having also demonstrated the interaction between *nHOTAIRM1* and ANXA11 through complementary biochemical, imaging and computational approaches, we next investigated whether the lncRNA is required to recruit its three partner transcripts into ANXA11-containing RNP complexes and contribute to their neuritic localization.

To this end, we quantified via qRT-PCR the enrichment of the three interactor mRNAs *TPM1*, *YKT6* and *SPTBN1* in ANXA11 CLIP fractions obtained in parallel from WT spMNs and from *nHOTAIRM1* KO counterparts generated in our previous study.^17^ *TPM1*, *YKT6* and *SPTBN1* were robustly enriched in ANXA11-bound CLIP fractions from WT cells, whereas their recovery was reduced upon *nHOTAIRM1* loss (Fig. 7A). Importantly, depletion of the lncRNA did not significantly alter the total cellular levels of the three transcripts (Fig. 7B), indicating that their reduced recovery (Fig. 7A) reflected impaired association with ANXA11-containing complexes rather than changes in transcript steady-state abundance.

**Figure 7.**
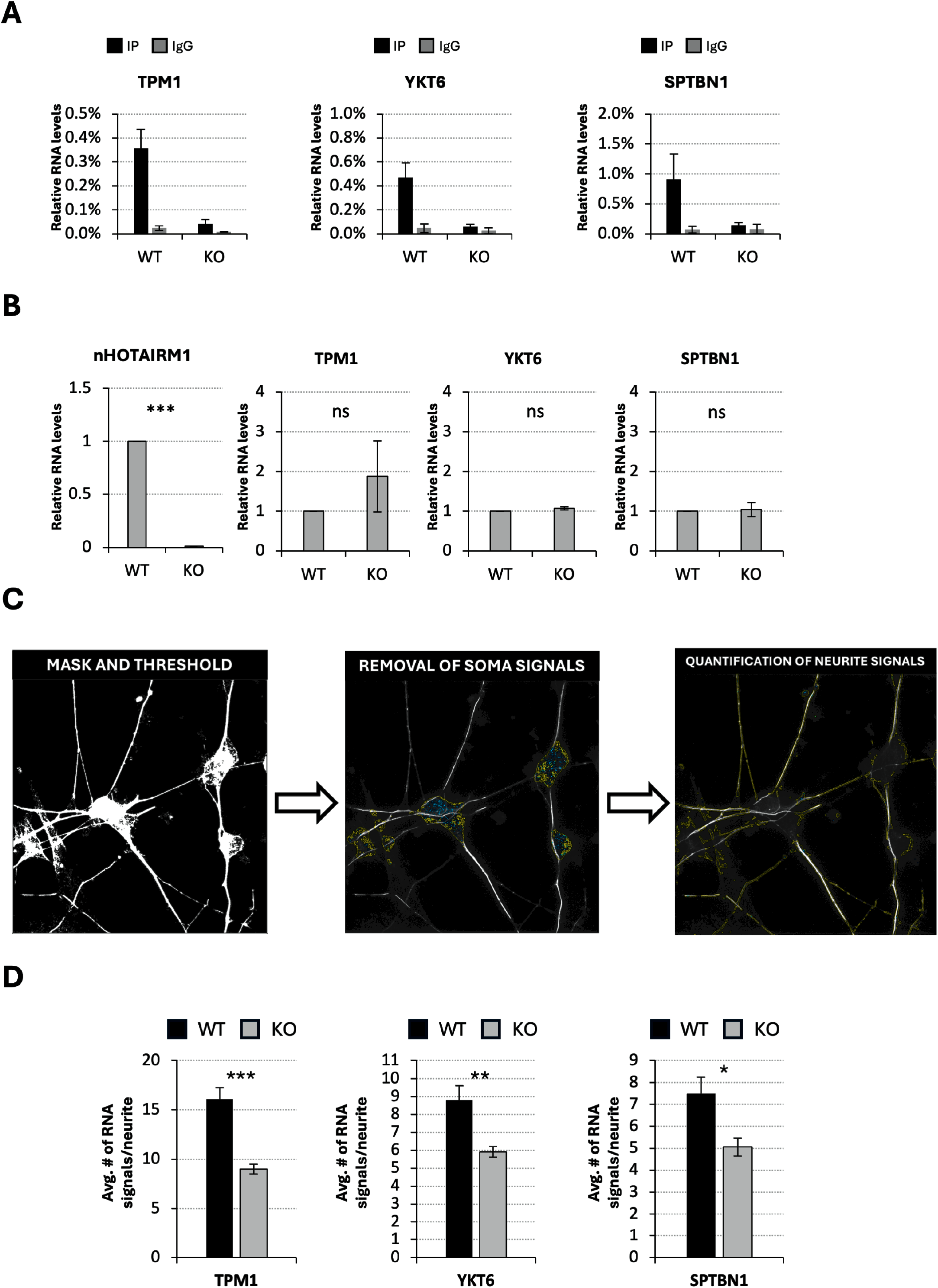
*nHOTAIRM1*-dependent association and neuritic localization of target mRNAs. (A) qRT-PCR analysis of *TPM1*, *YKT6* and *SPTBN1* mRNA enrichment in ANXA11 CLIP assays performed in WT and *nHOTAIRM1* KO spMNs. RNA enrichment is expressed as percentage of input. Experimental control is shown in Fig. 2A. Data represent mean ± SEM from three independent biological replicates *(N = 3)*. (B) qRT-PCR analysis of total *TPM1*, *YKT6* and *SPTBN1* mRNA expression levels in WT and *nHOTAIRM1* KO spMNs. Data are expressed in arbitrary units, relative to housekeeping gene control (*ATP5O*). Data represent mean ± SEM from three independent biological replicates *(N = 3)*. (C) Schematic representation of soma and neurite compartmentalization used for imaging-based quantification analyses in spMNs. (D) Quantification of the average number of RNA-FISH puncta per neurite for *TPM1*, *YKT6* and *SPTBN1* in WT and *nHOTAIRM1* KO spMNs. ≥50 cells per condition, derived from at least three independent biological replicates, were analyzed. Data are presented as mean□±□SEM. Statistical analyses were performed using biological replicates as independent units and significance was determined by unpaired two-tailed Student’s t test. *P < 0.05, **P < 0.01, ***P < 0.001.

We further leveraged *nHOTAIRM1* KO spMNs to investigate whether defective loading of cargoes into ANXA11-positive assemblies was accompanied by altered neuritic localization. We set up a neurite-restricted RNA imaging pipeline (Fig. 7C) in which (i) extra-cellular and background signals were excluded, (ii) somatic regions were masked out, and (iii) RNA-FISH puncta were quantified specifically within neurite projections. As shown in Fig. 7D, we found that *nHOTAIRM1* loss significantly reduced the average number of *TPM1*, *YKT6*, and *SPTBN1* RNA-FISH spots per neurite compared with WT spMNs, confirming that *nHOTAIRM1* is required to sustain both the association of these transcripts with ANXA11-containing complexes and their peripheral accumulation within neuronal processes.

### ANXA11 contributes to the neuritic localization of *nHOTAIRM1* mRNA cargoes

We finally asked whether ANXA11, the transport factor complexed with *nHOTAIRM1* and its associated transcripts, participated into RNA cargo neuritic localization. iPSC-derived spMNs were transfected at day 10 of differentiation with an equimolar mix of two independent siRNAs targeting distinct regions of ANXA11 transcript and analyzed 48 hours later. Western blot analysis confirmed an approximately 50% reduction in ANXA11 protein levels relative to

GAPDH (Supplementary Fig. 16A), whereas qRT-PCR, while confirming ANXA11 depletion at the RNA level, showed no significant changes in the total cellular levels of *nHOTAIRM1*, *TPM1*, *YKT6* and *SPTBN1* (Supplementary Fig. 16B). We then asked whether ANXA11 contributes functionally to the subcellular localization of these transcripts. We analyzed the neuritic distribution of *TPM1*, *YKT6*, and *SPTBN1* using the same neurite-restricted RNA-imaging pipeline already applied to *nHOTAIRM1* KO conditions. ANXA11 depletion significantly reduced the average number of *TPM1*, *YKT6*, and *SPTBN1* RNA-FISH puncta per neurite compared with the scramble-treated controls (Fig. 8A-C and Supplementary Figs. 17-19). Since the total cellular abundance of these transcripts, including *nHOTAIRM1*, was unaffected by ANXA11 knockdown (Supplementary Fig. 16B), their reduced neuritic accumulation reflects altered subcellular distribution rather than decreased expression levels. These results show that ANXA11 contributes to the efficient localization of *nHOTAIRM1*-associated mRNA cargoes within spMN projections.

**Figure 8.**
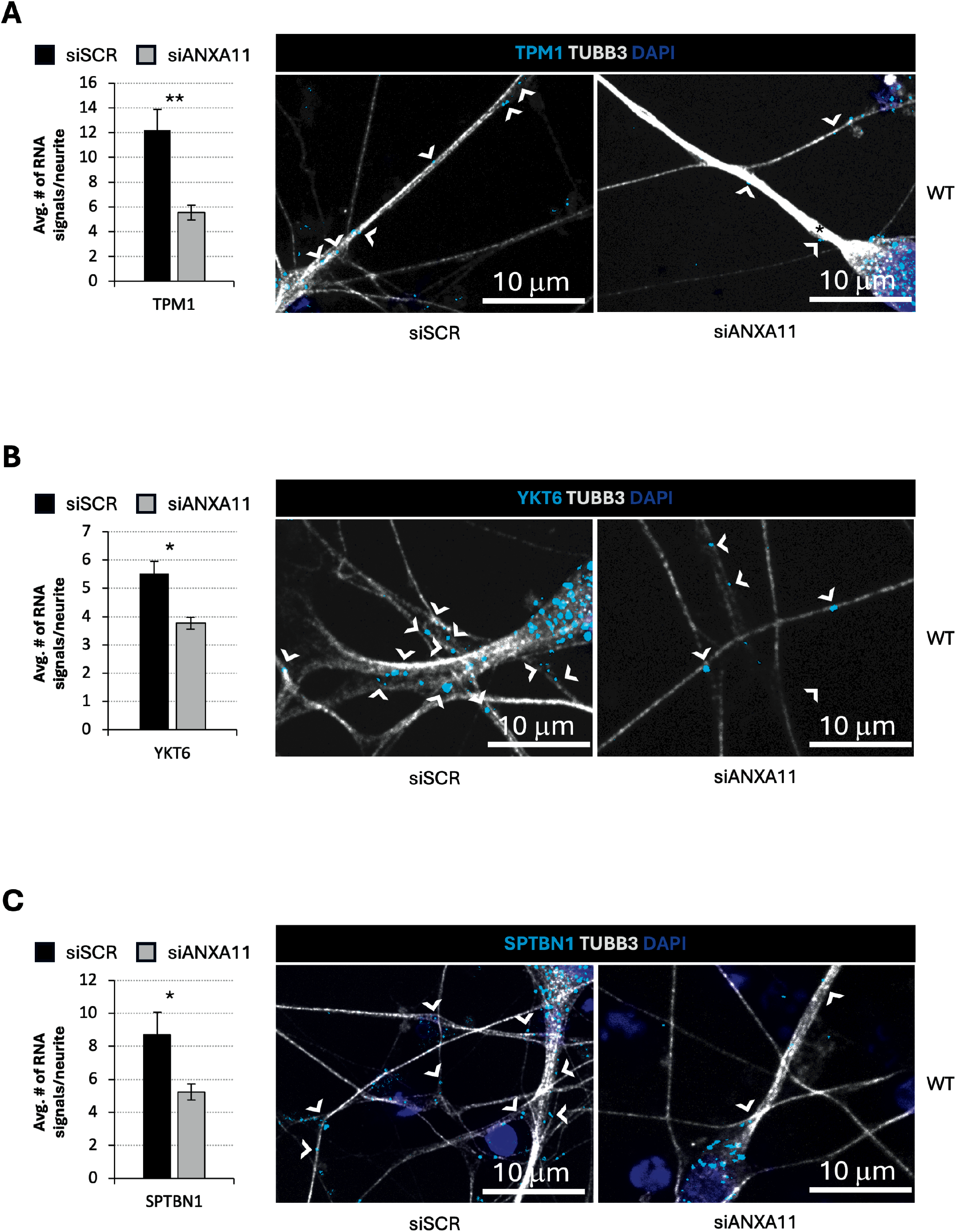
ANXA11 depletion impairs the neuritic localization of *nHOTAIRM1*-associated mRNA cargoes. (A-C) Quantification and representative RNA-FISH images of *TPM1* (A), *YKT6* (B) and *SPTBN1* (C) in WT spMNs transfected with scramble siRNA (siSCR) or an equimolar mixture of two siRNAs targeting ANXA11 (siANXA11). Histograms show the average number of RNA-FISH puncta per neurite. Representative images show RNA-FISH detection of the indicated mRNA (cyan), combined with IF for TUBB3 (gray) and DAPI nuclear staining (blue). Arrowheads indicate representative neuritic RNA puncta. At least 50 cells per condition from three independent biological replicates were analyzed. Data are presented as mean□±□SEM from three independent biological replicates (*N* = 3). Statistical significance was determined by unpaired two-tailed Student’s t test using biological replicates as independent units. Scale bars: 10□µm. *P < 0.05, **P < 0.01, ***P < 0.001.

Taken together, these complementary loss-of-function experiments support a transport axis (Supplementary Fig. 20) in which *nHOTAIRM1* promotes the selective incorporation of a defined set of axonal/synaptic mRNAs into ANXA11-positive RNA assemblies while ANXA11 contributes to their efficient localization within spMN projections, revealing an unappreciated role for lncRNAs in governing axonal RNA trafficking in human MNs.

## Discussion

Neuronal physiology depends on tightly regulated spatial gene expression. Cellular polarity, axonal growth, synaptic maintenance and cue-dependent plasticity require rapid responses that cannot rely solely on somatic protein delivery. A central mechanism enabling this compartmentalization is the subcellular partitioning of mRNAs, which influences local transcript fate and thereby sculpts the proteome of distal axonal and synaptic domains.^2,36,37^ This constraint is particularly pronounced in spMNs, where long projections magnify the impact of even modest perturbations in RNA delivery, retention and utilization, and intersect with pathways implicated in MN vulnerability.^38^

Axonal mRNA trafficking is commonly framed as an RBP-centric process, in which cis-acting localization elements recruit RBPs that assemble transport-competent RNP granules and couple them to molecular motors.^39,40^ While well supported by the literature, this framework leaves the mechanisms governing cargo selection and coordinated co-transport incompletely resolved. RNP granules are increasingly viewed as multicomponent, dynamic assemblies sustained by weak, multivalent interactions, raising the possibility that noncoding RNAs could contribute to organizational functions that cannot be easily recapitulated by protein-only mechanisms.^41^ In this context, lncRNAs are plausible determinants of granule composition and cargo specificity, as they can provide binding platforms for proteins while engaging selected RNA partners through sequence- and structure-dependent interactions.^42–45^ Although several lncRNAs have been detected in neuronal projections and implicated in local translation or cytoskeletal remodelling,^46^ mechanistic examples that connect a defined lncRNA to a specific axonal transport pathway and RNA cargo set remain limited, particularly in human MNs.

Against this background, our findings identify *nHOTAIRM1* as a transport-related adaptor operating in human iPSC-derived spMNs. Previous studies have associated this neuronal *HOTAIRM1* isoform with lineage specification^16^ and established its contribution to MN identity, neurite architecture and synaptic activity.^17^ The present work extends this biological framework by linking *nHOTAIRM1* to the selective partitioning of MN-relevant mRNAs.

*nHOTAIRM1* engages selected mRNA cargoes (*TPM1*, *SPTBN1* and *YKT6*) through lncRNA-mRNA interactions and promotes their delivery to distal neuronal compartments by coupling them to the ANXA11-dependent axonal transport machinery. In this view, the trafficking defects described here do not represent an isolated phenotype but provide a potential mechanistic layer contributing to the broader alterations in neurite branching, neuronal maturation and synaptic gene expression previously observed in the *nHOTAIRM1* KO model.^17^ The identity of the selected mRNA partners supports the functional coherence of this transport axis. *TPM1* and *SPTBN1* participate in actin-spectrin organization and neuronal structural integrity,^30,32,47^ whereas *YKT6* contributes to membrane fusion and vesicular trafficking.^29^ Their coordinated recruitment may therefore connect transport granule composition with cytoskeletal maintenance and membrane dynamics in distal neuronal compartments. Although not tested here, the perturbation of functionally related mRNAs may provide a plausible link between defective RNA trafficking and the previously observed alterations in neurite architecture and synaptic programs.^17^

Mechanistically, our data support a model in which *nHOTAIRM1* combines two complementary interaction properties: association with the lysosome-linked transport factor ANXA11^18^ and sequence-specific recognition of selected mRNA partners. Together, these interactions are consistent with a scaffolding function,^8,42,48^ in which *nHOTAIRM1* contributes to the inclusion of specific transcripts in ANXA11-linked RNP granules and their enrichment in distal neurites. To our knowledge, this represents one of the first mechanistic examples of a lncRNA contributing to axonal mRNA selection through direct RNA-RNA pairing.

Evidence for *nHOTAIRM1*-mRNA pairing is supported by convergent biochemical, computational, functional, and spatial evidence. Site-directed competition selectively reduced the association of *TPM1*, *YKT6*, and *SPTBN1* mRNAs with *nHOTAIRM1* without altering their abundance or that of the lncRNA, supporting the functional relevance of the predicted pairing regions. Further Identification of discrete *nHOTAIRM1* interaction modules suggests that cargo selection relies on modular RNA-binding surfaces, with direct RNA-RNA contacts contributing to multivalent RNP organization.^49^

This model is further supported by analysis of potential intermediary RNAs. Although computational predictions identified candidate bridging transcripts, triple-target RNA-FISH did not detect their stable recruitment to *nHOTAIRM1*-mRNA association sites, indicating that the predominant interactions in spMNs do not require stable RNA mediators, while not excluding the possibility of transient or low-abundance intermediates.

On the protein side, ANXA11 CLIP and reciprocal *nHOTAIRM1* RNA pull-down provide complementary biochemical evidence for the association of the two molecules. Together with their spatial colocalization, these RNA- and protein-centered approaches support the inclusion of *nHOTAIRM1* in ANXA11-positive assemblies without defining the molecular interface.

Our catRAPID predictions are consistent with a recent independent in silico analysis of ANXA11, ^19^ which identified three regions of maximal RNA-binding propensity centered at approximately residues 290, 410 and 460 within the C-terminal annexin core. Accordingly, multiple high-propensity *nHOTAIRM1*-binding regions predicted in our study (Supplementary Fig. 2A) overlap those regions around residues ∼300 and ∼400, supporting convergence on the same RNA-binding surfaces. This is further supported by recent biochemical evidence showing that the ANXA11 C-terminal core co-binds RNA and lipid vesicles in a Ca²⁺-dependent manner.^50^ Direct experimental mapping using targeted ANXA11 variants and complementary nHOTAIRM1 fragments will be necessary to define the biochemical determinants of this association.

The functional relationship between *nHOTAIRM1* and ANXA11 is supported by the parallel effects of their depletion. Loss of *nHOTAIRM1* markedly reduced the recovery of *TPM1*, *YKT6* and *SPTBN1* in ANXA11-associated fractions, despite unchanged total mRNA levels, indicating that the lncRNA contributes to their efficient cargo loading into ANXA11-containing pools. Conversely, partial ANXA11 depletion reduced the neuritic accumulation of the same transcripts without altering their overall abundance or *nHOTAIRM1* levels. The similar localization defects observed upon *nHOTAIRM1* loss and ANXA11 knockdown support their involvement in a shared transport pathway.

The different sizes of the effects detected by ANXA11 CLIP and neuritic RNA-FISH upon *HOTAIRM1* KO likely reflect the distinct biological layers captured by the two assays. CLIP specifically quantifies transcripts associated with ANXA11-containing assemblies, whereas RNA-FISH measures the total steady-state neuritic RNAs, which also reflects alternative transport pathways, local retention, transcript stability, and pre-existing RNAs. Consequently, a marked reduction in ANXA11-associated recovery can coexist with a more modest decrease in total neuritic signal. Consistent with this, depletion of either *nHOTAIRM1* or ANXA11 reduced, but did not abolish, neuritic localization, indicating that both promote efficient cargo recruitment and transport rather than acting as the sole determinants of mRNA localization. More broadly, the findings suggest a model for lncRNA-guided cargo selection in neurons. The multivalent interaction capacity of *nHOTAIRM1* may allow it to associate concurrently with a transport-linked protein and selected mRNAs, thereby influencing the RNA composition of ANXA11-positive assemblies. Such a mechanism would add a layer of cargo specificity to canonical RBP-dependent transport without requiring *nHOTAIRM1* to be an obligatory component of all neuronal RNA-transport events.

The involvement of ANXA11 also raises regulatory possibilities. ANXA11 is a Ca²⁺-responsive and membrane-associated protein that tethers RNP granules to a lysosomal transport machinery sensitive to changes in cellular state, including neuronal activity and stress.^51,52^ *nHOTAIRM1*-dependent RNA cargo loading onto ANXA11-positive assemblies could therefore represent a point of convergence between signaling and metabolic inputs and RNA dynamics. Beyond physiology, our findings can be relevant to MN pathology. During development and regeneration, proper axonal mRNA localization supports cytoskeletal integrity, synapse formation, and growth cone dynamics, and its disruption can alter neuronal connectivity.^37,53^ On the other hand, defects in axonal transport are increasingly viewed as contributors to degeneration rather than solely downstream consequences.^38,53,54^ ANXA11 mutations have been linked to ALS^18,55^ and can impair axonal transport while promoting aggregation-prone states; consequently, pathways that depend on ANXA11-mediated granule trafficking may be vulnerable in disease settings.^56^ Even though we have not tested ALS-associated ANXA11 variants or patient-derived neurons, the work provides a basis for targeted follow-up experiments assessing whether disruption of either *nHOTAIRM1* function or ANXA11 interface amplifies transport defects under disease-relevant stressors.

In conclusion, our findings support a model in which *nHOTAIRM1* contributes to axonal RNA logistics by coupling the transport factor ANXA11 to a defined set of MN-relevant mRNA cargoes through a combination of RNA-protein interaction and direct RNA-RNA pairing. This mechanism expands current views of neuronal RNA trafficking by illustrating how a lncRNA can participate in RNA cargo selection and distal enrichment. More broadly, it provides a tractable framework to investigate how lncRNA-guided RNP transport supports MN architecture and how perturbations of this layer may intersect with early transport-related vulnerability in MN disorders.

## Materials and methods

### MN differentiation

Human iPSCs were derived, maintained, and induced to differentiate into spMNs following the methods described in^57^.

### Immunofluorescence and double-target RNA-FISH assay

Immunofluorescence and double-target RNA-FISH assays were performed as described in^58^ and^59^ with minor modifications.

Briefly, iPSC-derived spMNs at D5 were plated on 12 mm-diameter glass coverslips pre-coated with 0.01% poly-L-ornithine (Sigma-Aldrich) overnight at 37°C followed by murine laminin (20 μg/ml, Sigma-Aldrich) and differentiated until D12. Then cells were fixed in 4% paraformaldehyde (Electron Microscopy Sciences, cat#15710) in PBS for 20 min at 4°C, permeabilized with 0.1-0.2% Triton X-100 in PBS for 15-30 min at RT, and blocked with 3% BSA/10% goat serum in PBS for 30 min at RT. For immunofluorescence (IF), primary antibodies (anti-TUBB3 1:300, anti-ANXA11 1:200) were incubated overnight at 4°C in blocking buffer. Samples were washed 3X with PBS and incubated with secondary antibodies (goat anti-mouse 488 1:300 Invitrogen, cat#A-11029; donkey anti-rabbit 594 1:300 Invitrogen, cat#A-21207) for 45 min at RT.

For RNA-FISH, HCR™ RNA-FISH technology (Molecular Instruments) was used according to^58,59^. For detection of *nHOTAIRM1, TPM1, YKT6* and *SPTBN1* transcripts, cells were hybridized with a specific set of probes (50 nM each) overnight at 37°C (Supplementary Table S2 reports the number of RNA-FISH probes used; sequences are available upon request), sequences are indicated in) and then with amplifier probes conjugated to 647 or 594 fluorophore for 3 h at room temperature. Immunofluorescence was performed sequentially as above. The nuclei were counterstained with DAPI (1 μg/ml, Sigma-Aldrich cat#D9542) for 5 min, and mounted with ProLong Diamond Antifade Mountant (Thermo Fisher, cat#P36961). Images were acquired on an inverted Olympus IX73 confocal microscope equipped with Crestoptics X-LIGHT V3 spinning disk system, Prime BSI Express Scientific CMOS camera, 100X (NA 1.3) oil objective, Z-step size of 0.2 µm, using MetaMorph software (Molecular Devices).

Post-acquisition processing was performed by FIJI tools: in particular, after background setting, spatial proximity of RNA signals was evaluated as centroid-to-centroid distance on maximum Z-projection by the Spots colocalization (ComDet) plug-in.

### Triple-target RNA-FISH assay

Triple-target RNA fluorescence in situ hybridization (RNA-FISH) assays were performed to simultaneously detect *nHOTAIRM1*, validated *nHOTAIRM1*-interacting mRNAs and predicted candidate mediator RNAs in spMNs.

Triple-target RNA-FISH assays were performed as described above for double-target RNA-FISH with minor modifications. Briefly, iPSC-derived spMNs at day 12 of differentiation were fixed with 4% paraformaldehyde (Electron Microscopy Sciences, cat#15710) in PBS for 20 min at 4°C, permeabilized with 0.1-0.2% Triton X-100 in PBS for 15-30 min at RT, and processed according to the HCR™ RNA-FISH protocol (Molecular Instruments). For simultaneous detection of *nHOTAIRM1*, validated *nHOTAIRM1*-interacting mRNAs and candidate mediator RNAs, cells were hybridized overnight at 37°C with independent sets of split-initiator probes (50 nM each) and subsequently incubated for 3 h at room temperature with amplifier probes conjugated to 647, 594 or 488 fluorophore. Each candidate mediator RNA was analyzed in an independent experiment together with *nHOTAIRM1* and one validated *nHOTAIRM1*-interacting mRNA (Supplementary Table S2 reports the number of RNA-FISH probes used; sequences are available upon request).

Nuclei were counterstained with DAPI (1 μg/ml, Sigma-Aldrich, cat#D9542) for 5 min and mounted with ProLong™ Diamond Antifade Mountant (Invitrogen, cat#P36961). Images were acquired on an inverted Olympus IX73 confocal microscope equipped with a CrestOptics X-Light V3 spinning disk system, Prime BSI Express Scientific CMOS camera, 100× (NA 1.3) oil objective, Z-step size of 0.2 μm, using MetaMorph software (Molecular Devices).

Post-acquisition processing was performed using FIJI. Following background subtraction, spatial proximity of RNA signals was evaluated on maximum Z-projections using the Spots Colocalization (ComDet) plug-in. *nHOTAIRM1*-mRNA colocalization events were first identified using the same centroid-to-centroid distance threshold applied for double-target RNA-FISH analyses. Triple-colocalization events were subsequently quantified exclusively within neurites as the percentage of candidate mediator RNA puncta colocalizing with *nHOTAIRM1*-mRNA colocalization events.

### CLIP assay

CLIP assay was performed on cytoplasmic extracts of iPSC-derived spMNs (D12) targeting ANXA11. Cells (75 × 10^6) were UV cross-linked at 254 nm (4000×100 μJ/cm²) using a Stratalinker, lysed in NP-40 buffer (50 mM Hepes-KOH pH 7.5, 150 mM KCl, 2 mM EDTA, 1 mM NaF, 0.5% NP-40, 0.5 mM DTT, 1X PIC, RiboLock RNase Inhibitor (Thermo Fisher, cat#EO038SKB011)), and clarified by centrifugation (18,000 × g, 10 min, 4°C). Lysates were incubated overnight at 4°C with Dynabeads™ Protein G (Thermo Fisher, cat#10004D) pre-bound to anti-ANXA11 antibody (10 μg) or IgG control (Santa Cruz Biotechnologies, cat#sc-2025). Beads were washed with high-salt buffer (50 mM Hepes-KOH, 500 mM KCl, 0.5 mM DTT, 0.05% NP-40), treated with Proteinase K (Thermo Fisher, cat#AM2546, 20 mg/ml, 30 min 50°C), and RNA extracted using miRNeasy Micro Kit (Qiagen) with DNase I. Protein fractions were eluted in Laemmli buffer for Western blot validation. RNA pull-down of *nHOTAIRM1* was analyzed by qRT-PCR. Full-length uncropped images of all western blots presented in the main and supplementary figures are provided in the Supplemental Material.

### Native RNA pull-down assay

Native RNA pull-down experiments were performed as described in^60^ with minor modifications (probe sequences are indicated in Supplementary Table S2).

### Native RNA pull-down coupled to ANXA11 protein detection

Native *nHOTAIRM1* RNA pull-down assays were performed as described above. Following RNA capture, bead-bound complexes were washed and divided into RNA and protein fractions. Protein samples were eluted in Laemmli sample buffer, resolved by SDS-PAGE and analyzed by western blot using anti-ANXA11 antibody (ABclonal, cat#A20841). GAPDH (Abcam, cat#ab8245) was used as a negative control. Immunoreactive bands were detected using HRP-conjugated secondary antibodies and WesternBright™ Sirius chemiluminescent substrate (Advansta, cat#K-12043-D20). Full-length uncropped images of all western blots presented in the main and supplementary figures are provided in the Supplemental Material.

### RNA Pull-down sequencing

RNA from *nHOTAIRM1* native RNA pull-down assays, input, and LacZ controls from iPSC-derived spMNs was extracted using miRNeasy Micro Kit (Qiagen). RNA libraries were prepared using the Illumina Stranded Total RNA Prep with Ribo-Zero Plus kit and sequenced on an Illumina NovaSeq 6000 platform in paired-end mode, generating an average of 52.5M of 100 base-long reads. Read quality was assessed with FastQC v0.11.9, which revealed a transient reduction in quality at the first base, residual contamination consistent with incomplete ribosomal RNA removal by Ribo-Zero Plus, and the presence of Illumina adapter sequences. Adapter trimming and removal of low-quality nucleotides were performed using cutadapt v3.2^61^ with: *-u 1 -U 1 --trim-n --nextseq-trim=20 -m 18* and Trimmomatic software v0.39^62^ with PE mode and the following parameters: *ILLUMINACLIP:adapter_path:2:30:10:8:true LEADING:3 TRAILING:3 SLIDINGWINDOW:4:20 MINLEN:18*. To eliminate remaining ribosomal RNA, trimmed reads were aligned to an rRNA-only reference (Table S1) using Bowtie2 v2.4.2,^63^ and only unmapped reads were retained. These filtered reads were subsequently aligned to the GRCh38 human genome using STAR v2.7.7a^64^ with the following parameters: *--outSAMstrandField intronMotif --outSAMattrIHstart 0 --outFilterType BySJout --outFilterMultimapNma× 100 --winAnchorMultimapNma× 100 -- alignSJoverhangMin 8 --alignSJDBoverhangMin 1 --outFilterMismatchNmax 999 -- outFilterMismatchNoverLmax 0.04 --alignIntronMin 20 --alignIntronMa× 1000000 --alignMatesGapMa× 1000000 --outFilterIntronMotifs RemoveNoncanonical --readFilesCommand zcat --peOverlapNbasesMin 50 --outReadsUnmapped Fastx*. Because pull-down-derived RNA libraries typically undergo additional PCR amplification, PCR duplicates were removed using Picard MarkDuplicates v2.24.1(https://broadinstitute.github.io/picard/) and resulted BAM files were further filtered to retain only alignments from properly paired reads using samtools v1.7^65^ with *view -f 2* parameters. Gene-level quantifications were performed with HTSeq-count v0.13.5 using Ensembl GTF gene annotation related to release 99^66^ and these parameters: *-s reverse -m union -t exon*. Read counts at each processing stage are reported in the accompanying summary (Table S1).

Genes were filtered to retain those with at least 10 raw counts in at least 2 samples. Filtered counts were used to construct an edgeR v3.34.1 (42) DGEList object. Library-size normalization factors were computed using calcNormFactors(“none”) imposing no between-sample normalization. Normalized expression matrices were generated as CPM using edgeR::cpm and edgeR::rpkm. gene lengths used to calculate FPKMs were computed from the Ensembl release 99 GTF as the total length of the union of all exonic intervals for each gene locus. Sample-to-sample similarity was assessed by computing a Pearson correlation matrix on CPM values, which was visualized using corrplot with hierarchical clustering order. For differential expression testing, a generalized linear model (GLM) framework was applied using edgeR. A design matrix was built with a blocking factor corresponding to biological replicates *(∼0 + condition + replicates*). Dispersion parameters were estimated with estimateDisp function with robust=TRUE parameter to reduce the influence of potential outliers. The negative binomial GLM was fitted with glmFit, and differential expression was assessed using likelihood ratio tests (glmLRT) for: the contrasts: EVEN vs INPUT or ODD vs INPUT; and a difference-of-differences (interaction-like) contrast defined as (EVEN − INPUT) − (LACZ − INPUT) or (ODD − INPUT) − (LACZ − INPUT) to calculate the enrichment specificity in *nHOTAIRM1* RNA pull-down compared to LACZ control.

### AMT-crosslinked RNA pull-down assay

AMT-crosslinked RNA pull-down experiments were performed as described in^17^ (probe sequences are indicated in Table S2).

### RNA-protein interaction prediction

Prediction of *nHOTAIRM1* and ANXA11 (UniProtKB P50995-1) interaction was performed using *cat*RAPID omics v2.1 web server. Interaction propensity scores were computed using full-length ANXA11 and *nHOTAIRM1* sequences. Interaction maps were generated according to the default server settings.

### RNA-RNA interaction prediction

The sequences of the transcripts recovered in the *nHOTAIRM1* RNA pull-down were retrieved from Ensembl release 99^67^ (Homo sapiens gene annotation, biomaRt^68^), together with the neuronal isoform of *nHOTAIRM1* (1,068 nt). To increase prediction robustness and minimize method-specific biases, all-versus-all pairwise RNA-RNA interactions were inferred using three independent predictors relying on complementary strategies: IntaRNA v2.3^69^ with the -n 10 parameter; RIsearch2 v1.2^70^; and RIME^71^, run in RIMEfull mode. Self-interactions were discarded from all predictions.

To retain only high-confidence duplexes, predictions were required to be supported by all three methods. Each IntaRNA duplex, encoded in BEDPE format, was retained only if both of its interacting windows reciprocally overlapped with at least one RIME window pair (RIME score > 0.50) and at least one RIsearch2 duplex predicted for the same transcript pair. Overlaps were calculated with BEDTools v2.29.1^72^ using pairtopair -type both parameters. For each transcript pair, the duplex with the lowest hybridization free energy in this consensus set was retained as the representative interaction.

Candidate mediators were defined as transcripts able to engage an RNA pull-down target at a site distinct from the one occupied by *nHOTAIRM1*, so that the resulting triplet is sterically compatible. For each target transcript T, the region contacted by *HOTAIRM1* was defined as the T-side window of the representative *nHOTAIRM1*-T duplex. All IntaRNA duplexes involving T and any transcript other than *nHOTAIRM1* were then intersected with this region using BEDTools v2.29.1 with pairtobed -type neither parameter and only duplexes falling entirely outside the *HOTAIRM1*-contacted window were retained. The resulting set was subjected to the same three-method consensus filter described above and reduced to the minimum-energy duplex per transcript pair; representative *nHOTAIRM1* duplexes were re-oriented and added back to obtain the final interaction network.

From this network, a transcript-by-transcript matrix of minimum hybridization free energies was assembled. For every candidate transcript M and every target T, we computed the differential binding energy ΔΔG = ΔG(T vs *nHOTAIRM1*) − ΔG(T vs M), so that positive values indicate a stronger predicted affinity of T for the mediator than for *nHOTAIRM1* and values close to zero indicate comparable engagement of both partners.

### DNA/LNA steric-blocking competition assay

To functionally interfere with the predicted RNA-RNA interaction interfaces between *nHOTAIRM1* and its target mRNAs without inducing transcript degradation, iPSC-derived spinal motor neurons (spMNs) were transfected with a pool of four non-degradative DNA/LNA steric-blocking mixmers (Eurogentec). Each 20-nt mixmer was fully complementary to a distinct computationally predicted *nHOTAIRM1* interaction region involved in RNA-RNA pairing with *TPM1*, *YKT6* or *SPTBN1* transcripts. To promote steric blockade while preserving target RNA stability, locked nucleic acid (LNA) residues were introduced at regular intervals (one LNA residue every three DNA nucleotides), following the steric-blocking strategy described by^73^. A non-targeting DNA/LNA steric-blocking mixmer with no predicted complementarity to *nHOTAIRM1* was used as scramble control. Competitor sequences are reported in Supplementary Table S2.

At day 10 of differentiation, the culture medium was replaced with Opti-MEM (Thermo Fisher, cat#31985062) for 1 h at 37°C. Cells were then transfected with either the pool of DNA/LNA steric-blocking mixmers (25 nM each; final concentration 100 nM) or the scramble control (100 nM final concentration) using SilentFect™ Lipid Reagent (Bio-Rad Laboratories, cat#1703360), according to the manufacturer’s instructions. After 5 h, the transfection medium was replaced with complete motor neuron differentiation medium lacking penicillin/streptomycin, and cells were maintained under standard differentiation conditions for an additional 48 h.

At day 12 of differentiation, cells were harvested and processed for native *nHOTAIRM1* RNA pull-down as described above. RNA recovered from RNA pull-down fractions and total cellular RNA was analyzed by qRT-PCR to quantify the enrichment and expression levels of *TPM1*, *YKT6* and *SPTBN1* relative to scramble-transfected cells.

### ANXA11 knockdown

ANXA11 knockdown was performed in iPSC-derived spMNs using an equimolar mixture of two independent Silencer™ Pre-designed siRNAs targeting distinct regions of the human ANXA11 transcript (Invitrogen, cat#AM16708; Assay IDs 147070 and 147071). A Silencer™ Negative Control siRNA (Invitrogen) was used as scramble control.

At day 10 of differentiation, the culture medium was replaced with Opti-MEM (Thermo Fisher, cat#31985062) for 1 h at 37°C. Cells were then transfected with either the ANXA11-targeting siRNA mixture (Silencer™ Pre-designed siRNA, Thermo Fisher Scientific, Assay ID147070 and Assay ID147071, cat#AM16708) (25 nM each; final concentration 50 nM) or the scramble siRNA (Silencer™ Negative Control No. 1 siRNA (Invitrogen, cat#AM4611) (final concentration 50 nM) using SilentFect™ Lipid Reagent (Bio-Rad Laboratories, cat#1703360), according to the manufacturer’s instructions. After 5 h, the transfection medium was replaced with complete motor neuron differentiation medium lacking penicillin/streptomycin, and cells were maintained under standard differentiation conditions for an additional 48 h.

At day 12 of differentiation, cells were harvested either for RNA and protein extraction or fixed with 4% paraformaldehyde for RNA fluorescence in situ hybridization (RNA-FISH), as described above. Knockdown efficiency was assessed by qRT-PCR and western blot analysis. Full-length uncropped images of all western blots presented in the main and supplementary figures are provided in the Supplemental Material.

### RNA extraction and analysis

Total RNA from iPSC-derived spMNs was extracted using Direct-zol RNA MiniPrep (Zymo Research, cat#R2052). Alternatively, miRNeasy Micro Kit (Qiagen, cat#217084) was used. For RNA pull-downs, miRNeasy Micro Kit was preferred. cDNA from total RNA (100-1000 ng) of iPSC-derived spMNs was synthesized using Takara PrimeScript RT Reagent Kit (Takara Bio, cat#RR037A). Alternatively, SuperScript VILO cDNA Synthesis Kit (Thermo Fisher, cat#11754-250) was used. qRT-PCR on cDNA from iPSC-derived spMNs was performed using SensiFAST SYBR Lo-Rox Kit (Bioline, cat#BIO-94020) on Quant Studio 3 Real-Time PCR System (Applied Biosystems). Primers are indicated in Table S2.

### Western blot assay

iPSC-derived spMNs were lysed in RIPA buffer (50 mM Tris-HCl pH 7.5, 150 mM NaCl, 1% NP-40, 0.5% sodium deoxycholate, 0.1% SDS, 1X cOmplete™ Protease Inhibitor Cocktail (Roche, cat#11873580001)) on ice 15-30 min, centrifuged (12,000 × g, 10 min, 4°C). Protein quantified by Bradford assay (Biorad). 20-50 μg loaded on 4-12% Bis-Tris NuPAGE gels (Thermo Fisher), transferred to nitrocellulose (Cytiva cat#10600002). Membranes were blocked in 5% milk/TBST for 1 h at RT. Primary antibodies (anti-ANXA11 1:1000, anti-GAPDH 1:5000) were incubated overnight 4°C, TBST-washed, and subject to HRP-secondary (Goat anti-Rabbit IgG (H+L) Secondary Antibody HRP cat#31460 and Goat anti-Mouse IgG (H+L) Secondary Antibody HRP cat#32430) 1:10000 for 1 h at RT. Signals were detected by WesternBright Sirius Chemiluminescent Detection Kit (Advansta, cat#K-12043-D20). Bands were quantified using Image Lab (BioRad Software). Full-length uncropped images of all western blots presented in the main and supplementary figures are provided in the Supplemental Material.

### Statistical analyses

Histograms show the mean□±□SEM of at least three independent biological replicates. N is indicated in Figure Legends. Errors were calculated from relative quantities and then appropriately propagated; statistical significance was determined using one-sample, paired, or unpaired two-tailed Student’s t tests, as indicated in the corresponding Figure Legends. A p-value□<□0.05 was considered statistically significant. *P□<□0.05, **P□<□0.01, ***P□<□0.001.

## Data availability

The RNA-Seq data presented in this study are available in GEO, accession number <u>GSE319005</u>. To review GEO accession number <u>GSE319005</u>, please visit and enter the token cnyjsyysfbahlmr. Raw data are available upon request.

## Acknowledgements

We thank A. Grandioso for explorative experiments, M. Marchioni for technical support and M. Caruso for assistance. We acknowledge the Imaging Facility of CLN²S@Sapienza, Istituto Italiano di Tecnologia, Rome, the support of PhD Valeria de Turris for confocal microscopy acquisitions, and the Microscopy Facility (part of Sapienza Research Infrastructure) of the Department of Biology and Biotechnologies “C. Darwin” for technical support in image acquisition. We are also grateful to Dr. Diego Vozzi and the Genomic Facility of IIT for support in RNA sequencing experiments and to the members of the IIT RNA Technologies Flagship for thought-provoking discussion.

## Funding

This work was supported by: (1) ERC-2019-SyG (855923-ASTRA) to IB; (2) European Union - Next Generation EU and MUR, NRRP - M4C2 - Action 1.4, Project “National Center for Gene Therapy and Drugs based on RNA Technology”, no. CN00000041, Spoke 3 “Neurodegeneration” to MB and IB, and (3) Spoke #6 “RNA Drug Development” to PL; (4) CNR of Italy projects (id. DBA.AD005.225-NUTRAGE-FOE2021 and DSB.AD006.371-InvAtt-FOE2022) to PL; (5) MUR PRIN 2017 (id. 2017P352Z4) to PL; (6) European Union - Next-Generation EU and MUR, NRRP - M4C2 - Action 1.1, Call “PRIN 2022” (id. 2022BYB33L) to MB and PL.

## Author contributions

PT: conceptualization, investigation (cell culture, MN differentiation, gene expression analysis, cloning, molecular and cellular assays, RNA-protein and RNA-RNA interactomics, imaging experiments, confocal microscopy acquisition and post-acquisition analysis), formal analysis, validation, methodology, data curation, visualization; writing-original draft manuscript. AS: bioinformatic analysis, formal analysis, data curation, software. JR and DM: RNA-seq library preparation and sequencing. WV: investigation (gene expression analysis, formal analysis), validation. JR: methodology, data curation. TS: methodology (imaging), data curation and post-acquisition analysis. AB: validation, data curation, visualization. MB: methodology, validation, resources, data interpretation. IB and PL: conceptualization, supervision, writing-original draft (main text); funding acquisition, project administration. PT, MB, PL: Writing - Review & Editing. All authors approved the final version of the manuscript.

## Conflict of interest

The authors declare no conflict of interests.

## Ethics

All methods were performed in accordance with the relevant guidelines and regulations. This study did not involve human participants, human personal data, or animals.

## Informed consent

The manuscript is approved by all authors for publication

Data represent three independent biological replicates unless otherwise indicated.

## Notes

https://www.ncbi.nlm.nih.gov/geo/query/acc.cgi?acc=GSE319005

### Competing Interest Statement

The authors have declared no competing interest.

