## Supplemental Material for "*nHOTAIRM1* scaffolds ANXA11-dependent assemblies to drive axonal mRNA localization in human motor neurons"

**Supplemental Material – Original uncropped western blots**

This file contains the full-length uncropped western blot images corresponding to the immunoblots presented in Figure 1A, Supplementary Figure 2B and Supplementary Figure 16A.
All membranes are shown prior to cropping for figure assembly.
Molecular weight markers and all detected bands are displayed.

The corresponding figure panels are indicated above each blot.

### Figure 1A - ANXA11 CLIP Immunoblot

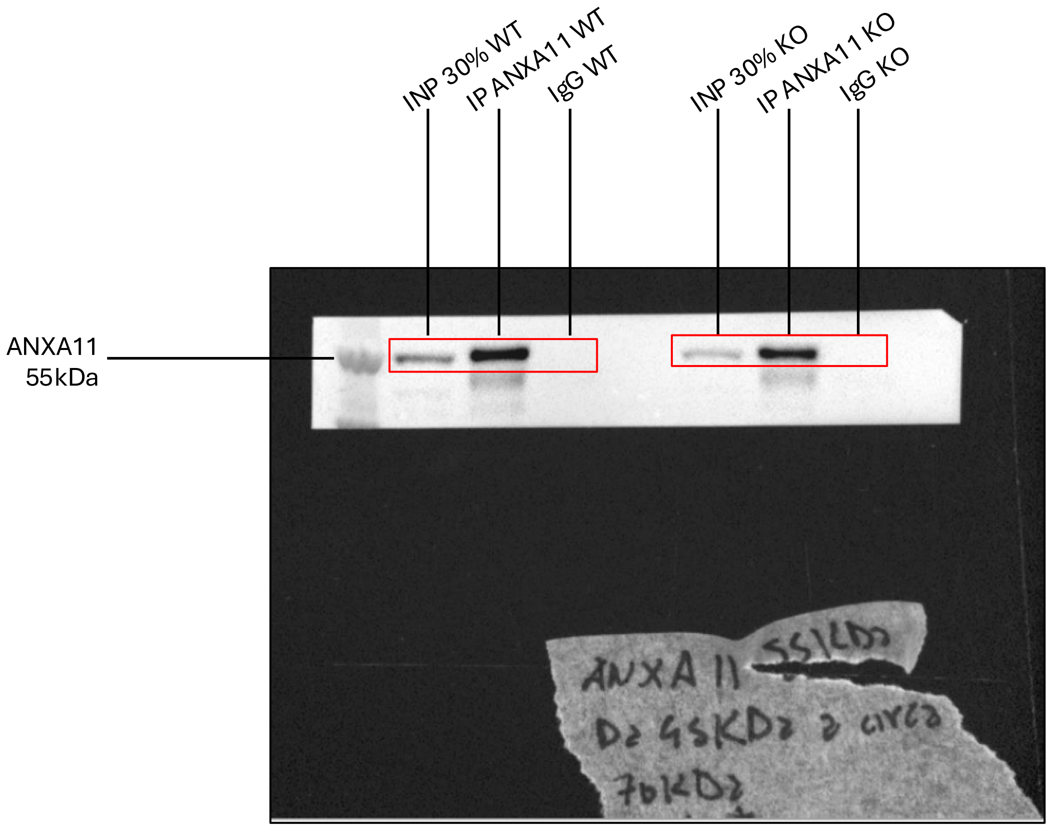


Membrane probed with anti-ANXA11 antibody.
Expected molecular weight: ~55 kDa.
Red box indicates the cropped region shown in Figure 1A of the manuscript.

**Supplementary Figure 2B – *nHOTAIRM1* RNA pull-down Immunoblot**

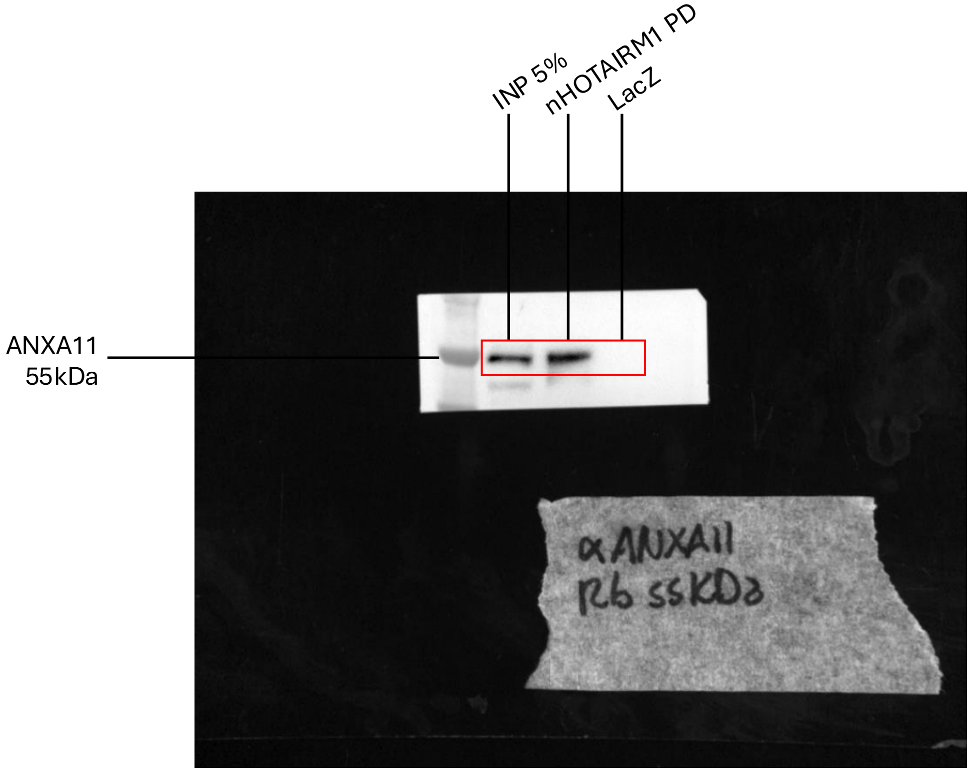


Membrane probed with anti-ANXA11 antibody.
Expected molecular weight: ~55 kDa.
Red box indicates the cropped region shown in Supplementary Figure 2B of the manuscript.

### Supplementary Figure 16A – ANXA11 knockdown Immunoblot

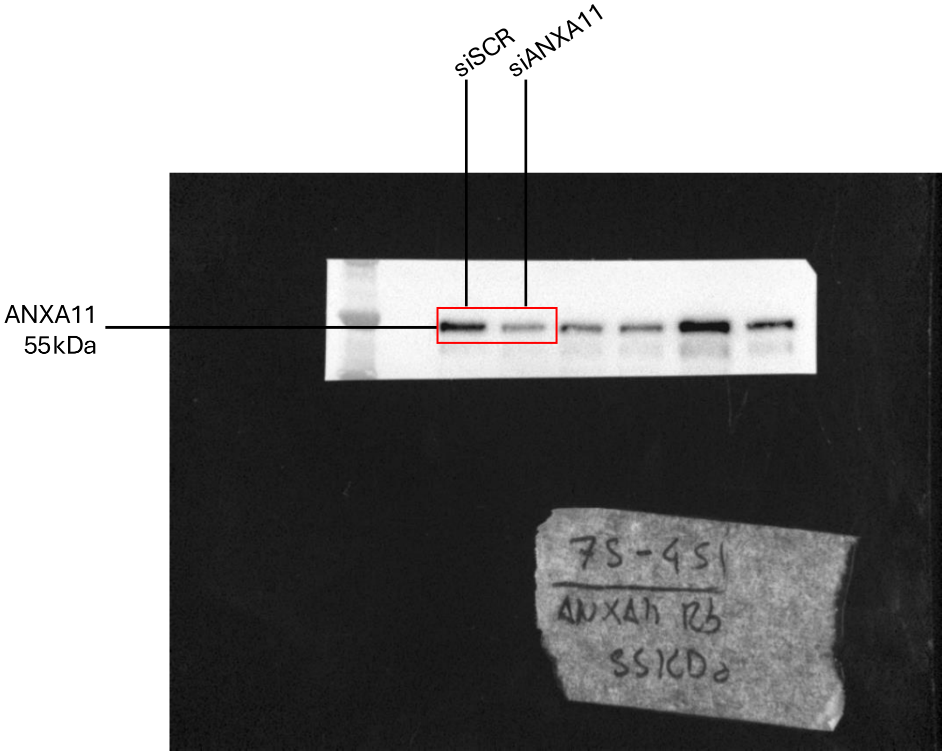


Membrane probed with anti-ANXA11 antibody.
Expected molecular weight: ~55 kDa.
Red box indicates the cropped region shown in Supplementary Figure 16A of the manuscript.

#### Supplementary Figure 16A – ANXA11 knockdown Immunoblot

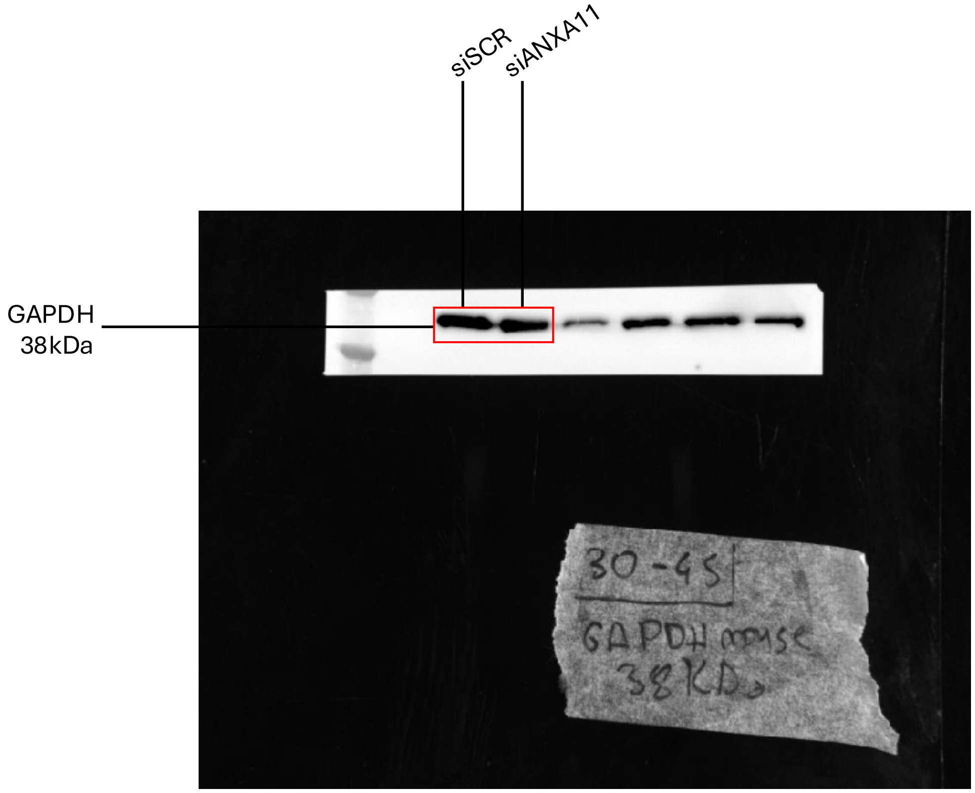


Membrane probed with anti-GAPDH antibody.
Expected molecular weight: ~38 kDa.
Red box indicates the cropped region shown in Supplementary Figure 16A of the manuscript.
