## Supplementary Information for "*nHOTAIRM1* scaffolds ANXA11-dependent assemblies to drive axonal mRNA localization in human motor neurons"

**Contents:**
Supplementary Figures S1-S20
Supplementary Figure Legends

### Supplementary Figure 1

**A**

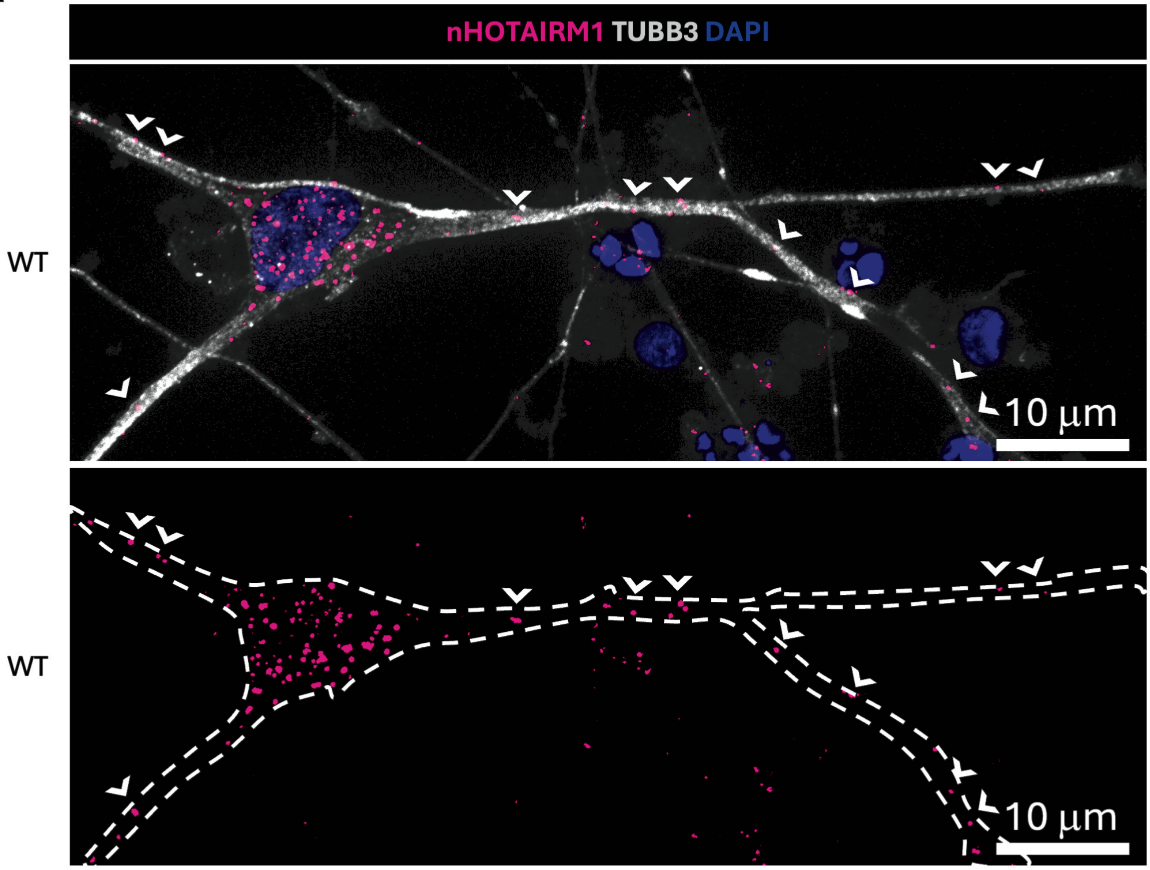


**B**

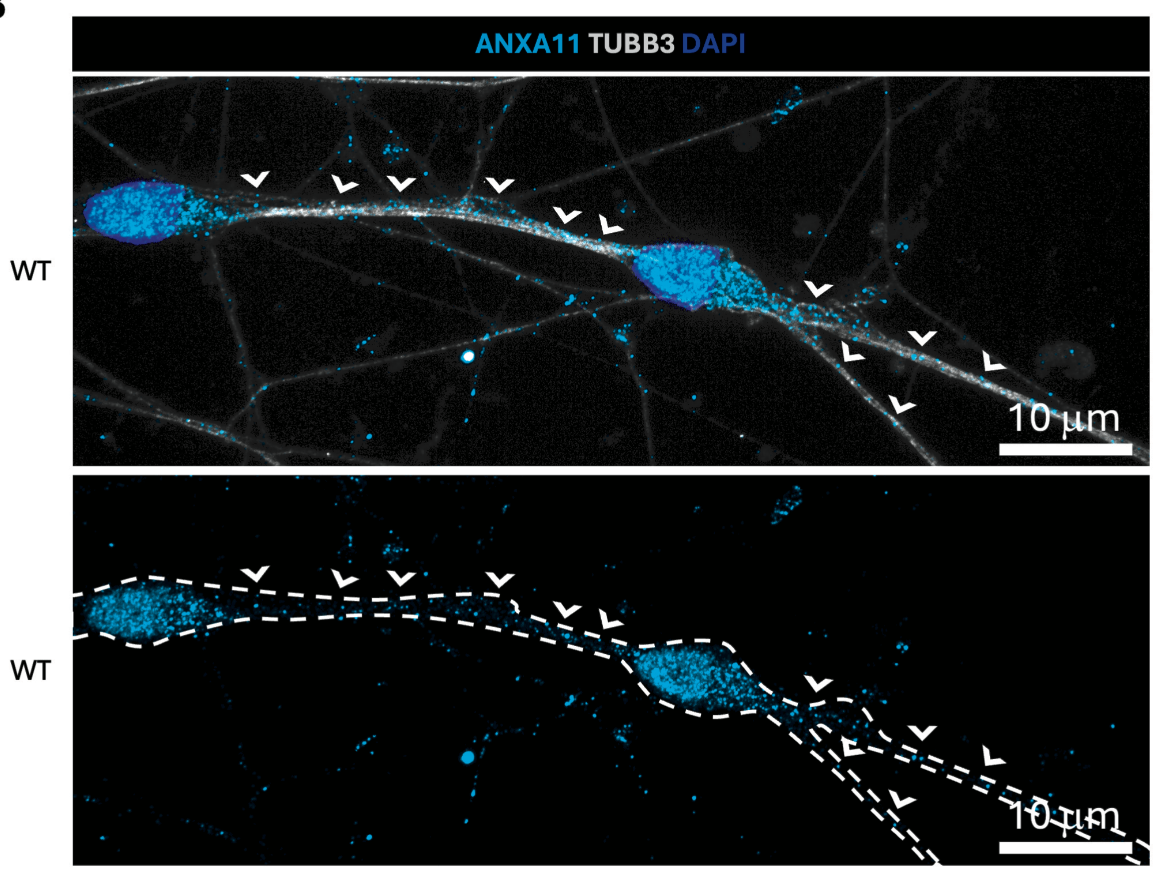

**Supplementary Figure 1. *nHOTAIRM1* localization in iPSC-derived spMNs**

(A) *nHOTAIRM1* localization. Representative images of RNA fluorescence in situ hybridization (RNA-FISH) for *nHOTAIRM1* (magenta), combined with immunofluorescence (IF) for the neuronal marker TUBB3 (gray) and DAPI nuclear staining (blue) in wild-type (WT) spMNs at day 12 of differentiation. Upper panel: merged image showing the co-distribution of signals; arrowheads indicate *nHOTAIRM1* RNA FISH spots localized within both the soma and the neurites. Lower panel: single-channel visualization of *nHOTAIRM1* (magenta); the dashed line outlines the neuronal morphology (based on TUBB3 signal) to highlight the extra-nuclear distribution of the RNA. Scale bars: 10 μm.

(B) ANXA11 localization. Representative IF for ANXA11 (cyan), TUBB3 (gray), and DAPI (blue) in WT spMNs at day 12 of differentiation. Upper panel: merged image showing overall protein distribution; arrowheads point to ANXA11-positive puncta along the neuronal processes. Lower panel: single-channel visualization of ANXA11 (cyan); the dashed line delineates the cell body and neurite boundaries to emphasize the protein's localization. Scale bars: 10 μm.

### Supplementary Figure 2

**A**

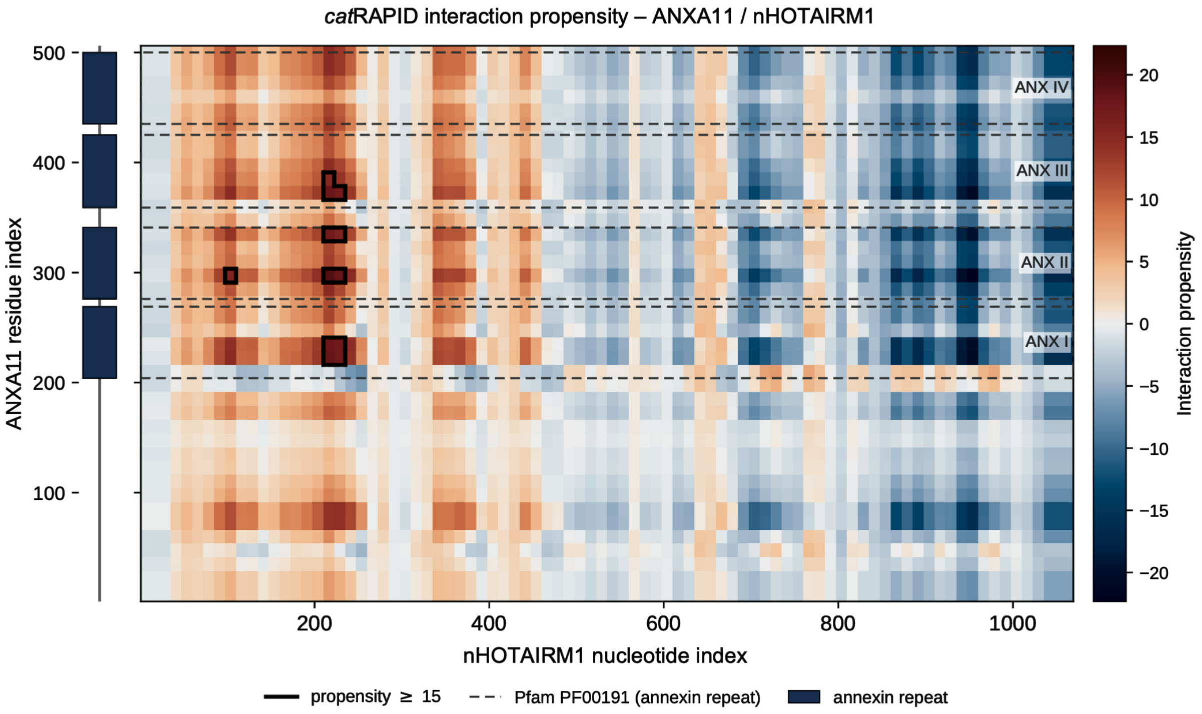


**B**

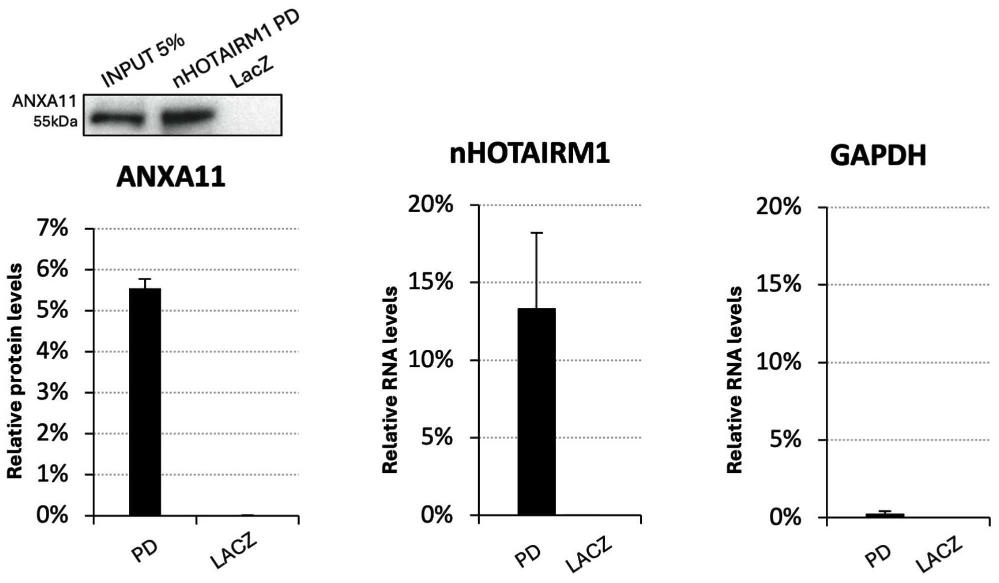

**Supplementary Figure 2. Computational and biochemical analysis of the *nHOTAIRM1*-ANXA11 interaction**

(A) catRAPID prediction of the interaction between the neuronal isoform of *nHOTAIRM1* and full-length human ANXA11. The heatmap shows interaction-propensity scores for all ANXA11-*nHOTAIRM1* fragment pairs, ranging from negative values (blue) to positive values (red). Black outlines indicate fragment pairs with interaction propensity ≥15. Dashed boxes indicate the RNA-binding annexin-repeat domains of ANXA11 corresponding to Pfam domain PF00191, and blue rectangles schematically indicate the same repeats along the protein sequence.

(B) Native *nHOTAIRM1* RNA pull-down followed by ANXA11 protein detection. Representative immunoblot shows ANXA11 enrichment in the *nHOTAIRM1* pull-down fraction relative to the LacZ control. Histograms show ANXA11 protein enrichment and qRT-PCR analysis of *nHOTAIRM1* and GAPDH RNA recovery in *nHOTAIRM1* pull-down and LacZ fractions. GAPDH was used as a negative control transcript. Data are presented as mean ± SEM from two independent biological replicates ($N = 2$).

### Supplementary Figure 3

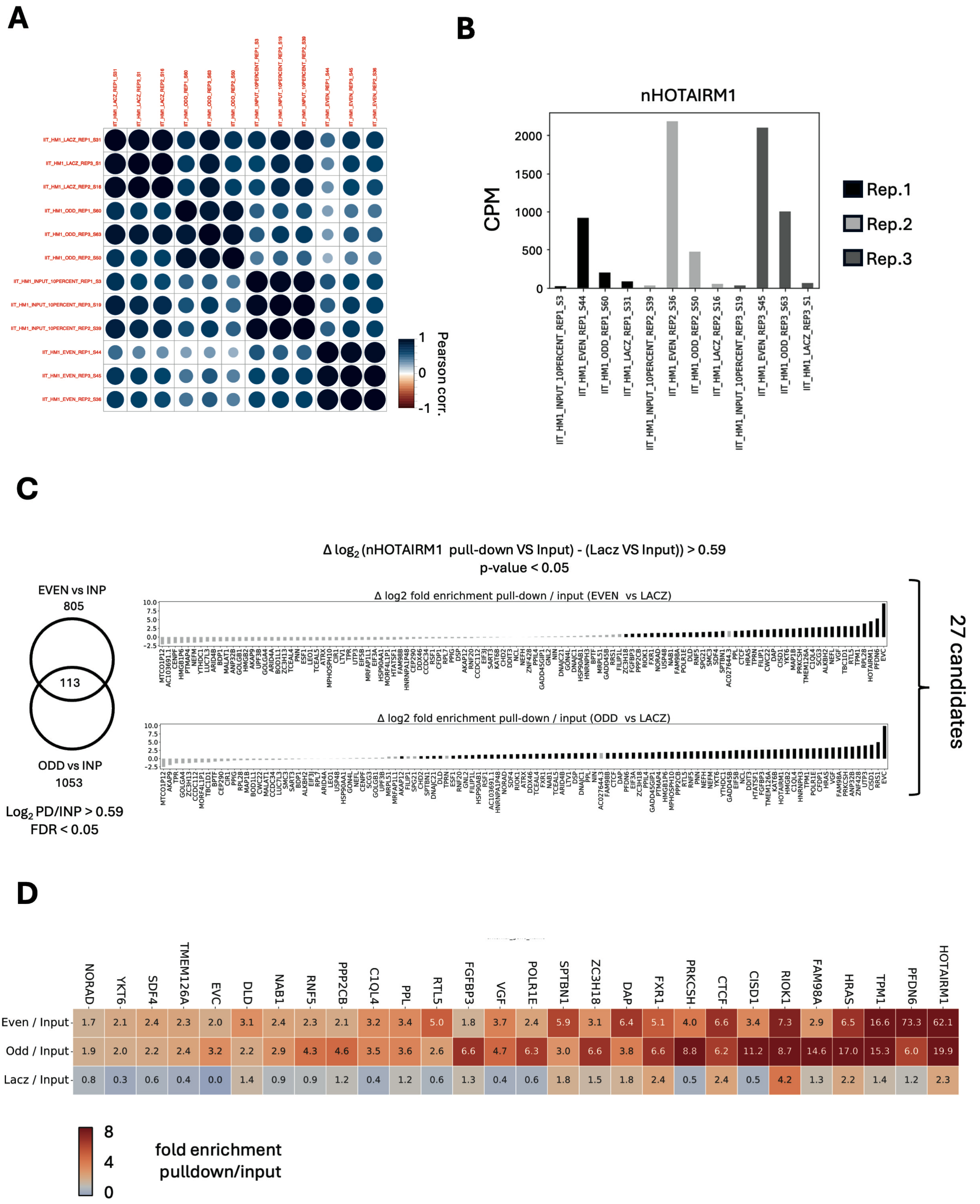

**Supplementary Figure 3. RNA-seq analysis of *nHOTAIRM1* native RNA pulldown in iPSC-derived spMNs**

(A) Sample-to-sample correlation matrix across input (INP) and pull-down (PD) libraries. Circle size and color represent Pearson correlation coefficients calculated on normalized gene expression values (CPM).

(B) *nHOTAIRM1* abundance (CPM) across input and pull-down samples.

(C) Identification of *nHOTAIRM1*-associated transcripts. Left: Venn diagram of genes significantly enriched in pull-down versus input (PD/INP; log2 fold-change > 0.59 and FDR < 0.05) using EVEN or ODD probe sets, with overlap indicating high-confidence candidates detected by both probe pools. Right: ranked bar plots showing Δlog2 fold enrichment relative to LacZ control ((EVEN or ODD pull-down vs input) - (LacZ vs input)); the 27 candidates meet threshold of Δlog2 fold-change > 0.59 and FDR < 0.05.

(D) Heatmap of selected candidate transcripts showing enrichment in *nHOTAIRM1* pull-down fractions. Values represent log2 fold-change (pull-down/input) for EVEN and ODD probe sets compared with LacZ control.

### Supplementary Figure 4

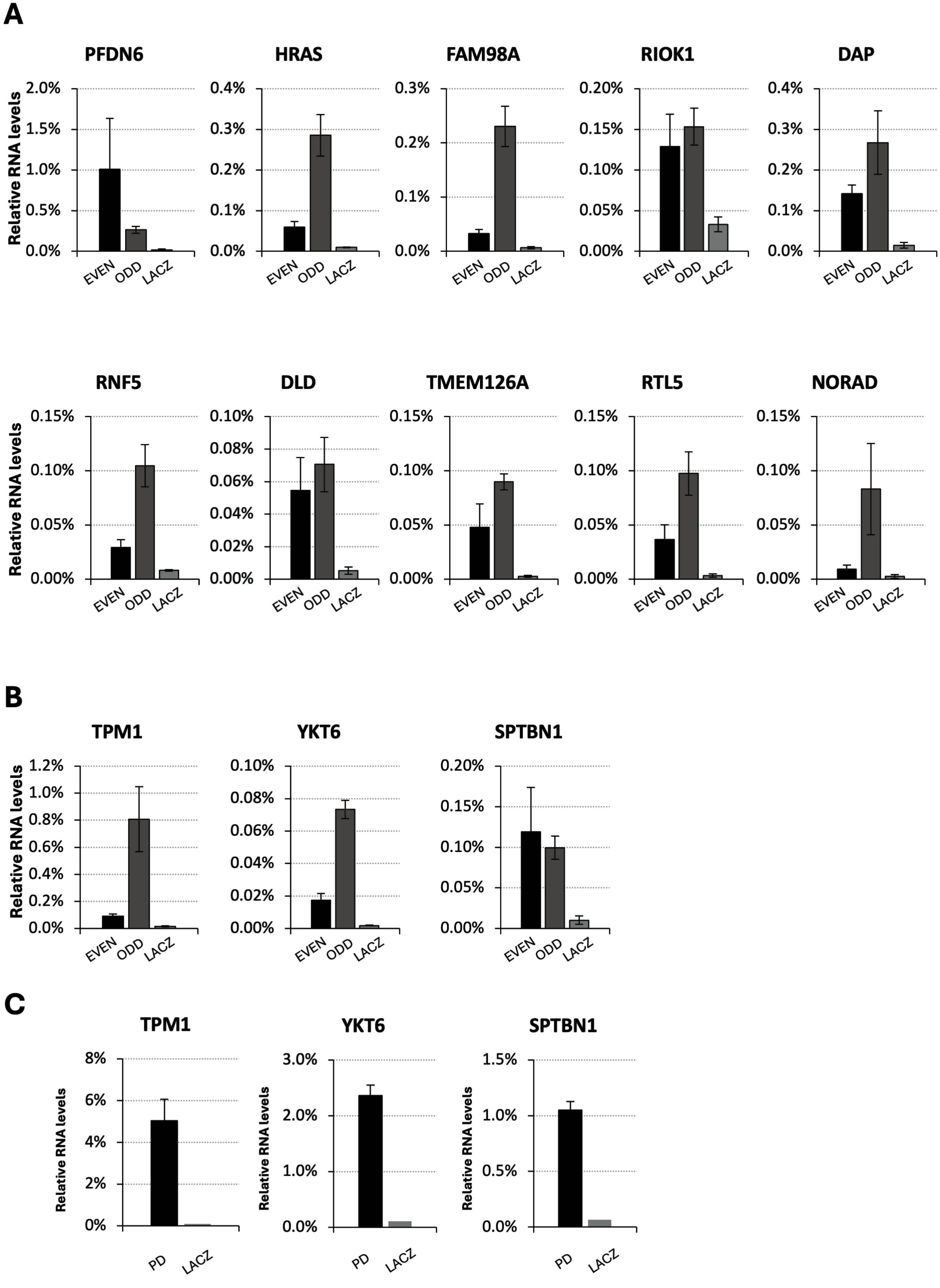

**Supplementary Figure 4. Validation of candidate *nHOTAIRM1* RNA interactors**

(A) qRT-PCR analysis of RNA enrichment for 10 candidate transcripts selected from native *nHOTAIRM1* RNA pull-down sequencing. RNA levels in EVEN, ODD and LacZ fractions were compared to input. Data are expressed as percentage of input. *N* = 3 independent biological replicates.

(B) qRT-PCR analysis of RNA enrichment for *TPM1*, *YKT6* and *SPTBN1* mRNA interactors selected from AMT-crosslinked *nHOTAIRM1* RNA pull-down assays. RNA levels in EVEN, ODD and LacZ fractions were compared to input. Data are expressed as percentage of input. *N* = 3 independent biological replicates.

Supplementary Figure 5

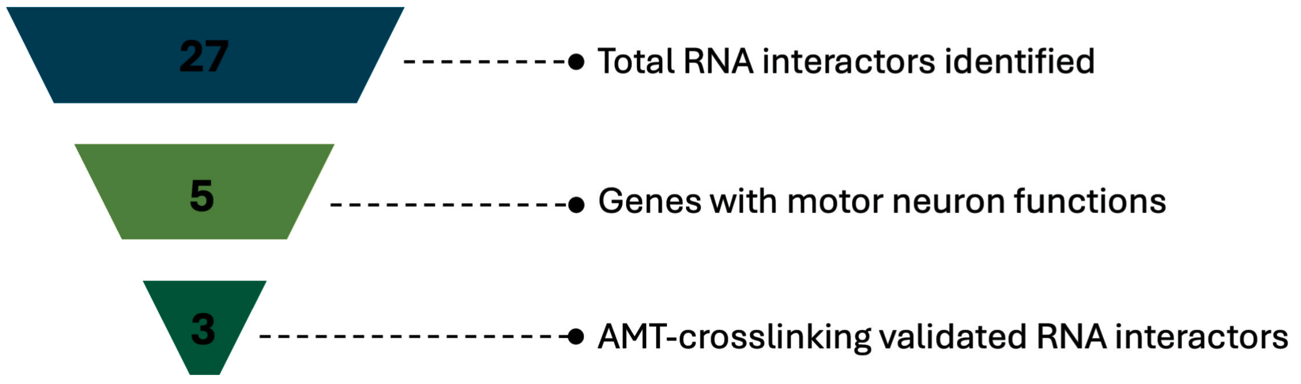

**Supplementary Figure 5. Selection pipeline of high-confidence *nHOTAIRM1* RNA interactors**
Schematic representation of the prioritization strategy used to identify high-confidence *nHOTAIRM1* RNA interactors from native RNA pull-down sequencing data. A total of 27 transcripts were initially identified as significantly enriched in *nHOTAIRM1* pull-down fractions compared to input and LacZ controls. Among these, 5 genes were annotated as having established roles in MN biology based on functional annotation and literature curation. Finally, 3 transcripts were further validated by AMT-crosslinked RNA pull-down assays, supporting direct RNA-RNA interaction with *nHOTAIRM1*. The funnel diagram illustrates the progressive refinement of candidate interactors through functional filtering and experimental validation.

### Supplementary Figure 6

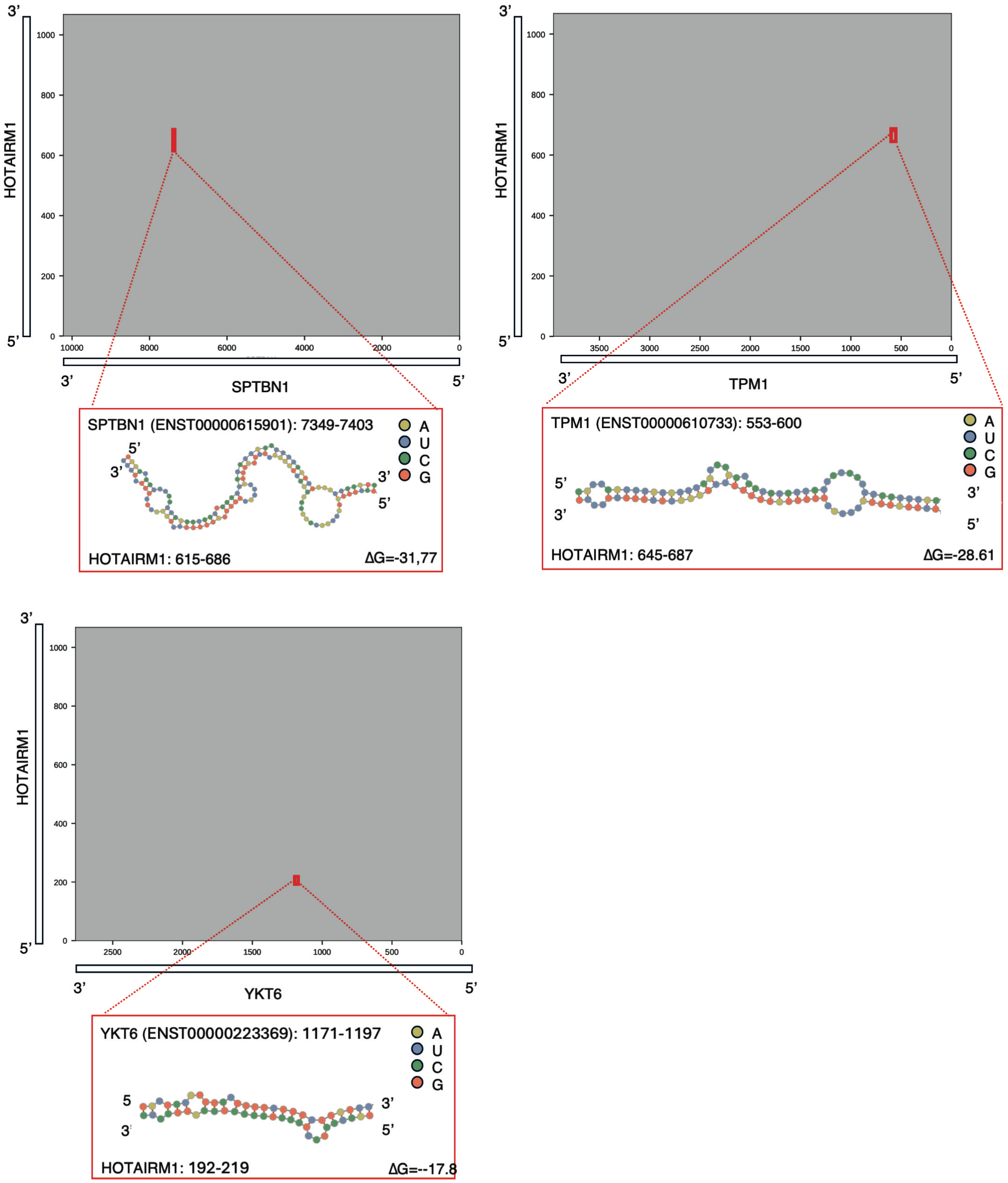

**Supplementary Figure 6. Predicted RNA-RNA duplexes between *nHOTAIRM1* and its validated mRNA partners**

Predicted duplexes between *nHOTAIRM1* and *SPTBN1* (A), *TPM1* (B) and *YKT6* (C). Interaction maps show the coordinate space of each transcript pair, with *nHOTAIRM1* plotted on the y axis in the 5'-3' direction and the target transcript plotted on the x axis in the 3'-5' direction. Red boxes indicate the minimum-free-energy interaction retained from the consensus prediction generated using IntaRNA, RIsearch2 and RIME. Insets show the corresponding hybrid structures, with nucleotides colored according to base identity and interacting intervals reported as transcript coordinates for the target transcript and *nHOTAIRM1*. ΔG values indicate predicted hybridization free energy in kcal $mol^{-1}$. Hybrid structures were visualized using the Forna web tool.

### Supplementary Figure 7

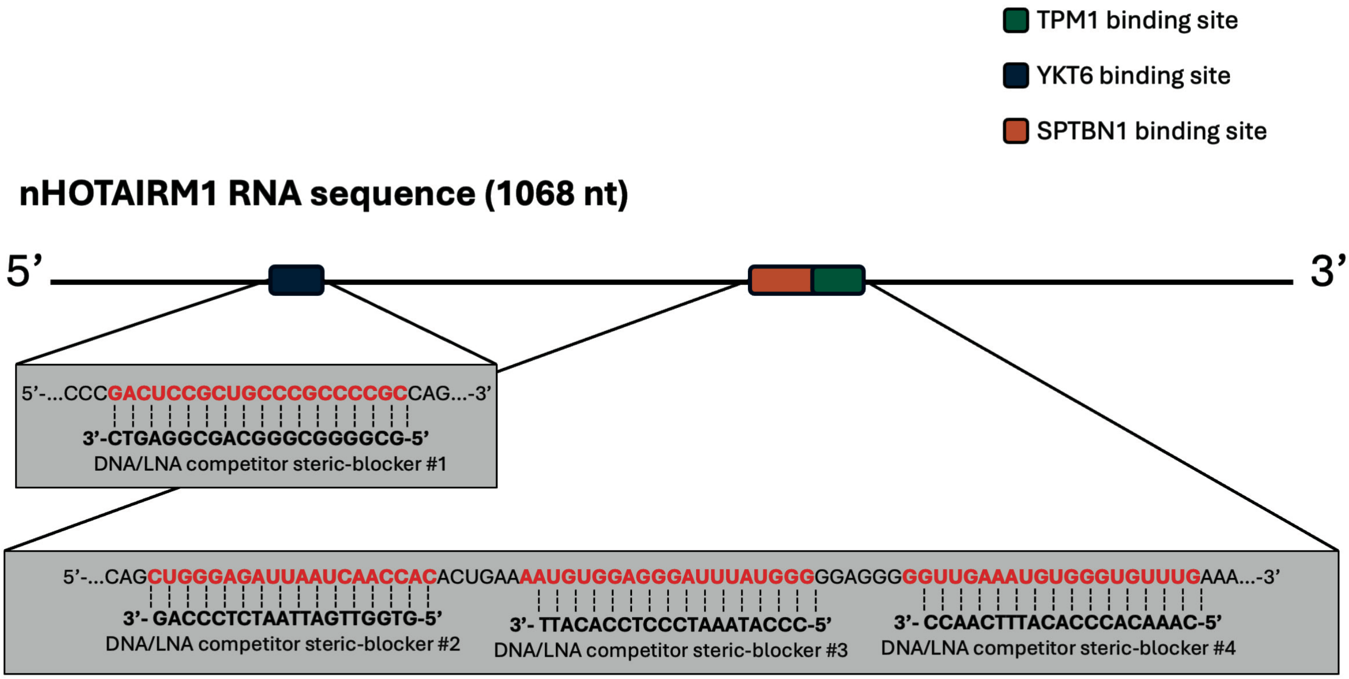


| COMPETITOR | SEQUENCE |
|---|---|
| DNA/LNA competitor steric-blocker #1 | GCGGGGCGGGCAGCGGAGTC |
| DNA/LNA competitor steric-blocker #2 | GTGGTTGATTAATCTCCCAG |
| DNA/LNA competitor steric-blocker #3 | CCCATAAATCCCTCCACATT |
| DNA/LNA competitor steric-blocker #4 | CAAACACCCACATTTCAACC |

**Supplementary Figure 7. Design of DNA/LNA steric-blocking competitors targeting predicted *nHOTAIRM1* interaction regions**

Schematic representation of the neuronal *nHOTAIRM1* transcript and the predicted binding regions for *TPM1*, *YKT6* and *SPTBN1*. Enlarged sequence views show the complementary binding sites targeted by the four non-degradative DNA/LNA steric-blocking competitors. The table reports the sequence of each competitor used in the competition assay.

Supplementary Figure 8

**A** SOMA

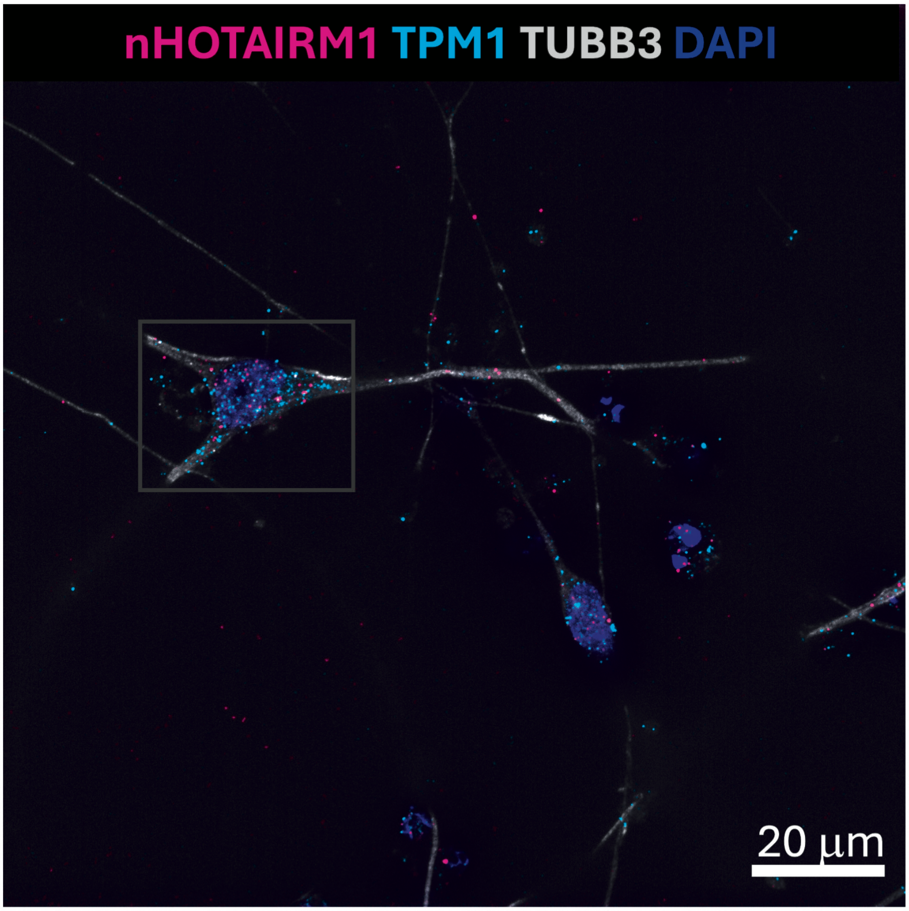


WT

**B** NEURITES

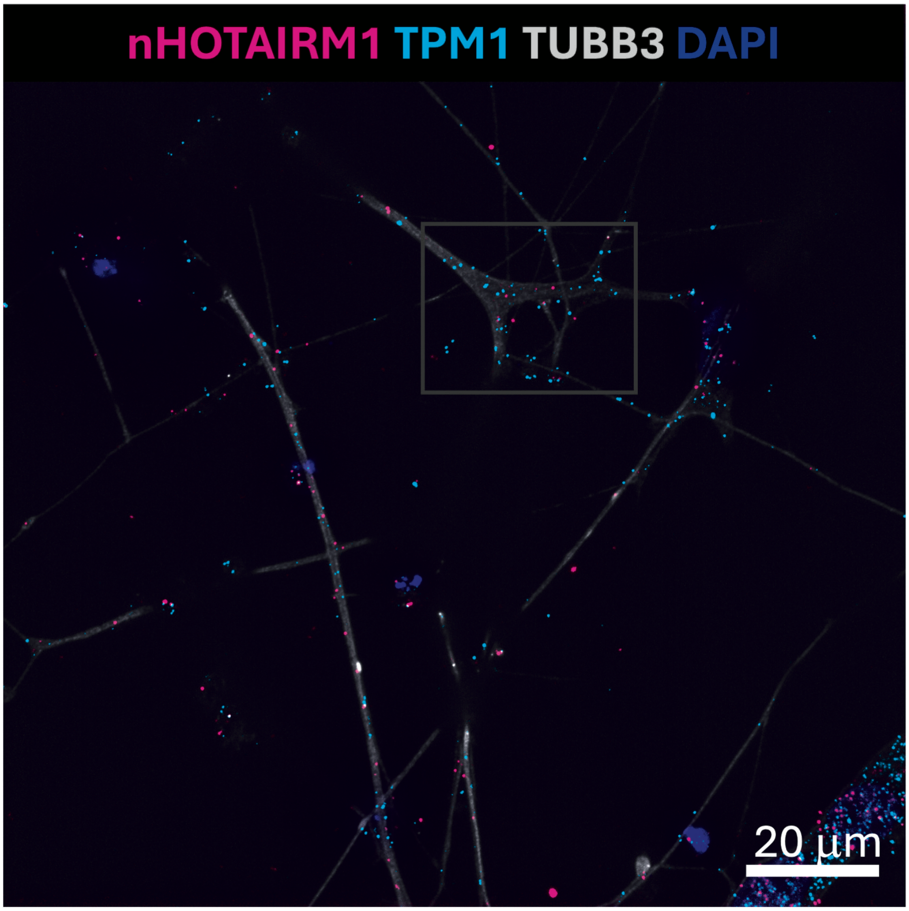


WT

**Supplementary Figure 8. Confocal imaging of *nHOTAIRM1* and *TPM1* mRNA in spMNs (related to main Fig. 4)**

Representative confocal Z-stack acquisition of spMN (A) soma and (B) neurites showing RNA-FISH for *nHOTAIRM1* (magenta) and *TPM1* mRNA (cyan), IF for TUBB3 (gray), and DAPI (blue). Images were acquired at 100x magnification using confocal microscopy with a Z-step size of 0.2 µm. The digital enlargements and 3D renderings shown in Figure 4 are derived from selected regions of these acquisitions. Scale bars: 20 µm.

Supplementary Figure 9

**A**

SOMA

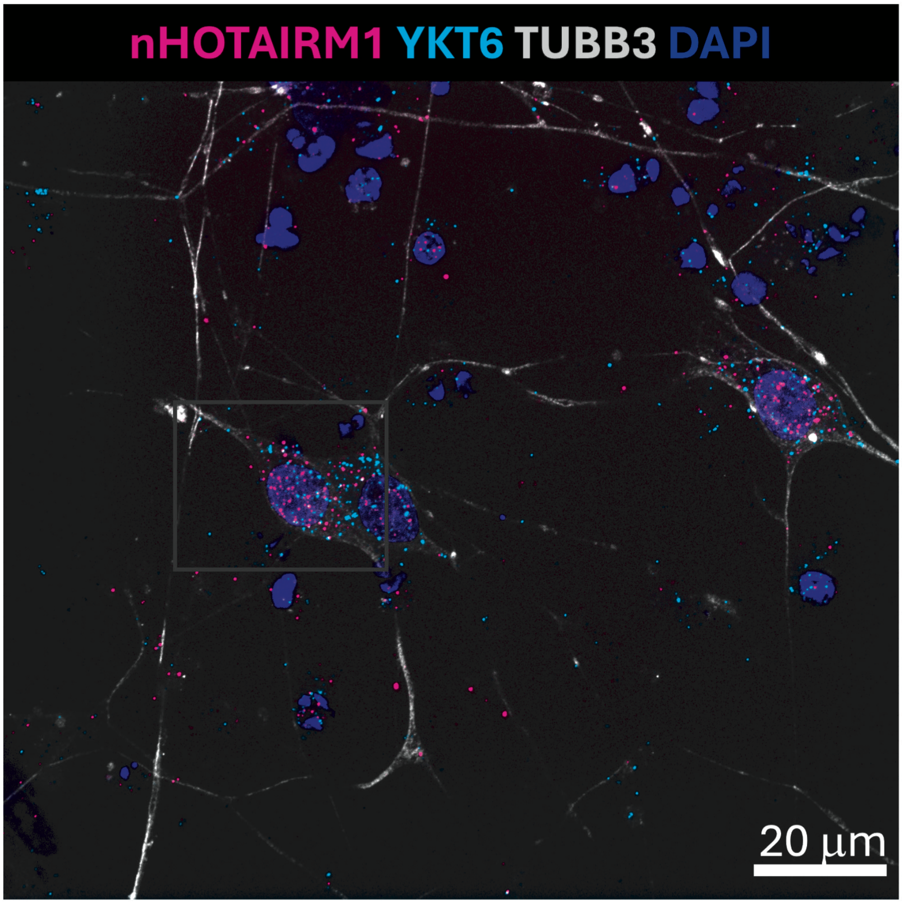


WT

**B**

NEURITES

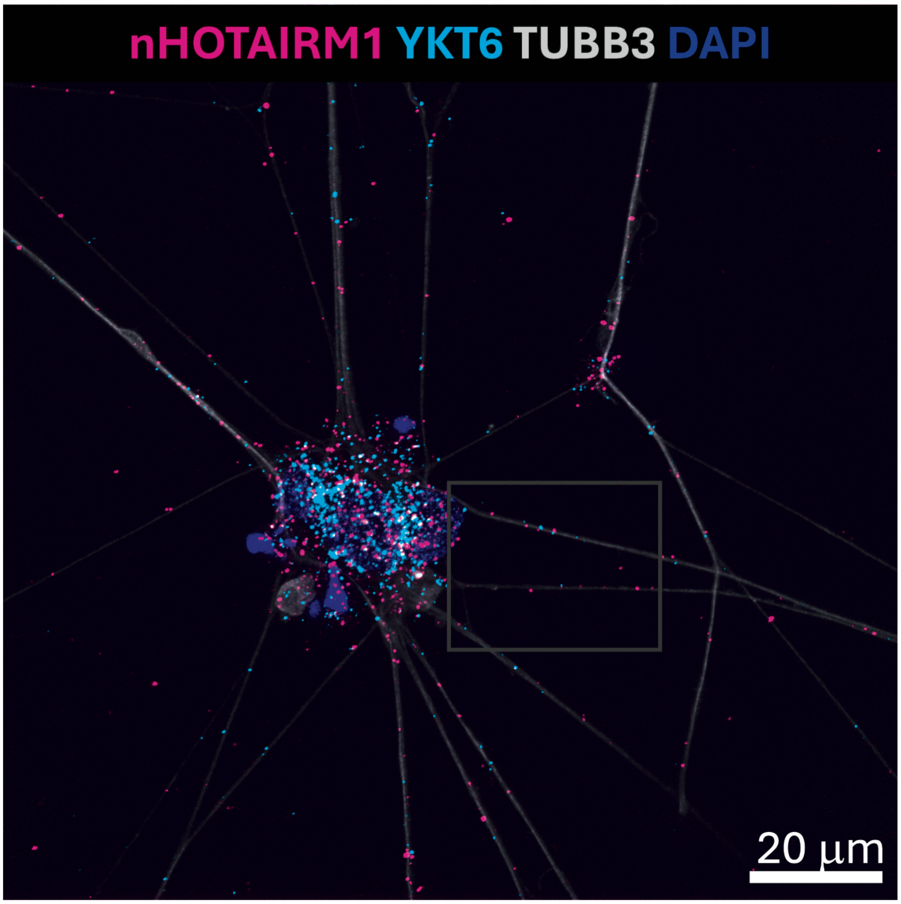


WT

**Supplementary Figure 9. Confocal imaging of *nHOTAIRM1* and *YKT6* mRNA in spMNs (related to main Fig. 5)**

Representative confocal Z-stack acquisition of spMN (A) soma and (B) neurites showing RNA-FISH for *nHOTAIRM1* (magenta) and *YKT6* mRNA (cyan), IF for TUBB3 (gray), and DAPI (blue). Images were acquired at 100x magnification using confocal microscopy with a Z-step size of 0.2 µm. The digital enlargements and 3D renderings shown in Figure 5 are derived from selected regions of these acquisitions. Scale bars: 20 µm.

Supplementary Figure 10

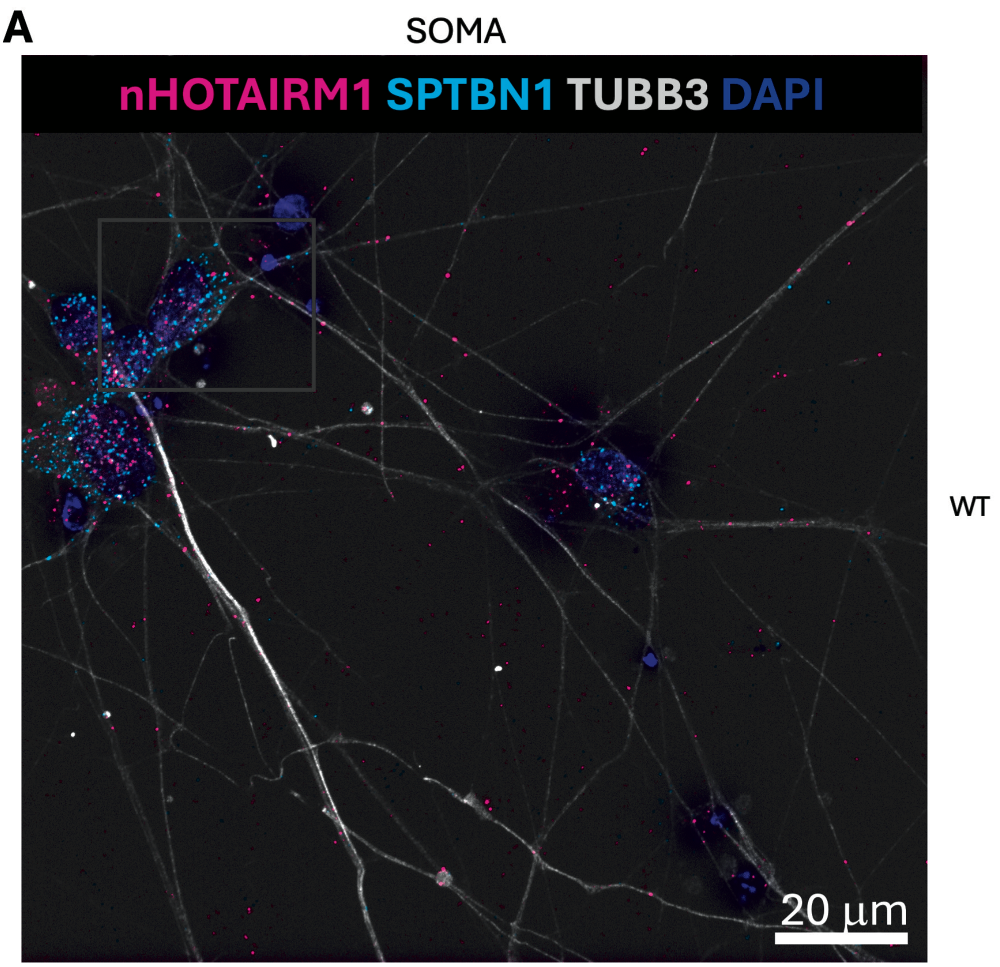


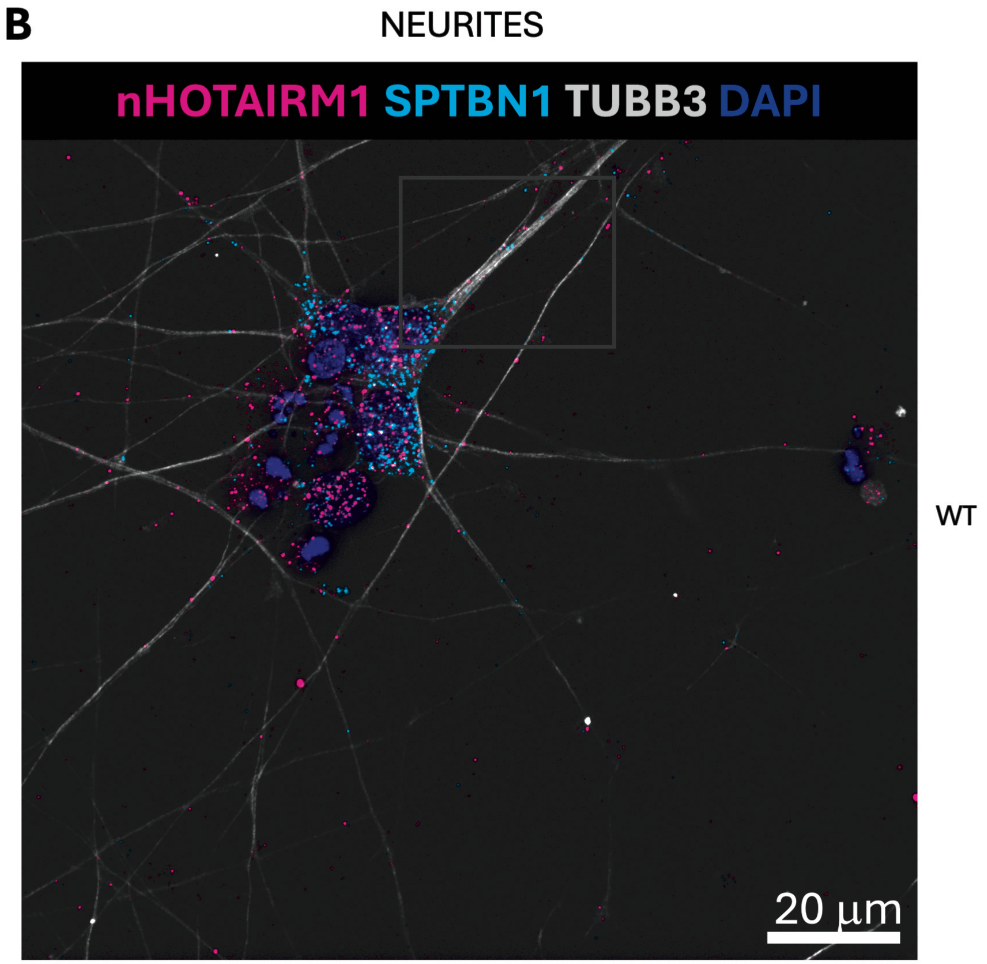

**Supplementary Figure 10. Confocal imaging of *nHOTAIRM1* and *SPTBN1* mRNA in spMNs (related to main Fig. 6)**

Representative confocal Z-stack acquisition of spMN (A) soma and (B) neurites showing RNA-FISH for *nHOTAIRM1* (magenta) and *SPTBN1* mRNA (cyan), IF for TUBB3 (gray), and DAPI (blue). Images were acquired at 100x magnification using confocal microscopy with a Z-step size of 0.2 μm. The digital enlargements and 3D renderings shown in Figure 6 are derived from selected regions of these acquisitions. Scale bars: 20 μm.

### Supplementary Figure 11

**A**

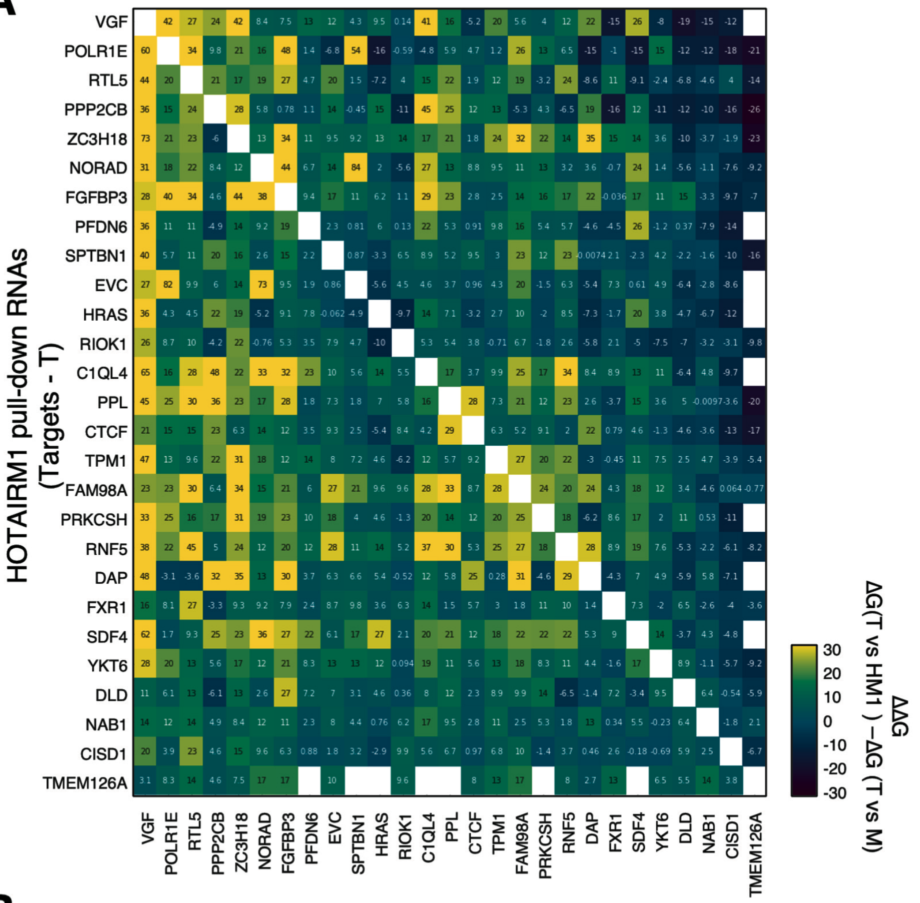


**B**

| Candidate mediator RNA | nHOTAIRM1 mRNA interactor(s) |
|---|---|
| ZC3H18 mRNA | TPM1 mRNA |
| POLR1E mRNA | YKT6 mRNA |
| RNF5 mRNA | SPTBN1 mRNA |
| FAM98A mRNA | TPM1 mRNA, YKT6 mRNA, SPTBN1 mRNA |

**C**

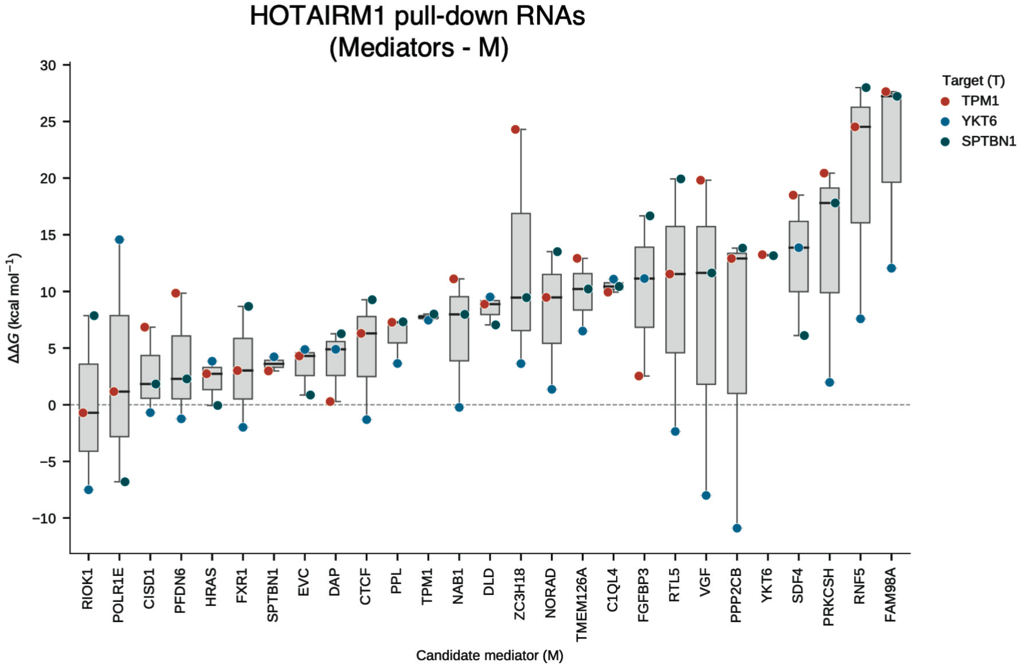

**Supplementary Figure 11. Computational identification of candidate mediator RNAs for *nHOTAIRM1*-mRNA interactions**

(A) Matrix of differential hybridization free energies (ΔΔG) for all ordered transcript pairs recovered in the *nHOTAIRM1* RNA pull-down. For each target transcript T and candidate mediator M, ΔΔG was calculated as ΔG(T versus *nHOTAIRM1*) − ΔG(T versus M), where ΔG(T versus *nHOTAIRM1*) represents the minimum free energy of the predicted direct duplex between *nHOTAIRM1* and the target, and ΔG(T versus M) represents the minimum free energy of the predicted duplex between the same target and the candidate mediator. Values were derived from interactions retained by the consensus prediction obtained using IntaRNA, RIsearch2 and RIME. Positive values indicate stronger predicted binding of the target to the candidate mediator than to *nHOTAIRM1*. White cells denote self-pairs or transcript pairs for which no consensus-supported duplex was retained. Values are expressed in kcal $mol^{-1}$.

(B) Summary table of candidate mediator RNAs selected for experimental validation. RNF5, ZC3H18 and POLR1E were identified as candidate mediators for the *nHOTAIRM1* interactions with *SPTBN1*, *TPM1* and *YKT6*, respectively, whereas FAM98A was identified as a shared candidate mediator for all three validated *nHOTAIRM1*-interacting mRNAs.

(C) Boxplots showing ΔΔG values between each candidate mediator and the three experimentally validated targets *TPM1*, *YKT6* and *SPTBN1*. Boxes indicate the median and interquartile range, and whiskers extend to the most extreme value within 1.5x the interquartile range. Candidate mediators are ordered according to increasing median ΔΔG. The dashed line indicates ΔΔG = 0.

### Supplementary Figure 12

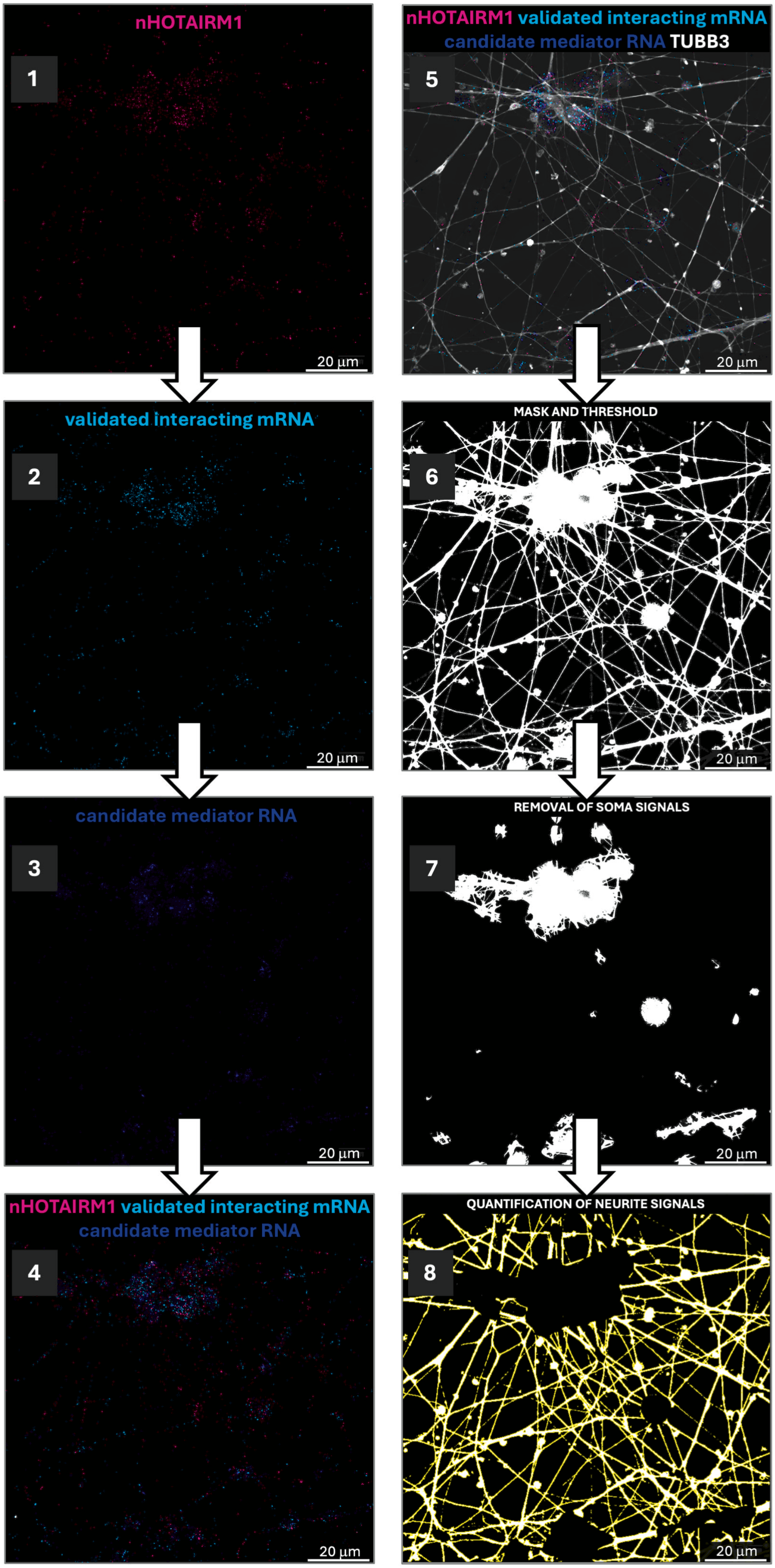

**Supplementary Figure 12. Image-analysis workflow for quantification of triple RNA colocalization events in neurites**

Representative workflow used for triple-target RNA-FISH analysis. Individual channels show *nHOTAIRM1* (1), a validated *nHOTAIRM1*-interacting mRNA (2) and a candidate mediator RNA (3). The three RNA signals were merged (4) and overlaid with TUBB3 IF to identify neuronal morphology (5). A binary neuronal mask was generated by thresholding the TUBB3 signal (6), and somatic regions were subsequently removed (7). The resulting neurite-restricted mask was used to quantify RNA signals and triple-colocalization events exclusively within neuronal projections (8). Scale bars: 20 μm.

### Supplementary Figure 13

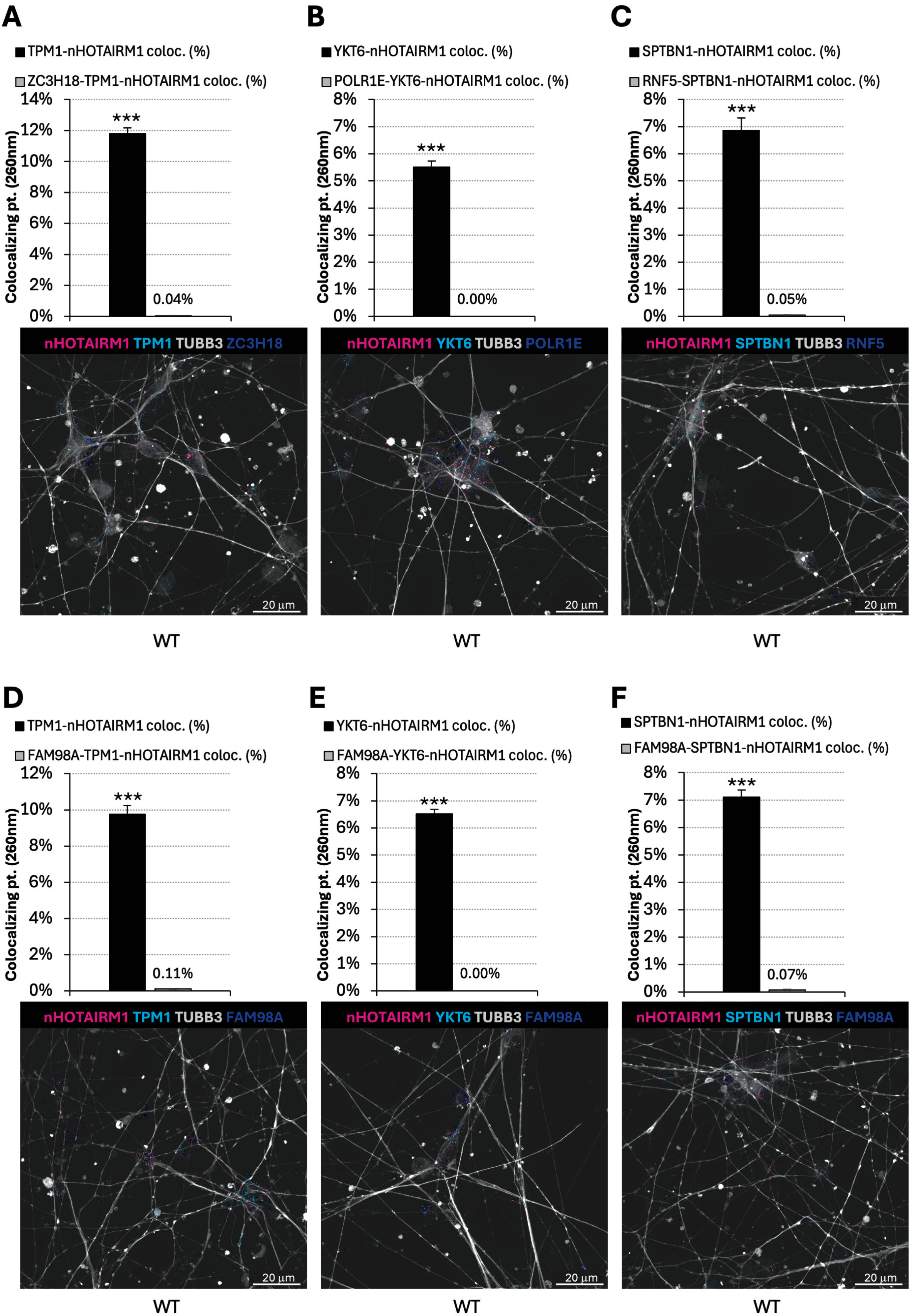

**Supplementary Figure 13. Triple-target RNA-FISH analysis of candidate mediator RNAs in neurites of iPSC-derived spMNs**

(A-C) Quantification and representative triple-target RNA-FISH images for the candidate mediators predicted for individual *nHOTAIRM1*-mRNA pairs: ZC3H18 with *nHOTAIRM1* and *TPM1* (A), POLR1E with *nHOTAIRM1* and *YKT6* (B), and RNF5 with *nHOTAIRM1* and *SPTBN1* (C).

(D-F) Quantification and representative triple-target RNA-FISH images for the shared candidate mediator FAM98A analyzed together with *nHOTAIRM1* and *TPM1* (D), *YKT6* (E), or *SPTBN1* (F).

Histograms show the percentage of *nHOTAIRM1*-mRNA colocalization events detected within neurites and the percentage of candidate mediator RNA puncta colocalizing with those *nHOTAIRM1*-mRNA events. Triple-colocalization events were defined using a centroid-to-centroid distance threshold of ≤260 nm. Representative images show *nHOTAIRM1* (magenta), the validated interacting mRNA (cyan), the candidate mediator RNA (blue), and TUBB3 (gray). Data are presented as mean ± SEM from three independent biological replicates. Scale bars: 20 μm. $^{*}P < 0.05$, $^{**}P < 0.01$, $^{***}P < 0.001$.

### Supplementary Figure 14

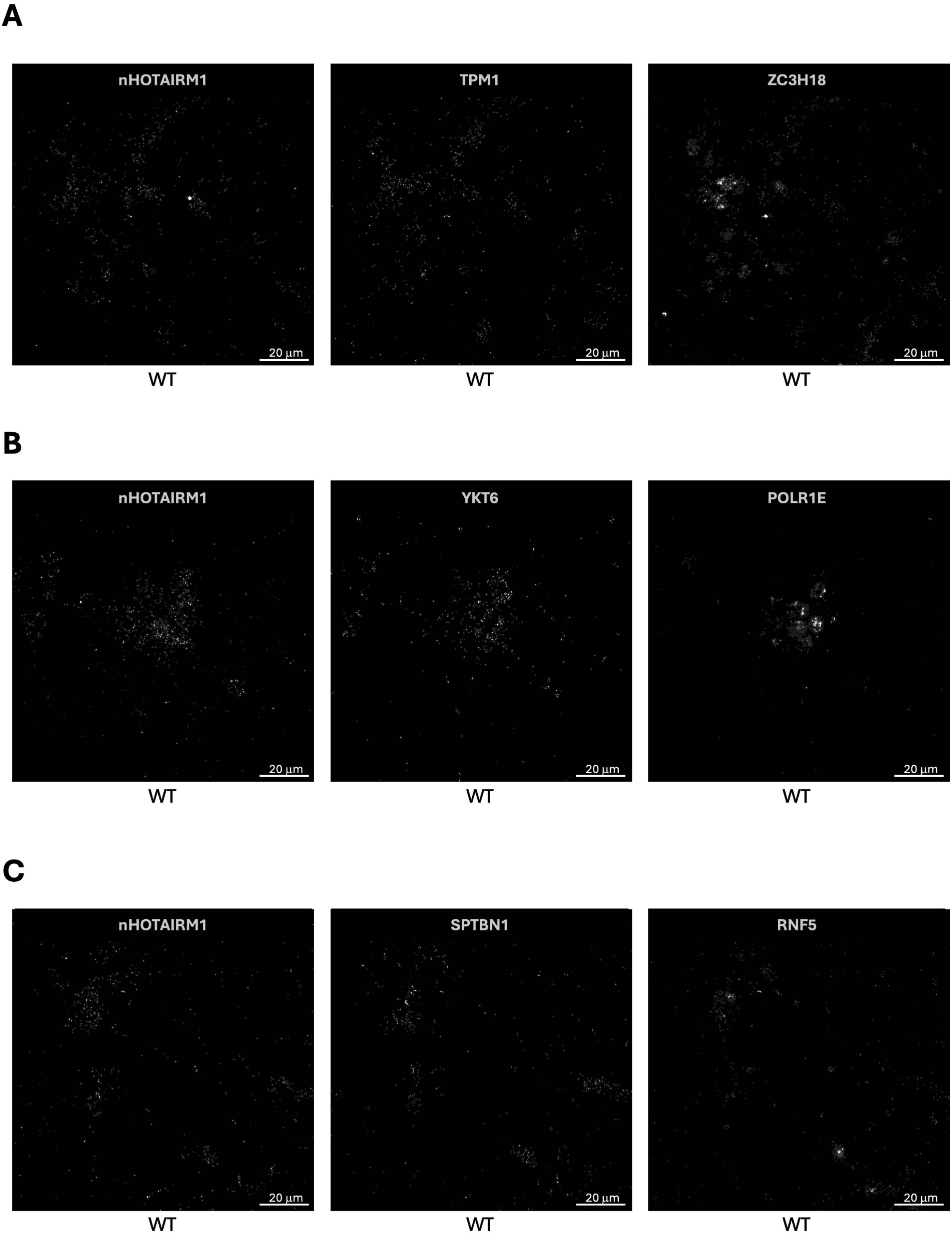

**Supplementary Figure 14. Binarized RNA-FISH channels for candidate mediators specific to individual *nHOTAIRM1*-mRNA pairs**

(A-C) Binarized single-channel images corresponding to the representative triple-target RNA-FISH acquisitions shown in Supplementary Figure 13A-C. Images show *nHOTAIRM1*, *TPM1* and ZC3H18; *nHOTAIRM1*, *YKT6* and POLR1E; and *nHOTAIRM1*, *SPTBN1* and RNF5, respectively. Binary signals were generated following background subtraction and thresholding and were used for neurite-restricted colocalization analysis. Scale bars: 20 µm.

### Supplementary Figure 15

**A**

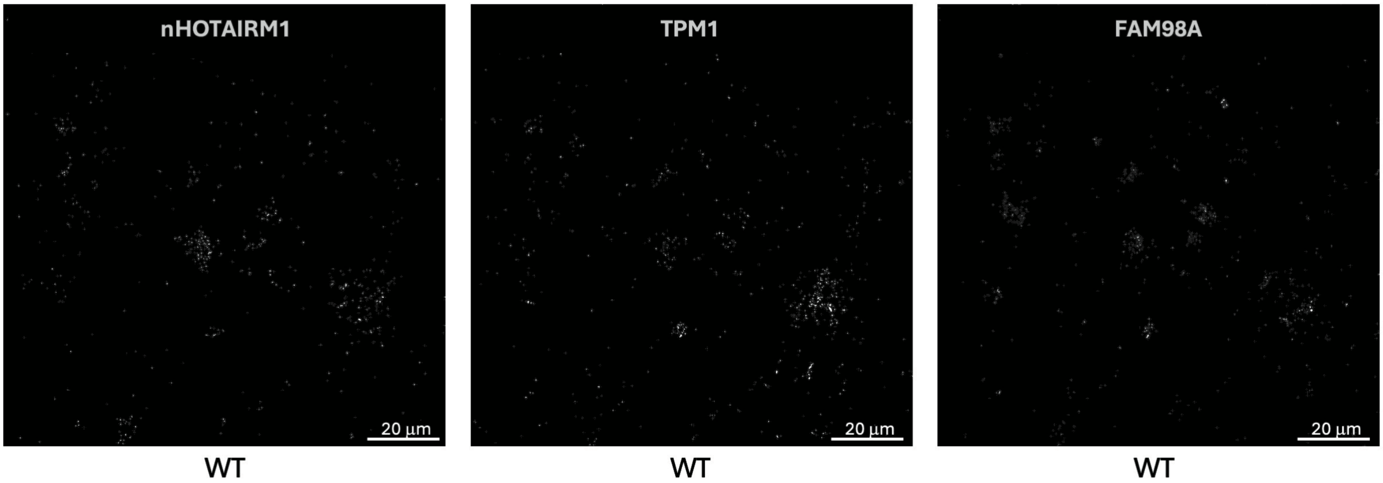


**B**

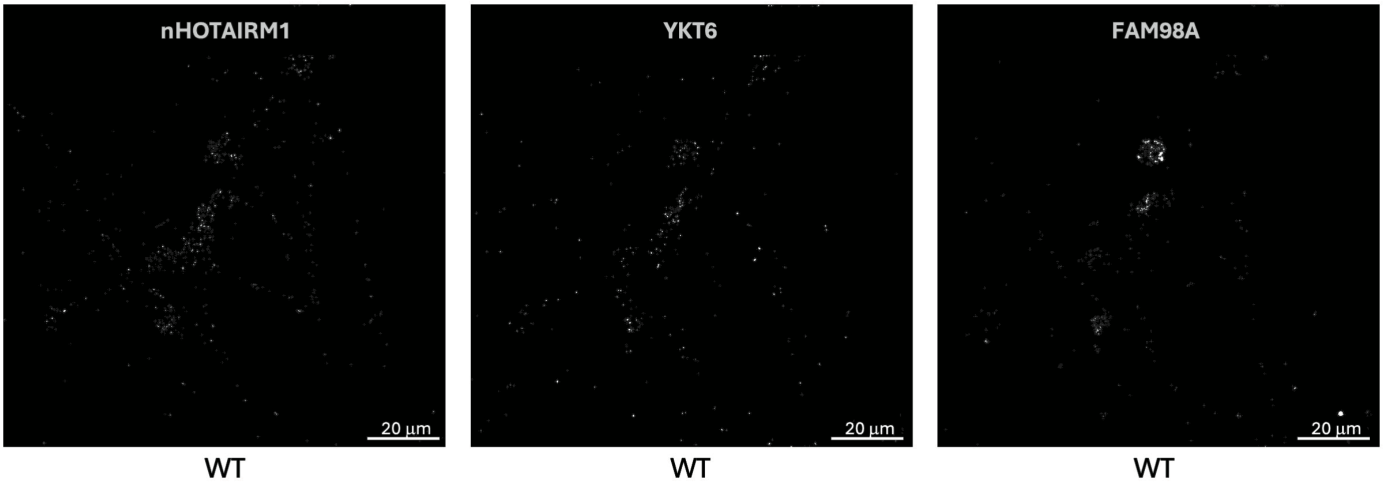


**C**

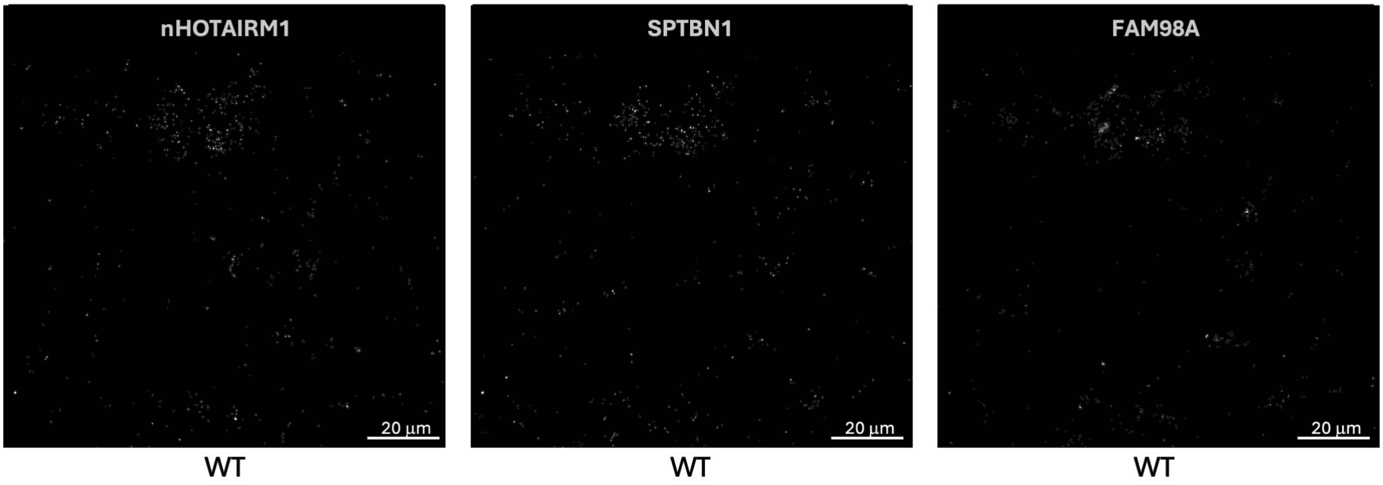

**Supplementary Figure 15. Binarized RNA-FISH channels for FAM98A-containing triple-target experiments**

(A-C) Binarized single-channel images corresponding to the representative triple-target RNA-FISH acquisitions shown in Supplementary Figure 13D-F. Images show *nHOTAIRM1*, *TPM1* and FAM98A; *nHOTAIRM1*, *YKT6* and FAM98A; and *nHOTAIRM1*, *SPTBN1* and FAM98A, respectively. Binary signals were generated following background subtraction and thresholding and were used for neurite-restricted colocalization analysis. Scale bars: 20 µm.

### Supplementary Figure 16

**A**

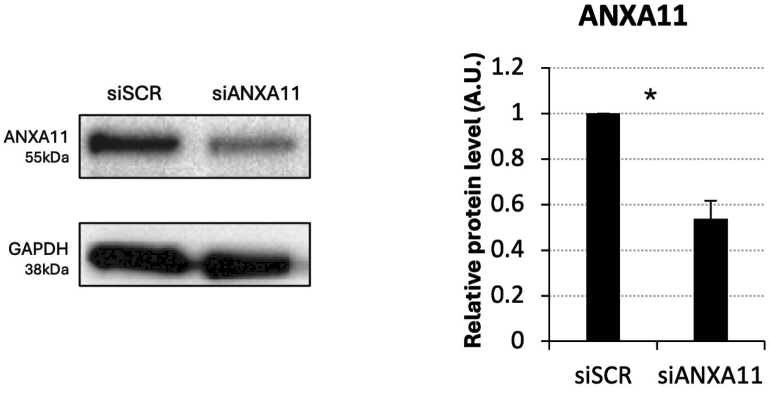


**B**

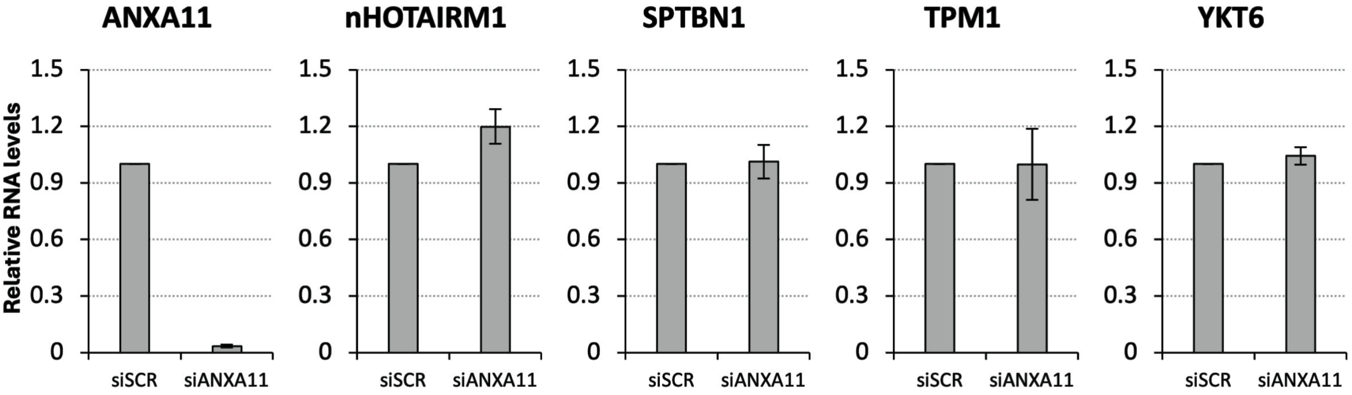

**Supplementary Figure 16. Validation of ANXA11 knockdown in iPSC-derived spMNs**

(A) Representative immunoblot analysis of ANXA11 and GAPDH in spMNs transfected with scramble siRNA (siSCR) or an equimolar mixture of two siRNAs targeting ANXA11 (siANXA11). The histogram shows ANXA11 protein levels normalized to GAPDH and expressed relative to the siSCR condition. The immunoblot is representative of three independent experiments and was cropped for clarity. Full-length uncropped blots are provided in Supplementary Data 1. Data are presented as mean ± SEM from three independent biological replicates (*N* = 2).

(B) qRT-PCR analysis of ANXA11, *nHOTAIRM1*, *SPTBN1*, *TPM1* and *YKT6* RNA levels in siSCR- and siANXA11-transfected spMNs. RNA levels were normalized to the corresponding scramble-control condition. Data are presented as mean ± SEM from two independent biological replicates (*N* = 2). *P < 0.05, **P < 0.01, ***P < 0.001.

Supplementary Figure 17

**A**

siSCR

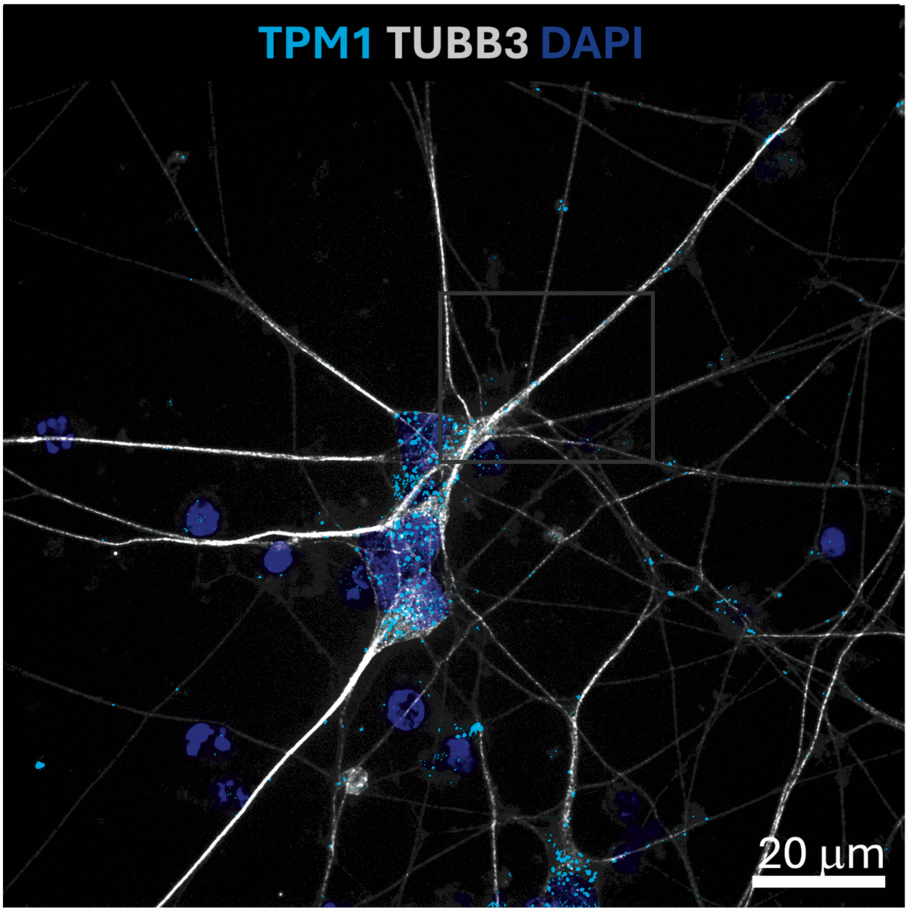


WT

**B**

siANXA11

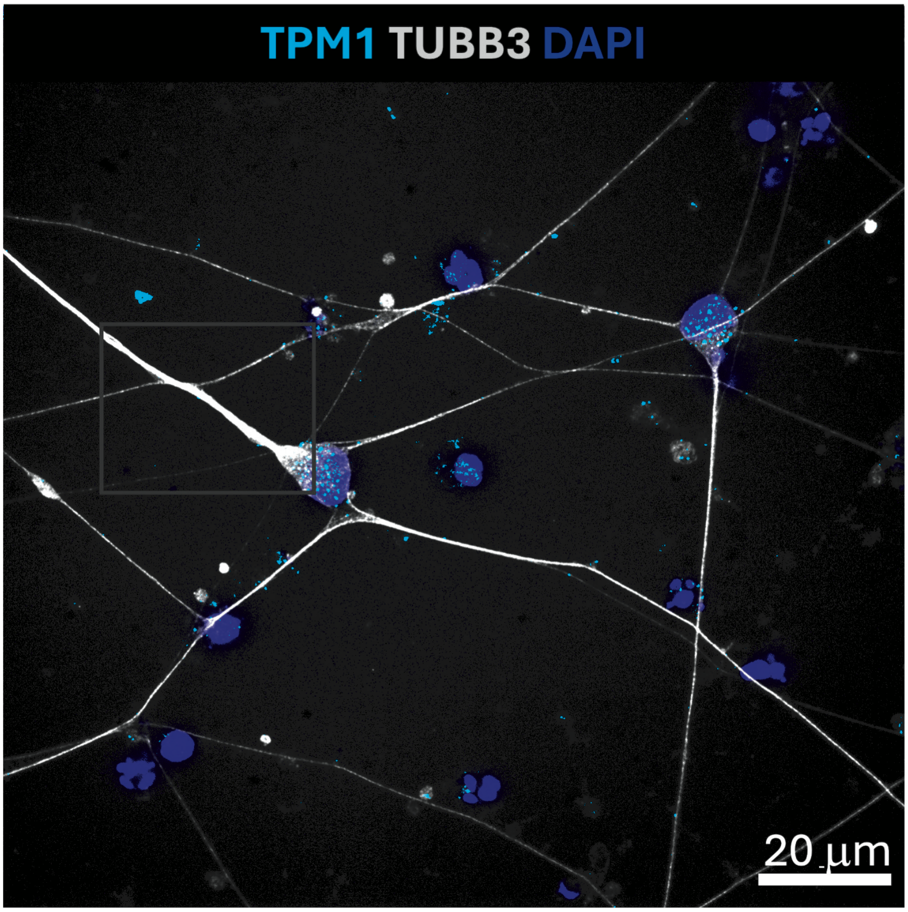


WT

**Supplementary Figure 17. Confocal imaging of *TPM1* mRNA following ANXA11 depletion related to Figure 8A**

Representative confocal Z-stack acquisitions of WT spMNs transfected with siSCR (A) or siANXA11 (B), showing RNA-FISH for *TPM1* mRNA (cyan), combined with IF for TUBB3 (gray) and DAPI nuclear staining (blue). Images were acquired at 100x magnification with a Z-step size of 0.2 µm. The representative neuritic regions shown in Figure 8A were selected from these acquisitions. Scale bars: 20 µm.

Supplementary Figure 18

A

siSCR

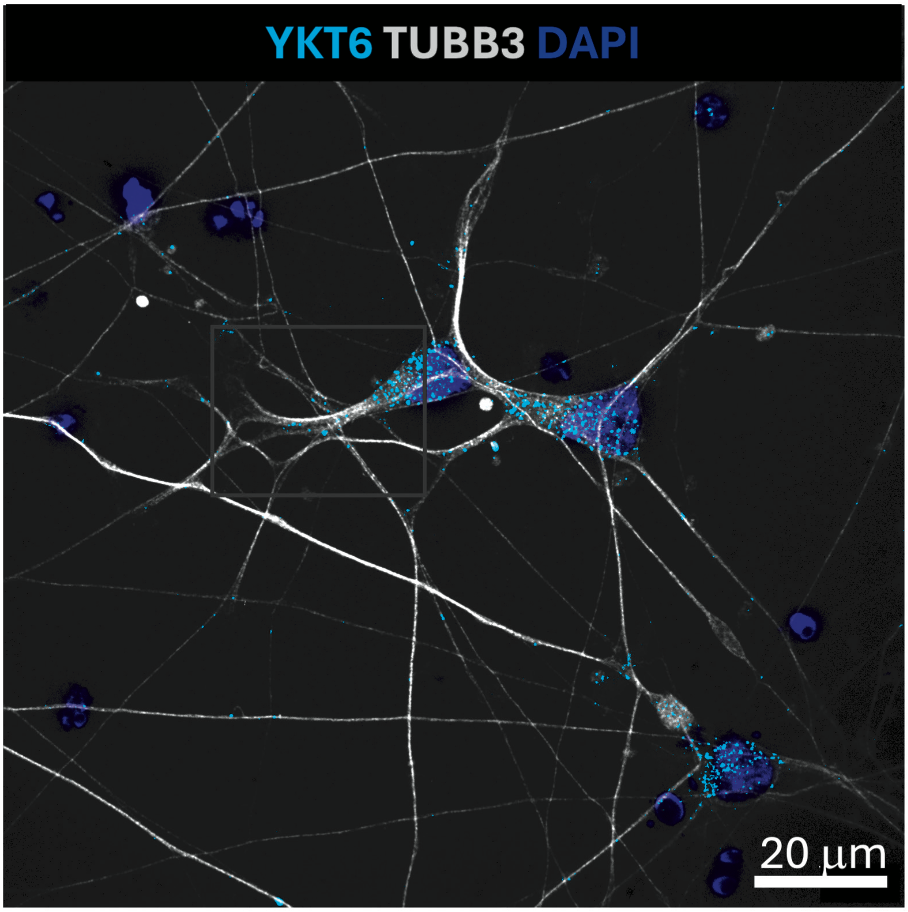


WT

B

siANXA11

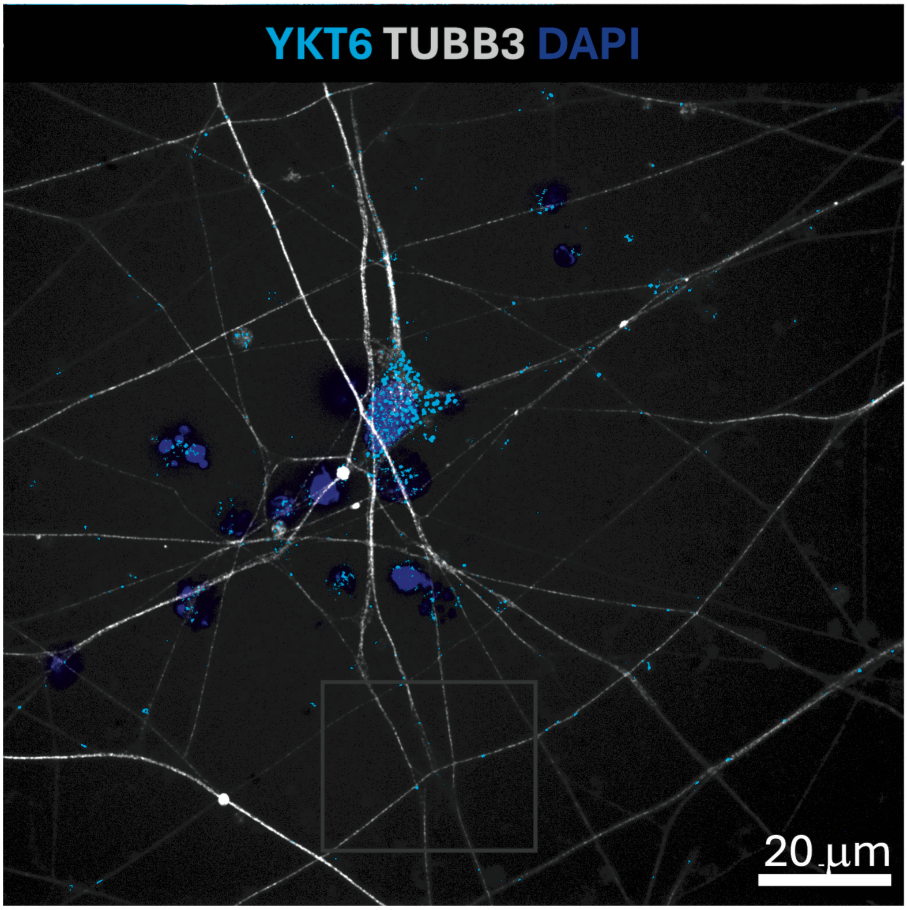


WT

**Supplementary Figure 18. Confocal imaging of *YKT6* mRNA following ANXA11 depletion related to Figure 8B**

Representative confocal Z-stack acquisitions of WT spMNs transfected with siSCR (A) or siANXA11 (B), showing RNA-FISH for *YKT6* mRNA (cyan), combined with IF for TUBB3 (gray) and DAPI nuclear staining (blue). Images were acquired at 100x magnification with a Z-step size of 0.2 μm. The representative neuritic regions shown in Figure 8B were selected from these acquisitions. Scale bars: 20 μm.

Supplementary Figure 19

**A** siSCR

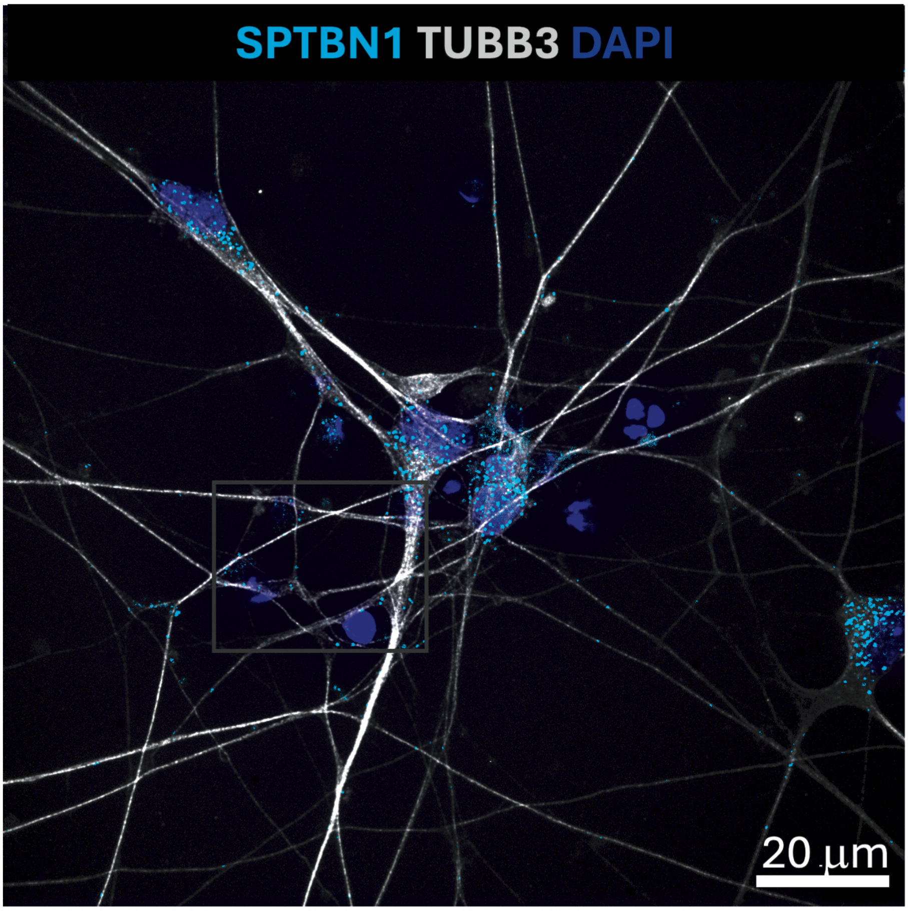


WT

**B** siANXA11

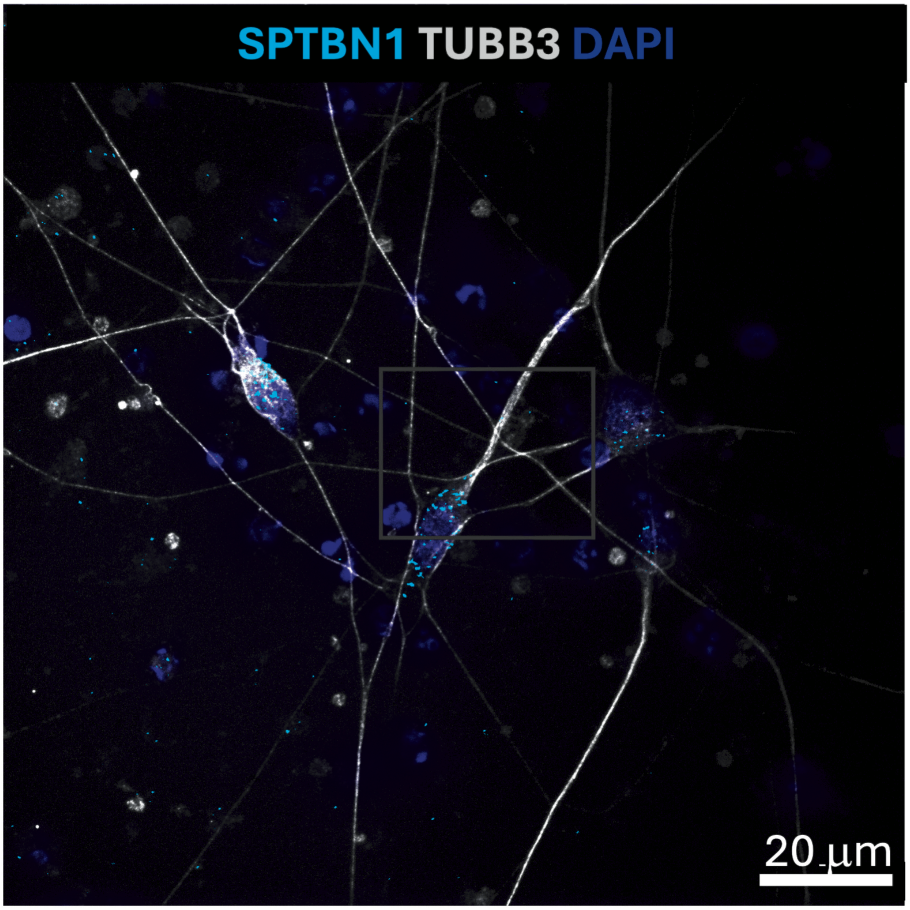


WT

**Supplementary Figure 19. Confocal imaging of *SPTBN1* mRNA following ANXA11 depletion related to Figure 8C**

Representative confocal Z-stack acquisitions of WT spMNs transfected with siSCR (A) or siANXA11 (B), showing RNA-FISH for *SPTBN1* mRNA (cyan), combined with IF for TUBB3 (gray) and DAPI nuclear staining (blue). Images were acquired at 100x magnification with a Z-step size of 0.2 μm. The representative neuritic regions shown in Figure 8C were selected from these acquisitions. Scale bars: 20 μm.

### Supplementary Figure 20

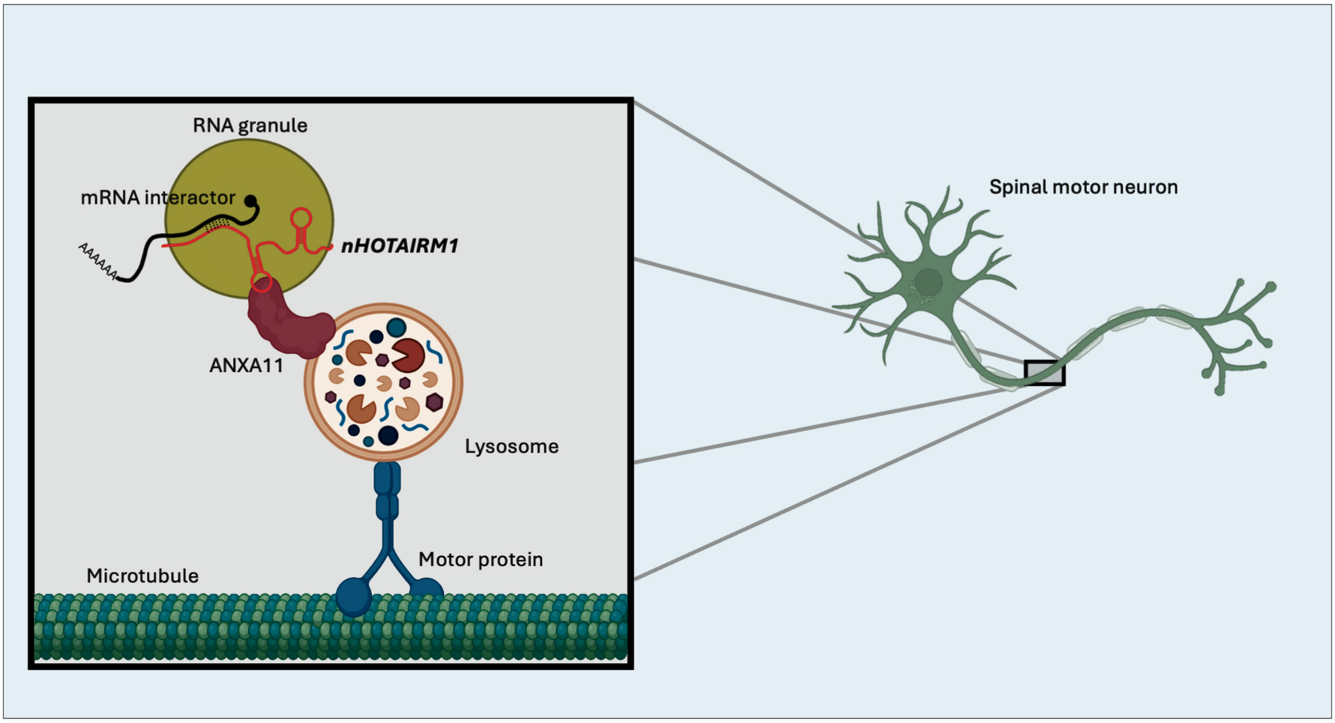

**Supplementary Figure 20. Illustrating model of *nHOTAIRM1*-mediated recruitment of target mRNAs into ANXA11-positive transport assemblies and their subsequent localization to distal neurites.**

**Supplementary Table S1. RNA-seq processing metrics and differential enrichment analyses of *nHOTAIRM1* native RNA pull-down**

The table reports RNA-seq processing metrics and differential enrichment analyses for *nHOTAIRM1* native RNA pull-down experiments performed in iPSC-derived spMNs. The “RNA-seq processing” sheet summarizes per-sample read counts across preprocessing, rRNA depletion, genome alignment and duplicate-removal steps. The “rRNA_ID” sheet lists the ribosomal RNA transcript and accession identifiers used for in silico rRNA read removal. The remaining sheets report edgeR differential analyses for EVEN versus INPUT and ODD versus INPUT comparisons, as well as pull-down specificity analyses comparing EVEN or ODD *nHOTAIRM1* enrichment relative to the LacZ control. Reported parameters include normalized expression values, log2 fold changes, likelihood-ratio statistics, P values and false discovery rates, together with gene annotations and associated analysis metrics where applicable.

**Supplementary Table S2. DNA probes, oligonucleotides and steric-blocking competitors used in this study**

The table reports the DNA probes and oligonucleotides used throughout the study. The “RNA Pulldown DNA Probes” sheet lists the sequences of antisense DNA probes used for *nHOTAIRM1* RNA pull-down assays. The “RNA FISH DNA Probes” sheet reports the HCR RNA-FISH probe sets used for detection of *nHOTAIRM1*, its interacting mRNAs and candidate mediator RNAs; probe sequences are available upon request. The “List of oligonucleotides” sheet reports primer sequences used for qRT-PCR and other oligonucleotide-based analyses. The “Competitor sequences” sheet lists the sequences of the non-targeting scramble control and the four DNA/LNA steric-blocking competitors designed to interfere with the predicted *nHOTAIRM1-mRNA* interaction regions.

**Supplemental Material** (PDF) reports the full-length uncropped original western blot images corresponding to the immunoblots presented in the main and supplementary figures. These images include all lanes and molecular weight markers prior to cropping for figure presentation.
